# Adaptive signal evolution and gene flow drive variable speciation outcomes in bark anoles

**DOI:** 10.64898/2026.11.15.760982

**Authors:** Tanner C. Myers, Pietro L. H. de Mello, Javier Torres, Paul M. Hime, Anthony J. Geneva, Richard E. Glor

## Abstract

New species evolve in a complex and non-linear fashion, resulting in situations where genomic and phenotypic variation between diverging lineages become decoupled. For example, natural selection acting on traits commonly used to diagnose species can instead drive trait variation to disagree with genomic evidence for species boundaries, challenging systematists aiming to delimit species. In this study, we combine phylogenetic, population genomic, and coalescent-based gene flow analyses to test whether convergence of a signaling trait has a shared genomic basis and to evaluate the evidence for unrecognized species level diversity in the *Anolis distichus* species group, a polytypic clade of lizards spanning the island of Hispaniola and the Bahamas in which diversity of a sex-linked signaling trait conflicts with genetic evidence for species boundaries. Comparing evolutionary histories of autosomes, sex chromosomes, and mitochondrial DNA, we find that the entire spectrum of signaling trait variation is present in four major geographic clades of the *A. distichus* species group and recover support for the independent evolution of this convergent signal. By quantifying isolation-by-distance and inferring migration across all contact zones on Hispaniola, we find evidence for lineages spanning the entire speciation continuum—from interbreeding populations to lineages exhibiting signatures of strong reproductive isolation. Together, our study highlights how phenotypic and genomic divergence may decouple during speciation, and provides a robust framework for gene flow-informed taxonomic revisions of the *A. distichus* species group.

## 1 Introduction

Speciation is a complex process involving genomic, phenotypic, and ecological divergence (Coyne and Orr, 2004; De Queiroz, 2007). As a consequence, phenotypic and genomic divergence are often uncoupled in diverging lineages: morphologically cryptic lineages may hide deep genomic splits (Bickford et al., 2007; Struck et al., 2018), while marked phenotypic differences can persist across porous genetic boundaries (Mundy et al., 2004; Lamichhaney et al., 2016). This decoupling has profound implications for species delimitation and phylogenetic inference: incomplete lineage sorting yields heterogeneous gene-tree histories and introgression generates reticulate networks that cannot be captured by strictly bifurcating models (Maddison, 1997; Leaché et al., 2014; Sólis-Lemus et al., 2016; Edwards et al., 2016). Comparing populations along the speciation continuum therefore provides a powerful framework to assess the association between phenotypic and genomic divergence and evaluate how introgression impacts the reconstruction of lineage relationships.

Sexual signaling traits provide a particularly useful lens on these processes. Within species, these traits mediate mate choice and influence competitive interactions (Endler, 1992; Price, 1998; While et al., 2015). Between species, they can contribute to species recognition and reproductive isolation, with reproductive character displacement accentuating signal differences at contact zones (Sætre et al., 1997; Boughman, 2001; Pfennig, 2016). Because sexual signaling traits are conspicuous, their differences have also been widely used for species delimitation (Padial et al., 2010), yet signal divergence need not reflect genome-wide lineage divergence (Harris et al., 2018). Instead, signals and the sensory systems that detect them may evolve in response to local environmental conditions, generating sharp geographic transitions in phenotype even among populations that continue to exchange genes (McLean et al., 2014; Poelstra et al., 2014). Such transitions can therefore mimic species boundaries while arising through processes operating within lineages (Zamudio et al., 2016; Cadena and Zapata, 2021). Although this complicates taxonomic inference, these situations offer a unique opportunity to identify the genetic basis and ecological drivers of phenotypic divergence during speciation.

### 1.1 Study System

The *Anolis distichus* species group consists of small lizards distributed across Hispaniola, the Bahamas, and surrounding islands. Notably among *Anolis*, these anoles exhibit striking inter-population variation in a sexually dimorphic, extensible throat fan (i.e., dewlap) used by males for visual communication (Nicholson et al., 2007; Losos, 2009). The group includes two major clades: the *A. brevirostris* and the *A. distichus* species complexes (Schwartz, 1968; Arnold, 1980; Hedges et al., 2019) (Fig. 1). The *Anolis brevirostris* complex consists of *A. brevirostris*, *A. caudalis*, *A. marron*, and *A. websteri* restricted to Haiti and the southwestern Dominican Republic. In this complex, dewlap color variation is largely consistent with reproductive character displacement at contact zones—*A. caudalis* has a white dewlap when co-occurring with *A. websteri*, but an orange dewlap elsewhere (Webster and Burns, 1973; Lambert et al., 2013). The *A. distichus* complex consists of the red-bodied *A. altavelensis*, endemic to the Hispaniolan satellite island of Alto Velo, and the polytypic *A. distichus* across Hispaniola and the Bahamas. The latter has traditionally been divided into 18 subspecies defined primarily by dewlap color and pattern, traits implicated in conspecific recognition (Schwartz, 1968; Williams and Rand, 1977; Case, 1990; Nicholson et al., 2007). In contrast to the *A. brevirostris* complex, dewlap color in *A. distichus sensu lato* does not reliably predict evolutionary relationships (Geneva et al., 2015; MacGuigan et al., 2017). Subspecies defined by a singular dewlap color can be polyphyletic, whereas markedly divergent color morphs can occur within a single polymorphic clade (Glor and Laport, 2012; Geneva et al., 2015; Ng et al., 2016; MacGuigan et al., 2017). Dewlap color in *A. distichus* instead tracks local light environments: orange and red dewlaps predominate in darker, mesic broadleaf forests, whereas yellow and white dewlaps are more common in bright, open xeric habitats (Ng et al., 2013b). Contact zones span a continuum from sharp ecological transitions with reduced gene flow, as between *A. d. ravitergum* and *A. d. ignigularis*, to gradual transitions with extensive introgression, as between adjacent dewlap morphs of the polymorphic *A. d. favillarum* (Case and Williams, 1984; Ng and Glor, 2011; Ng et al., 2016; MacGuigan et al., 2017; Geneva and Glor, 2025).

**Figure 1.**
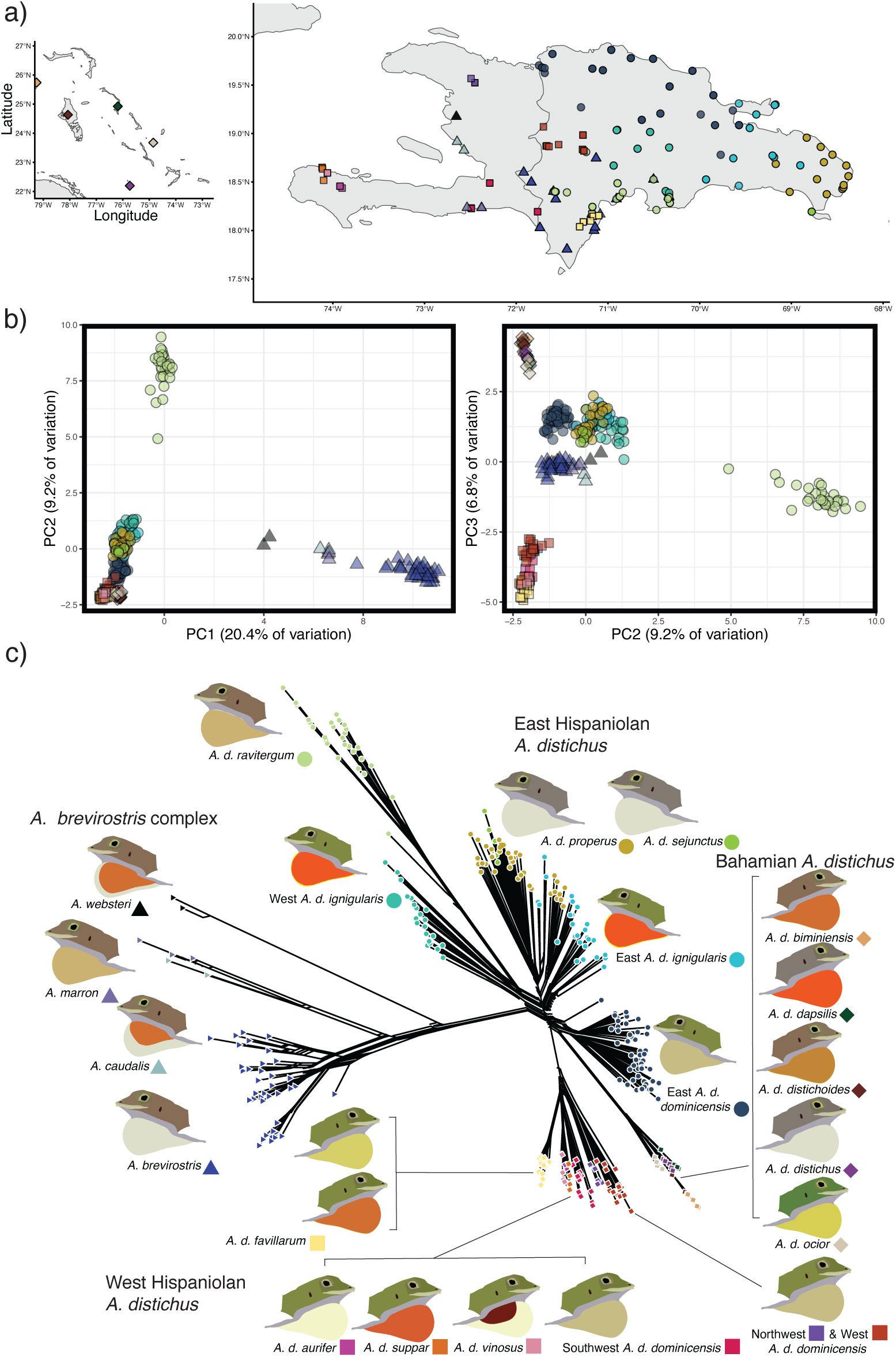
Sampling and Population Genomics of *Anolis distichus* species group. A. Sampling localities for the 297 anoles meeting filters for population genomic and phylogenetic analyses. Different colors distinguish the species and subspecies of *A. distichus* and shapes denote the “clade” to which specimens belong: Eastern *A. distichus* subspecies (circles), Bahamian *A. distichus* subspecies (diamonds), Western *A. distichus* subspecies (squares), and the *A. brevirostris* species complex (triangles). B. Scatterplots visualizing the first three principal components of the PCA performed on the 90% complete filtered SNP dataset. *Anolis distichus* and *A. brevirostris* complex-specific PCAs are visualized in Fig. S1-2. C. Neighbor-network estimated from the 90% complete filtered SNP dataset.

The striking phenotypic diversity of *A. distichus* is facilitated by a relatively simple genetic architecture. Dewlap coloration is highly heritable and influenced by a few loci of major effect (Ng et al., 2013a; Behere et al., 2024), with color variation associated with a small number of differentially expressed genes (de Mello et al., 2021). In the polymorphic subspecies *A. d. favillarum*, an orange-associated Y-chromosome haplotype accounts for a disproportionate share of color variation (de Mello et al., 2026). Together, the variability in reproductive isolation between populations and the association between dewlap color and environmental conditions make the *A. distichus* species group an ideal system to empirically assess the genetic basis of phenotypic divergence during speciation and to investigate how ongoing gene flow during divergence affects phylogenetic inference.

To tackle these systematic and evolutionary goals, we pursue three primary objectives: (1) infer phylogenetic relationships and patterns of gene flow among named *A. distichus* species group lineages to test whether traditional taxonomic designations track distinct evolutionary lineages or permeable phenotypic variants; (2) leverage a chromosome-level reference genome and the group’s neo-XXY sex-determination system (Falvey et al., 2026) to test whether recurrent orange–yellow dewlap polymorphisms is explained by a shared pair of Y-linked haplotypes segregating across island-wide contact zones (de Mello et al., 2026); and (3) evaluate the impact of incomplete lineage sorting versus reticulation on phylogenetic reconstruction of this clade, using phylogenetic species tree and coalescent-based migration rate analyses.

## 2 Material and Methods

### 2.1 Sampling and genomic library preparation

To obtain genome-scale markers for our phylogenetic and population genomic analyses, we sampled genomic DNA from 305 lizards across Hispaniola and the Bahamas. Our sampling includes representatives of 13 subspecies of *A. distichus*, *A. altavelensis*, and the four related species in the *A. brevirostris* complex: *A. brevirostris*, *A. caudalis*, *A. marron*, and *A. websteri*) (Fig. 1A; Fig. S1A; Fig. S2A; Table S1). We downloaded whole genome resequencing data of *A. pulchellus* for use as an outgroup (BioProject PRJNA949240; Wogan et al. (2023)). To represent the mitochondrial genome, we downloaded sequence data for the NADH dehydrogenase 2 (ND2) protein for 1,080 individuals from GenBank (Table S2, Fig. S3) using the “read.GenBank” function of the ape v.5.8-1 R package (Paradis and Schliep, 2019).

We prepared reduced-representation genomic libraries using a modified double-digest restriction site-associated DNA sequencing (ddRADseq) protocol (Peterson et al., 2012), following Myers et al. (2025). Genomic DNA was extracted from tail or liver tissue preserved in 95% ethanol at *−*80°C. Libraries were digested with SbfI and MspI, size-selected for fragments of 487-595 bp, and sequenced as 150-bp paired-end reads on an Illumina HiSeq X (Novogene Co.).

### 2.2 Bioinformatics

After sequencing, we assembled ddRADseq loci using the reference-based approach in Stacks v2.65 (Rochette and Catchen, 2017; Rochette et al., 2019). We first mapped reads with BWA MEM (Li, 2013) to the “AnoDis1.0” assembly of a male *A. d. ravitergum* (BioProject PRJNA1200026; BioSample SAMN45894292; Falvey et al. (2026)). We then sorted and indexed the resulting BAM files with samtools v1.15 (Danecek et al., 2021) prior to obtaining the catalog of ddRAD loci using the module gstacks. Finally, we used the populations module to output files for further filtering for downstream analyses.

For SNP-based analyses, we used bcftools v1.21 to filter the complete VCF including all samples (Danecek et al., 2021). Prior to filtering for high-quality calls, we excluded individuals with large amounts of missing data (*≥*85% of sites), eliminating 8 samples and leaving 297 for analysis. After excluding samples with excessive missingness, we applied depth, allele count, and additional missingness filters to retain high quality biallelic SNPs (O’Leary et al., 2018). For analyses assuming unlinked markers, we used PLINK v1.9 to keep only 1 SNP per ddRAD locus (Purcell et al., 2007). To minimize the impact of allelic dropout and maximize SNP yields, we applied these filters across all samples and for the *A. distichus* and *A. brevirostris* species complexes separately. Details for the SNP and whole locus datasets used for this study are found in Table S3.

### 2.3 Identifying evolutionary lineages

#### 2.3.1 Quantifying discrete population genetic structure

We performed principal components analysis (PCA) using the “glPca” function in adegenet v.2.1.11 (Jombart and Ahmed, 2011) on three datasets: all samples, *A. distichus* only, and the *A. brevirostris* species complex only. We visually inspected the first three principal components of each dataset. To estimate ancestry proportions and identify the number of ancestral populations (*K*), we used ADMIXTURE v1.3.0 (Alexander et al., 2009). Because its underlying model may be inappropriate for analyses across species boundaries (Pritchard et al., 2000), we analyzed *A. distichus* and *A. brevirostris* separately. For each complex, we tested *K* =1-10 with 10 independent runs per value and identified the optimal *K* based on cross-validation error.

We estimated pairwise *F_ST_* among species in the *A. distichus* species group and among *A. distichus* subspecies using StAMPP v1.6.3 (Weir and Cockerham, 1984; Pembleton et al., 2013). Each subspecies was treated as a population, except *A. d. dominicensis*, which previous studies suggest comprises four non-monophyletic geographic lineages (Geneva et al., 2015), and *A. d. ignigularis*, which our concatenated phylogenetic analysis resolved into eastern and western lineages (Fig. 3, S12). We estimated nucleotide diversity (*π*) using the Stacks v2.65 populations module and the complete ddRAD locus catalog to avoid biases introduced by filtering (Shafer et al., 2017).

We visualized genetic distance patterns across the *A. distichus* species group using the “Neighbor-Net” approach (Bryant and Moulton, 2004). We estimated a Nei’s *D* distance matrix (Nei, 1972) from the 90%-complete SNP dataset using StAMPP v1.6.3 and inferred the network with the “neighborNet” function in phangorn v2.12.1 (Pembleton et al., 2013; Schliep, 2011).

#### 2.3.2 Genomic differentiation and evidence for spatial isolation

Population clustering approaches such as ADMIXTURE can mistake isolation by distance (IBD) for discrete population structure, particularly with uneven geographic sampling (Wright, 1943; Bradburd et al., 2018). To evaluate whether genetic structure reflected IBD or spatial discontinuities consistent with restricted gene flow, we compared pairwise genetic and geographic distances. We estimated pairwise *F_ST_* between individuals using BEDASSLE v1.6 (Bradburd et al., 2013) and geographic distances using the “earth.dist” function in fossil v0.4.0 (Vavrek, 2011). We then assessed the association between linearized *F_ST_* and geographic distance using Pearson correlations in R.

#### 2.3.3 Evolutionary history of the nuclear and mitochondrial genomes

We inferred a maximum-likelihood phylogeny from concatenated ddRADseq loci using IQ-TREE2 v.2.4.0 (Minh et al., 2020). To recover complete loci for phylogenetic inference, we generated a whitelist of loci containing 90%-complete linkage-pruned SNPs and used the Stacks “populations” module to export the corresponding sequences in PHYLIP format. We used ModelFinder (Kalyaanamoorthy et al., 2017) to identify the optimal substitution model (TVM+F+I+R7) and evaluated node support using 1,000 ultrafast bootstrap and SH-aLRT replicates (Hoang et al., 2018; Shimodaira and Hasegawa, 1999; Guindon et al., 2010).

For mitochondrial phylogenetic inference, we aligned ND2 sequences using MAFFT v.7.475 (Katoh and Standley, 2013) and inferred a maximum-likelihood phylogeny with IQ-TREE2 v.2.4.0 (Minh et al., 2020). ModelFinder identified TN+F+I+R4 as the optimal substitution model (Kalyaanamoorthy et al., 2017), and node support was evaluated using 1,000 ultrafast bootstrap and SH-aLRT replicates.

### 2.4 Does the Y-chromosome explain dewlap color polymorphism?

To test whether Y-chromosome haplotypes explain orange/yellow polymorphisms across *A. distichus*, we reconstructed the evolutionary history of Y-linked markers and contrasted them with those of the autosomes and X chromosomes. We filtered SNPs from the relevant scaffolds of the AnoDis1.0 assembly corresponding to the Y-chromosome, the ancestral X chromosome (*X*_1_), and the neo-X (*X*_2_) (Falvey et al., 2026; de Mello et al., 2026). Considering only specimens identified *a priori* as males in the field, we dropped specimens from consideration that had *>* 80% missing data across the Y-linked scaffolds before excluding sites with depth less than 5 and depth greater than three times the third quartile (72), keeping biallelic SNPs that were at least 50% complete. Due to the excessive allelic dropout observed after filtering SNPs across the entire *A. distichus* species group, we only considered the subspecies of *A. distichus* for our sex chromosome analyses. To characterize sex chromosome population structure, we performed PCA on the three datasets of varying missingness and estimated pairwise *F_ST_* among populations using StAMPP in our 50% complete Y SNP dataset (Pembleton et al., 2013). We reconstructed four maximum-likelihood phylogenies of 182 male *A. distichus* using IQ-TREE2: (1) an autosomal phylogeny based on a 90% complete and linkage-filtered SNP whitelist, (2) a Y-chromosome phylogeny based on a 50% complete SNP whitelist, (3) an ancestral X-chromosome (*X*_1_) phylogeny based on a 50% complete SNP whitelist, and (4) a neo-X chromosome (*X*_2_) phylogeny based on a 50% complete SNP whitelist. To explore patterns of discordance among the autosomes and sex chromosomes, we quantified Robinson-Foulds distances between all tree pairs.

### 2.5 Disentangling effects of incomplete lineage sorting and gene flow

#### 2.5.1 Accounting for ILS using species tree approaches

We inferred species trees using two approaches: the polymorphism-aware phylogenetic model (PoMo) and SVDquartets under the multispecies coalescent. PoMo models population-level allele frequencies and accounts for ancestral and extant polymorphism (De Maio et al., 2015; Schrempf et al., 2016). We assigned individuals to populations according to current taxonomy, except where previous studies identified variation within subspecies (e.g., *A. d. dominicensis*). Using “FastaToCounts.py” (Schrempf et al., 2016), we generated per-population allele counts across 1,938,099 alignment sites and inferred a maximum-likelihood PoMo tree in IQ-TREE2 v2.4.0 under GTR+G4+P+N9 (general time-reversible substitution model, gamma rate heterogeneity, virtual population size of 9).

We also inferred a species tree from unlinked SNPs using SVDquartets in PAUP* v4.0a168 under the multispecies coalescent (Chifman and Kubatko, 2014, 2015). SVDquartets considers unlinked SNP data and uses singular value decomposition (SVD) to reconstruct the species tree topology under the multispecies coalescent model. We assessed support using 1,000 bootstrap replicates and summarized the resulting trees using SumTrees in DendroPy (Moreno et al., 2024). We treated well-supported monophyletic groups identified through ADMIXTURE as single terminals: the Bahamian subspecies, the subspecies from Haiti’s Tiburón Peninsula, and *A. d. dominicensis* populations west of the Cordillera Central.

#### 2.5.2 Quantifying gene flow across the speciation continuum

We used two approaches to characterize gene flow across the *A. distichus* species group. First, we used Dsuite (commit a547f99; Malinsky et al. (2020)) to detect excess allele sharing in the 90%-complete VCF containing all samples and *A. pulchellus* as the outgroup (Table S3). We ran “Dtrios” to estimate D-statistics and *f*_4_-ratios across all taxon trios, specifying the PoMo species-tree topology, and used “Fbranch” to summarize branch-specific excess allele sharing (Patterson et al., 2012; Malinsky et al., 2018). Results were visualized using dtools.py.

Second, we estimated gene flow between geographically adjacent populations using the multispecies coalescent with migration (MSC-M) implemented in BPP v4.7.0 (Flouri et al., 2018, 2023). Because the number of population pairs precluded simultaneous analysis, we analyzed each pair separately, allowing bidirectional migration, and evaluated support for each event *post hoc* (Ji et al., 2023). We used loci from the 90%-complete, linkage-pruned whitelist, retaining only loci present in both focal populations and all included individuals. To improve MCMC mixing, we downsampled populations to 8-12 individuals.

The MSC-M estimates the population migration rate *M* = *mN*, where *m* is the proportion of immigrants per generation and *N* is the effective size of the recipient population; *M* thus represents the expected number of immigrants per generation (Flouri et al., 2023). We assigned inverse-gamma priors IG(3, 0.002) to *θ* (mean 0.001) and IG(3, 0.008) to the root age *τ*_root_ (mean 0.004). Each analysis used a burn-in of 20,000 iterations, followed by 400,000 samples collected every two iterations, and was repeated four times with different random seeds. We assessed convergence using the potential scale reduction factor (PSRF) in CODA v0.19-4.1 (Plummer et al., 2006) and combined replicate posterior samples to estimate parameters and effective sample sizes.

We tested for gene flow using an approximate Savage-Dickey Bayes factor, defining a near-zero migration interval with *ɛ* = 0.01 and considering *BF*_10_ *>* 20 evidence for gene flow (Ji et al., 2023). Finally, we regressed estimated *M* against pairwise *F_ST_* using ordinary least-squares regression, predicting a negative relationship between migration and genetic differentiation.

## 3 Results

### 3.1 Identifying evolutionary lineages

#### 3.1.1 Quantifying discrete population genetic structure

Considering the entire *A. distichus* species group, the first principal component (20.4% of the cumulative variation) distinguishes *A. distichus* from members of the *A. brevirostris* species complex whereas the second principal components (9.2%) separates the subspecies *A. d. ravitergum* from the other *A. distichus* subspecies (Fig. 1B). Additionally, along the first principal component, other species of the *A. brevirostris* species complex (*A. caudalis*, *A. marron*, and *A. websteri*) cluster between *A. brevirostris* and *A. distichus sensu lato*. The third principal component (6.8%) distinguishes the Bahamian subspecies (*A. d. biminiensis*, *A. d. dapsilis*, *A. d. distichoides*, *A. d. distichus*, and *A. d. ocior*) and the subspecies found west of the Cordillera Central (western *A. d. dominicensis*, *A. d. favillarum*, *A. d. aurifer*, *A. d. suppar*, and *A. d. vinosus*) from those *A. distichus* populations found to the east (*A. d. ravitergum*, *A. d. ignigularis*, *A. d. properus*, and eastern *A. d. dominicensis*).

Considering only the *A. distichus* subspecies, principal component 1 (13.4%) primarily separates *A. d. ravitergum* from the other subspecies, with more subtle separation of the subspecies found southeast of the Cordillera Central and on Isla Saona (*A. d. ignigularis* (Fig. S1B). Principal component 2 (10.0%) distinguishes the Bahamian and western Hispaniolan *A. distichus* subspecies whereas principal component 3 (7.8%) primarily distinguishes populations found to the east and north of the Cordillera Central excluding *A. d. ravitergum* (Fig. S1C). Within the *A. brevirostris* species complex, principal component 1 (23.3%) distinguishes *A. brevirostris* from *Anolis caudalis*, *Anolis marron*, and *Anolis websteri* while principal component 2 (14.5%) separates *A. brevirostris* from the western Dominican Republic with clusters of consisting of populations adjacent to Lago Enriquillo and of populations from the Pedernales province (Fig. S2B). Principal component 3 (10.7%) primarily separates *A. websteri* from the other species (Fig. S2C).

ADMIXTURE recovered evidence for multiple populations within both *A. brevirostris* and *A. distichus*. For *A. brevirostris*, *K* =4 was characterized by the lowest cross-validation error (Fig. S4). However, clustering between *K* =3 and *K* =5 were similar and largely supported the subspecies of *A. brevirostris* (Fig. S5; Arnold (1980)). A population found in the western Dominican Republic (*A. b. brevirostris*) and a population found in Pedernales (*A. b. wetmorei*) were consistently recovered by the best supported run from *K* =3 to *K* =5. The differences were in whether the range of *A. b. deserticola* was split into a western and eastern populations and whether *A. brevirostris* from Barahona were considered admixed between the populations corresponding to *A. b. deserticola* and *A. b. wetmorei* or a distinct population.

The optimal value of *K* was less clear for *A. distichus*. While *K* =9 had the lowest cross-validation error in addition to reduced variance across runs, support for increasing values of *K* plateaus starting at *K* =6 (Fig. S6). Additionally, the populations identified across replicates varied even for the same value of *K* (ADMIXTURE results across all runs can be found here:). For *K* =6, the run with the lowest cross-validation error aligns closely to previous results inferred from a subset of these specimens (Myers et al., 2025), recovering populations corresponding to (1) Bahamian subspecies, (2) eastern *A. d. dominicensis*, (3) *A. d. ignigularis*, (4) *A. d. properus*, (5) *A. d. ravitergum*, and (6) *A. distichus* found west of the Cordillera Central (Fig. S7). The best supported *K* =9 run recovered the same populations as *K* =6, except the western population is split into Tiburón penninsula-endemic subspecies, *A. d. favillarum*, and western *A. d. dominicensis*, and western *A. d. ravitergum* was recovered as a population separate from eastern *A. d. ravitergum*. With increasing values of *K*, the additional populations identified by ADMIXTURE lack consistency across *K*. For example, while the best supported run for *K* =7 recovers *A. d. favillarum* as a population, the best supported *K* =8 scenario lumps *A. d. favillarum* with the subspecies of the Tiburón penninsula.

Pairwise comparisons of relative differentiation (*F_ST_*) ranged widely from 0.06 to 0.9 among the populations considered, spanning the population-species continuum (Fig. S8). Greater *F_ST_* appears to mirror increasing geographic isolation, with the Bahamian populations and other members of the *A. brevirostris* complex being involved in some of highest differentiation comparisons, although variation in sample sizes among populations may inflate these estimates. Nucleotide diversity (*π*) ranged from 0.00087 to 0.0047 (Fig. S8), exhibiting similar patterns to *F_ST_* with the lowest estimates corresponding to the Bahamian populations of *A. distichus* and highest estimates generally corresponding to the connected populations of *A. distichus* found east of the Cordillera Central, although the Isla Saona-endemic subspecies, *A. d. sejunctus*, exhibited the greatest diversity of any population. Additionally, members of the *A. brevirostris* complex were less diverse than the *A. distichus* complex.

The unrooted phylogenetic network based on Nei’s D recovered nine distinct clusters corresponding to the four species of the *A. brevirostris* complex, the Bahamian *A. distichus* subspecies, the *A. distichus* populations found west of the Cordillera Central, *A. d. ravitergum*, the eastern population of *A. d. dominicensis*, and the subspecies *A. d. ignigularis*, *A. d. properus*, and *A. d. sejunctus* (Fig. 1C). The longest internal branch is the one leading to the *A. brevirostris* complex, whose members are also separated by relatively long branches. Branches within the *A. distichus* complex are shorter between geographically connected populations (e.g., among western *A. distichus* subspecies), with the exception of *A. d. ravitergum*. In addition to *A. d. dominicensis*, *A. d. ignigularis* is also rendered non-monophyletic with individuals from the western end of its range clustering with *A. d. ravitergum* and those found to the east clustering among *A. d. properus*, which we hypothesize is due to gene flow where these populations come into contact.

#### 3.1.2 Genomic differentiation and evidence for spatial isolation

To assess whether evidence for genetically distinct populations might be due to isolation-by-distance, we compared genetic distances with geographic distances estimated within and between geographically adjacent populations. These results corroborate that *A. brevirostris* and *A. distichus* represent distinct species separated by strong reproductive isolation (Fig. 2, S9). The IBD relationships between *A. brevirostris* and the *A. distichus* subspecies *A. d. ravitergum* and *A. d. favillarum*, which overlap in the southwestern Dominican Republic and Sierra de Baoruco respectively, are discontinuous. Within *A. distichus*, the most discontinuous IBD relationships involve *A. d. ravitergum* (Fig. 2, S10-11). Additionally, eastern and western populations of *A. d. dominicensis* exhibit a mostly discontinous IBD pattern, supporting the role played by the Cordillera Central as a biogeographic barrier, although these populations may come into contact near the border of Haiti and the Dominican Republic in northern Hispaniola (Fig. S10). All other comparisons explored between the subspecies of *A. brevirostris* and among populations of *A. distichus* exhibited partial to complete overlap, suggesting IBD explains the recovered evidence for population strucuturing between them.

**Figure 2.**
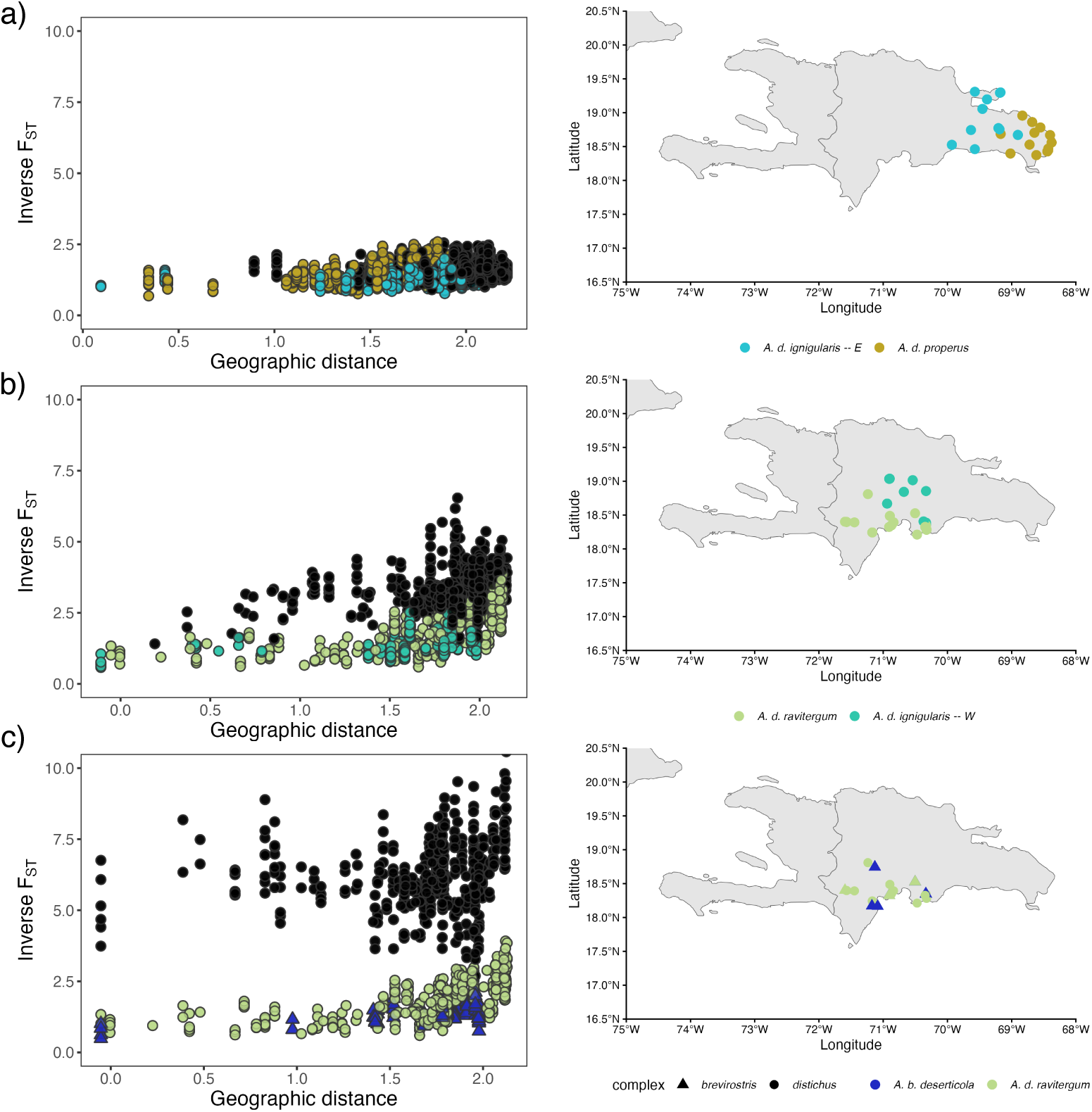
IBD relationships illustrate speciation continuum. Comparisons ranging from (a) populations that vary in dewlap color but appear to hybridize frequently (*A. d. properus* and *A. d. ignigularis*), to (b) those farther along the speciation continuum that exhibit some degree of reproductive isolation (*A. d. ravitergum* and *A. d. ignigularis*), and, finally, (c) recognized species (*A. b. deserticola* and *A. d. ravitergum*) recover increasing genetic discontinuity with greater reproductive isolation.

#### 3.1.3 Evolutionary history of the nuclear and mitochondrial genomes

The maximum likelihood reconstruction of relationships from concatenated ddRAD loci was well-resolved and largely mirrored the clustering observed in our phylogenetic network inferred from genetic distances (Fig. S12). Members of the *A. brevirostris* complex formed a clade, with *A. websteri* placed as sister to *A. brevirostris* and a clade of *A. caudalis* and *A. marron*. The subspecies *A. d. ravitergum* is sister to the other subspecies of *A. distichus*. Within that ingroup, the subspecies *A. d. ignigularis*, *A. d. properus*, and *A. d. sejunctus* found east of the Cordillera Central form a clade opposite the eastern population of *A. d. dominicensis* and the Bahamaian subspecies and the populations found west of the Cordillera Central. The only geographic populations identified by our population genomic analyses to be recovered as non-monophyletic were *A. d. ignigularis* and *A. d. properus*.

The ND2 gene tree topology mirrors those recovered by previous studies (Fig. S13, S15– 22) (Glor and Laport, 2012; Ng et al., 2016), but differs from the concatenated ddRAD tree at three main places (Fig. S14). First, rather than being sister to the rest of *A. distichus* as suggested by the nuclear genome phylogeny, the subspecies *A. d. ravitergum* is nested within a clade of *A. d. ignigularis*. Second, rather than being nested within the subspecies *A. d. properus*, the Isla Saona-endemic subspecies *A. d. sejunctus* was recovered as sister to a clade consisting of *A. d. ignigularis*, *A. d. properus*, and *A. d. ravitergum*. Third, eastern populations of *A. d. dominicensis* were recovered as sister to the rest of *A. distichus*. In addition to the discordant placement of subspecies, we observed one individual shared by the ddRAD and mtDNA datasets that possessed discordant nuclear and mitochondrial ancestry between the species complexes. Identified in the field as *A. brevirostris*, GLOR 4320 clustered among *A. brevirostris* in the concatenated ddRAD tree and among *A. d. ignigularis* in the ND2 tree (Fig. S14, S20).

### 3.2 Does the Y-chromosome explain dewlap color polymorphism?

The Y-chromosome is characterized by more elevated *F_ST_* than the genome-wide average (Fig. S8, S23), even as population clustering largely mirrors genome-wide patterns (Fig S24). Relationships inferred from Y-chromosome loci more closely resembled the phylogeny inferred from autosomal loci than expected if shared Y-chromosome haplotypes explained orange/yellow polymorphisms across *A. distichus* (Fig. 3, S25–26). Rather than forming distinct clades of orange- and yellow-dewlapped populations, the Y-chromosome phylogeny largely mirrors geographic patterns of divergence, grouping together populations that vary dramatically in dewlap coloration such as the Bahamas and Tiburon penninsula subspecies. The Y-chromosome phylogeny also clusters *A. distichus* subspecies with greater accuracy than the autosomal tree. For example, disagreements between the two trees include the Y phylogeny placing the Southwestern *A. d. dominicensis* population with other Western *A. d. dominicensis* populations rather than with the other Tiburon peninsula subspecies, and placing Western *A. d. ignigularis* sister to Eastern *A. d. ignigularis*. In addition, the Bahamas subspecies renders Eastern *A. d. dominicensis* paraphyletic, and *A. d. properus* partially clusters with *A. d. ignigularis*.

**Figure 3.**
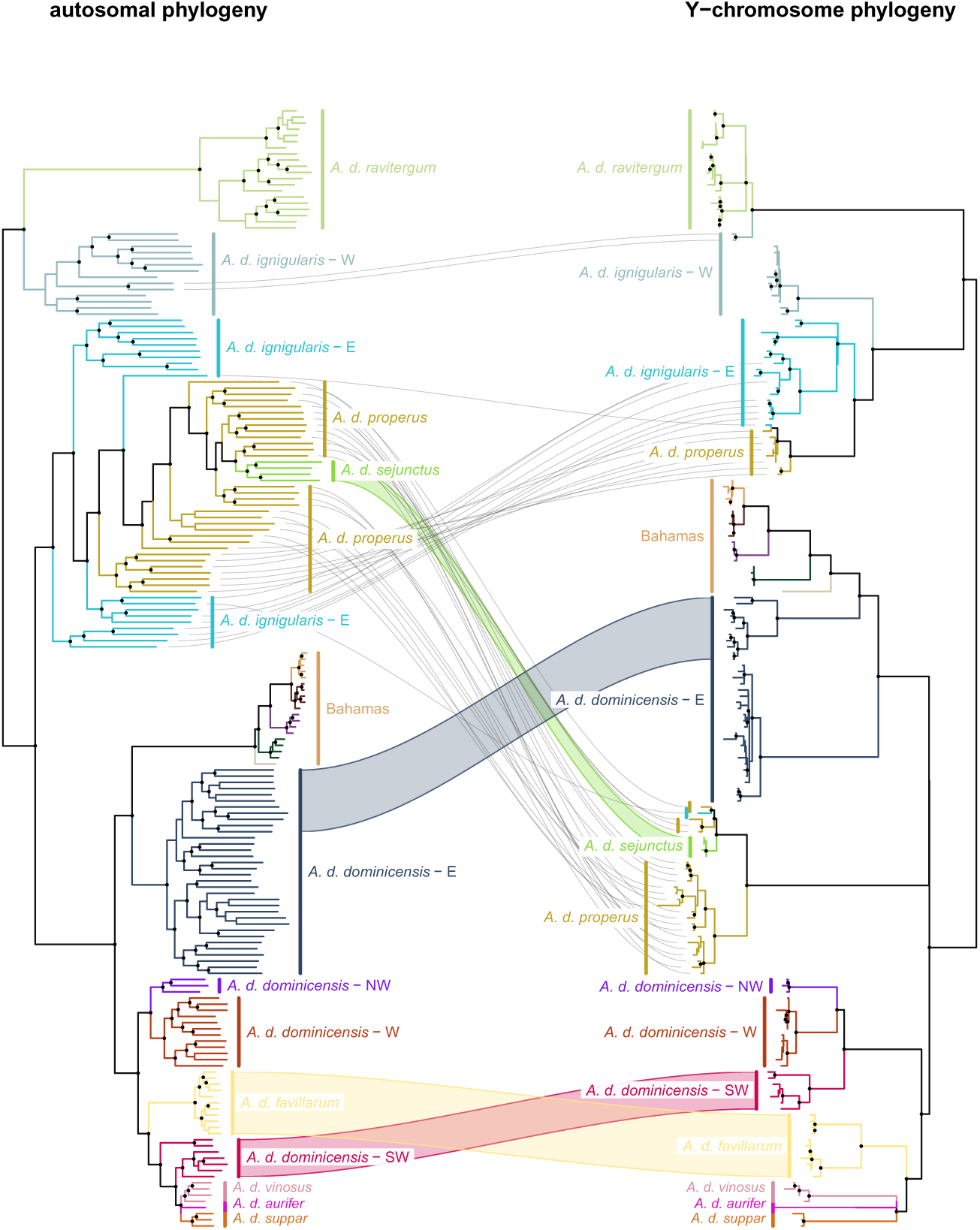
Y-chromosome relationships do not recapitulate dewlap color patterns. The evolutionary history of the Y-chromosome largely mirrors autosomal relationships and therefore, geography, instead of dewlap color pattern differences. Cases of Y-autosome discordance is largely restricted to geographically proximate lineages where introgression may have occurred. Predictably, the Y-chromosome, characterized by strongly reduced recombination and degradation, appears less permissive to gene flow than autosomes.

Our topological comparisons across genomic partitions confirm that the ancestral X (*X*_1_) tree is the topologically closest to the autosomes out of the sex chromosomes (normalized Robinson-Foulds distance nRF = 0.592, sharing 73 of 179 internal splits). The ancestral X (*X*_1_) and neo-X (*X*_2_) phylogenies largely agree with each other and mirror the autosomal organismal backbone (Fig. S26–29). Although raw tip-level comparisons show elevated discordance between *X*_1_ and *X*_2_ (nRF = 0.749), collapsed-backbone analysis reveals high deep-level concordance (backbone nRF = 0.235, sharing 13 of 17 splits), supporting that localized tip reticulation between Eastern *A. d. ignigularis* and *A. d. properus* predominantly drives sex chromosome discordance (Fig. S29).

### 3.3 Disentangling effects of incomplete lineage sorting and gene flow

#### 3.3.1 Accounting for ILS using species tree approaches

The two species tree approaches largely agreed with the concatenated phylogeny. The primary difference between the SVDquartets tree and the concatenated tree was the lack of support (55% of bootstrap replicates) for the clade of eastern *A. d. dominicensis* and the Bahamian subspecies of *A. distichus*, and the placement of that clade as sister to the populations found west of the Cordillera Central (*<*50%; Fig. S12, S31). The PoMo tree exhibited toplogical differences from the concatenated tree, namely by *A. d. ignigularis* being rendered non-monophyletic by the western population being found sister to the rest of *A. distichus* except *A. d. ravitergum*, and by the Bahamian subspecies being sister to the western populations of *A. distichus* instead of eastern *A. d. dominicensis* (Fig. S32).

#### 3.3.2 Quantifying gene flow across the speciation continuum

Fbranch tests recovered 17 significant (FDR-adjusted *p*-value *<* 0.05) cases of excess allele sharing between members of the *A. distichus* species group (Fig. S33; Table S7). The strongest signals involved eastern *A. d. ignigularis* and the neighboring *A. d. properus* (fb = 0.494, Z-score = 5.42, *p*=1.14 *×* 10*^−^*^7^), eastern *A. d. dominicensis* (fb = 0.499, Z-score = 5.42, *p* = 1.14 *×* 10*^−^*^7^), and western *A. d. ignigularis* (fb = 3.89, Z-score = 3.76, *p* = 0.00019553). Beyond these cases where excess allele sharing is associated with contemporary range overlap and likely gene flow, several significant cases involve geographically disjunct populations separated by a population with a range that overlaps with both (e.g., eastern *A. d. dominicensis* and *A. d. properus*: fb = 0.236, Z-score = 4.54, *p* = 9.58 *×* 10*^−^*^6^). Additionally, we recover evidence for interspecific excess allele sharing between geographically overlapping populations of *A. brevirostris* and *A. d. ravitergum* (fb = 0.092, 5.67, *p* = 3.98 *×* 10*^−^*^8^). The other significant results involve geographically disparate and often phylogenetically distantly related taxa, such as between *A. websteri* and eastern *A. d. ignigularis* (fb = 0.200, Z-score = 5.42, *p* = 1.14 *×* 10*^−^*^7^).

Convergence diagnostics for 8 comparisons of geographically adjacent taxa suggested that convergence was reached across all replicates with potential scale reduction factor (PSRF) values close to 1.0 and (Table S8; Fig. S34-40). Another comparison (Tiburon *A. distichus* subspecies and west *A. d. dominicensis*) appeared to have reached convergence in three of the four chains; we chose to present results from these replicates due to the possibility that the fourth chain identified a local optimum (Fig. 41). The other three comparisons tested here (*A. d. ignigularis* and *A. d. properus*; east and west *A. d. ignigularis*; east and west *A. d. dominicensis* (Fig. S42-44)) disagreed across replicates for the two migration rate parameters while the overall likelihood of the model appeared stationary. For these chains, each direction included two chains corresponding to “high” migration and two corresponding to “low” migration suggesting possible “label-switching” took place, a known issue for gene flow tests using the multispecies coalescent (Yang and Flouri, 2022). While these comparisons could not reliably indicate which direction of gene flow was more frequent, we pooled the “high” and “low” migration chains together from the two directions and reported those statistics.

Based on Bayes factors, every direction tested except for *A. d. favillarum* into *A. b. wetmorei* was recovered as significant (BF_10_ *>* 20). Migration rates between *A. brevirostris* and *A. distichus* are generally lower than those estimated from comparisons within the *A. brevirostris* complex and *A. distichus* complex, although certain directions of migration between some of these pairs (e.g., *A. d. ravitergum* to *A. d. ignigularis*) are comparable to between species migration rates (Fig. 4). The highest rates of migration involve *A. d. ignigularis* and the neighboring eastern *A. d. dominicensis* and *A. d. properus*, which approximate or exceed even the level of migration inferred between western and eastern populations of *A. d. ignigularis*. Migration rates within species complexes (i.e., not between *A. brevirostris* and *A. distichus*) were negatively associated with *F_ST_* (*p*=0.0405, *R*^2^=0.2668), with increasing values of *M* associated with reduced differentiation (Fig. S45).

**Figure 4.**
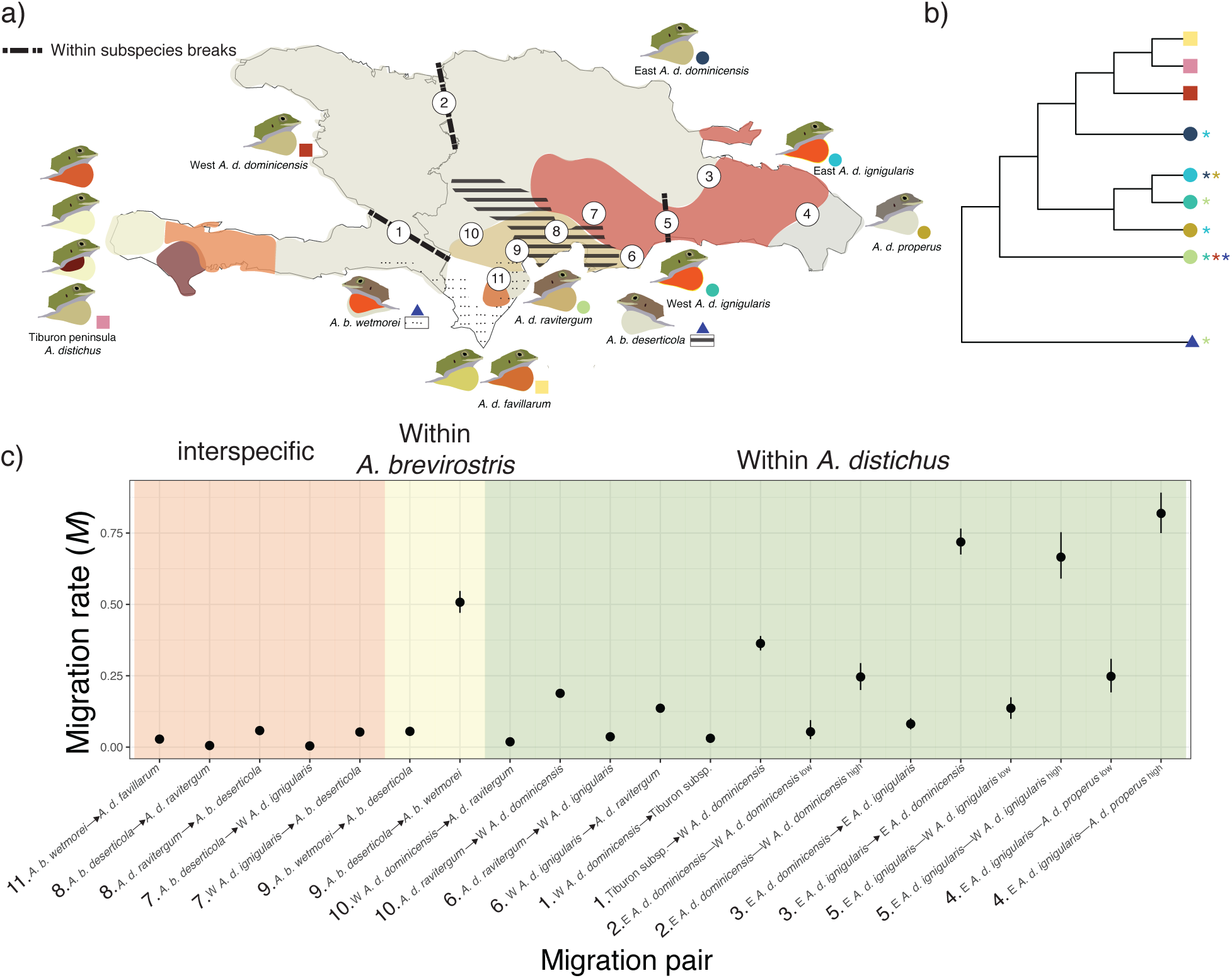
Gene flow within *A. distichus* is comparable to interspecific rates. a) We identified 11 contact zones between both sister and more distantly related taxa from which to infer migration. The subspecies ranges depicted here are modified from the maps of Schwartz (1968) and Arnold (1980). b) Subsetted species tree inferred using SVDquartets for the *A. distichus* species group (Fig. S31) with asterisks denoting previously observed instances of hybridization based on morphology or allozyme data (Schwartz, 1968; Webster, 1977; Case and Williams, 1984; Williams and Case, 1986). c) Estimates of migration rate (*M*) and 95% HPD are sorted from by the class of comparison (between *A. distichus* and *A. brevirostris*, within *A. brevirostris*, and within *A. distichus*) and in order of ascending *M*.

## 4 Discussion

Speciation is a dynamic, non-linear process wherein genomic, phenotypic, and ecological dimensions frequently decouple, creating severe challenges for species delimitation and phylogenetic inference. By comparing concatenated phylogenies with distance-based networks, species trees, and coalescent-based migration analyses in the *A. distichus* species group, we show how traditional, gene-flow-agnostic methods misrepresent evolutionary history in reticulate radiations. Our framework confirms deep lineage splits within the *A. brevirostris* complex, while revealing that *A. distichus* subspecies span a broad speciation continuum—ranging from porous, freely interbreeding polymorphic populations to strongly isolated lineages deserving species status. Crucially, our reconstruction of the *A. distichus* Y-chromosome phylogeny rejects the hypothesis that island-wide orange–yellow dewlap polymorphisms are maintained by a single, shared sex-linked haplotype. Together, our findings highlight how phenotypic and genomic divergence decouple during speciation, providing a robust framework for species delimitation among lineages undergoing differentiation.

### 4.1 Phylogenetic inference in a reticulate radiation

Concatenated phylogenetic approaches often yield strongly supported, yet misleading, topologies when applied to rapid radiations shaped by pervasive gene flow (Kubatko and Degnan, 2007; Mendes and Hahn, 2018; Steenwyk et al., 2023). Within *A. distichus*, we demonstrate that while species-tree models accounting for incomplete lineage sorting (SVDquartets and PoMo) expose deep conflicts obscured by concatenation, they remain inherently vulnerable to reticulation. Because they enforce strictly bifurcating histories, these ILS-aware models produce conflicting placements for highly admixed taxa. For instance, SVDquartets recovers *A. d. ignigularis* as monophyletic, whereas PoMo resolves it as paraphyletic, both with a high degree of statistical support (Figs. S31–32). Recognizing that bifurcating models fall short in reticulate systems, we combined distance-based network analyses (Neighbor-Net; (Bryant and Moulton, 2004)) with explicit gene-flow modeling under the multispecies coalescent with migration (MSC-M; (Flouri et al., 2023)) to characterize evolutionary relationships across the group (Figs. 1, 4). Rather than a simple bifurcating hierarchy, our network analysis recovers nine distinct clusters connected by varying degrees of reticulation (Fig. 1C). While well-delimited species-level lineages (i.e., members of the *A. brevirostris* complex) form discrete, relatively isolated clusters, subspecies of *A. distichus*, particularly those east of the Cordillera Central, form a complex mosaic of interconnected lineages. Importantly, Western and Eastern *A. d. ignigularis* populations display significant gene flow with neighboring *A. d. ravitergum* and *A. d. properus* populations, respectively (Fig. 4). This reticulate pattern aligns with the PoMo topology, which places Western *A. d. ignigularis* closer to *A. d. ravitergum* than to Eastern *A. d. ignigularis* (Fig. S32). Consequently, relying solely on bifurcating topologies fails to capture the biological reality of the *A. distichus* radiation, underscoring the necessity of integrating network representations with explicit demographic modeling to accurately delineate lineages.

### 4.2 Gene flow and lineage status along the speciation continuum

Genomic estimates of admixture expose a highly dynamic speciation continuum across the *A. distichus* complex. Crucially, our findings corroborate classic observations—dating back to early morphological (Schwartz, 1968) and allozyme (Webster, 1977; Case and Williams, 1984; Williams and Case, 1986) surveys, that suggested that gene flow occurred to varying degrees among *A. distichus* subspecies. Furthermore, our analyses confirm previous anecdotal evidence of rare, highly restricted introgression events between the distinct, yet often sympatric, species *A. distichus* and *A. brevirostris* (Webster (1977); Fig. 4). By scaling up from historical single-locus and morphological markers to genome-wide networks and multispecies coalescent migration models, our results demonstrate that these contact zones span a full spectrum of isolation, ranging from permeable population boundaries to entrenched species-level barriers.

#### 4.2.1 Phenotypic character displacement in the *A. brevirostris* complex

In agreement with their current taxonomic status, the Haitian species within the *A. brevirostris* complex–*A. caudalis*, *A. marron*, and *A. websteri* –represent well-defined, independently evolving lineages deserving of species recognition. Their well-supported reciprocal monophyly demonstrates that the *A. brevirostris* complex consists of distinct evolutionary species, corroborating the hypothesis that geographically structured dewlap variation in this group is driven by reproductive character displacement at contact zones (Webster and Burns, 1973; Lambert et al., 2013). In a textbook example of reproductive character displacement, *A. caudalis* is orange dewlapped in sympatry with yellow-dewlapped *A. brevirostris* in the south, but pale dewlapped when co-occurring with orange-dewlapped *A. websteri* in the north of its range.

We recover evidence for localized mitochondrial introgression and support for low levels of nuclear gene flow between *A. distichus sensu lato* and the *A. brevirostris* complex, corroborating early cytogenetic observations of rare putative hybrids (Webster, 1977). Although statistical support for this inter-complex genetic exchange is high (BF_10_ *≥* 20), the inferred magnitude of migration is nominal (*M ≤* 0.03 migrants per generation). We suggest that these low estimates do not reflect biologically meaningful contemporary gene flow capable of homogenizing lineage boundaries. Instead, because the multispecies coalescent with migration (MSC-M) framework integrates genetic signals across the entire divergence history of a clade, this baseline parameter estimate may represent rare episodic hybridization, ancient post-divergence introgression, or the retention of ancestral lineage sorting (Flouri et al., 2023; Thawornwattana et al., 2026, 2025; Cheng et al., 2026). Irrespective of the precise mechanism, our analyses confirm that despite widespread sympatry, *A. distichus* and the *A. brevirostris* complex remain distinct, reproductively isolated species complexes separated by strong barrier traits and only nominal genomic leakage (Fig. 4).

#### 4.2.2 A continuum of reproductive isolation and divergence within the *A. distichus* group

The Bahamian *A. distichus* lineages exhibit reduced genetic diversity consistent with singular founder events (Fig. S8), and are usually recovered as sister to the Eastern *A. d. dominicensis* population (Fig. 3, S12, but see Fig. S32). Because these island populations are geographically isolated by marine barriers, they follow independent evolutionary trajectories relative to Hispaniolan populations (De Queiroz, 2007). Consistent with other Greater Antillean *Anolis* radiations that have colonized adjacent archipelagos (Glor et al., 2005; Reynolds et al., 2020; Torres et al., 2025), we suggest that the Bahamian lineages represent a distinct evolutionary species. Although dynamic sea-level fluctuations across the Bahama Banks may have historically reconnected and fragmented these island populations (Pregill and Olson, 1981), isolation from Hispaniola preserves their distinctness despite potential internal paraphyly during interglacial periods.

Moving along the continuum toward more admixed Hispaniolan populations, the *A. distichus* lineages of southwestern Hispaniola present a compelling case of phenotypic polymorphism amid geographic isolation (Fig. 1). These populations—western *A. d. dominicensis*, *A. d. favillarum* of the Sierra de Baoruco, and the subspecies endemic to the Tiburon peninsula—are consistently recovered as reciprocally monophyletic. Remarkably, two of these monophyletic clades exhibit sharp color polymorphism across extremely narrow (*<* 1 km) contact zones (de Mello et al., 2026). Despite its dewlap color polymorphism, the combination of reciprocal nuclear monophyly, geographic isolation across deep physiographic depressions (e.g., the Neiba Valley), and severe contemporary habitat fragmentation supports treating *A. d. favillarum* and the Tiburon peninsula lineages as distinct, independently evolving species.

Unlike the *A. brevirostris* complex, where dewlap color divergence tracks strict species boundaries, eastern *A. distichus* populations reveal an evolutionary history dominated by gene flow. East of the Cordillera Central, *A. d. ignigularis*, *A. d. properus*, and eastern *A. d. dominicensis* exchange nuclear alleles at the highest rates quantified across the complex, despite displaying stark dewlap color differences (Fig. 4). In fact, *A. d. properus* and eastern *A. d. ignigularis* form a single, well-supported nuclear clade, indicating that *A. d. properus* represents a pale-yellow coastal morph within a broader, geographically widespread polymorphic lineage (Ng et al., 2016; MacGuigan et al., 2017; Myers et al., 2025). Underscoring this complexity, our genomic analyses reveal for the first time that the uniformly orange-dewlapped *A. d. ignigularis* comprises two cryptic lineages: an Eastern clade nested with *A. d. properus*, and a Western clade pulled toward *A. d. ravitergum* in network space. We hypothesize that this paraphyly reflects isolation-by-distance across Eastern Hispaniola (Fig. 2) coupled with secondary contact and localized introgression with *A. d. ravitergum* south of the Cordillera Central near Baní, which drew the Western *A. d. ignigularis* lineage toward *A. d. ravitergum* in both network analyses and ILS-aware bifurcating trees.

Despite this historical footprint of secondary contact, the contemporary nuclear genome of *A. d. ravitergum* remains remarkably distinct amidst the reticulate network of Eastern Hispaniola. Genomic migration rates between *A. d. ravitergum* and adjacent *A. d. ignigularis* populations are currently restricted to nominal baseline levels—comparable to the interspecific gene flow observed between the deeply divergent *A. brevirostris* and *A. distichus* complexes. Coupled with experimental evidence demonstrating that reproductive isolation is maintained by natural selection and postmating barriers (Geneva and Glor, 2025; Myers et al., 2025), these patterns confirm that *A. d. ravitergum* operates on an independent evolutionary trajectory. Consequently, *A. d. ravitergum* warrants elevation to full species status, as localized mitochondrial introgression and historical secondary contact have failed to erode its nuclear integrity.

Superimposed on this nuclear reticulation are multiple instances of mito-nuclear discordance across the radiation (Fig. S14). Despite exchanging nuclear genes with adjacent populations, Eastern *A. d. dominicensis* retains a deeply divergent, monophyletic mitochondrial clade sister to all other *A. distichus* subspecies. This discordance may reflect male-biased dispersal and female philopatry, wherein nuclear gene flow mediated by dispersing males swamps the nuclear genome without displacing local, maternally inherited mitochondrial lineages (Toews and Brelsford, 2012). In contrast, the discordant placement of the satellite island endemic *A. d. sejunctus*, sister to all Eastern *A. distichus* in the mitochondrial tree, points to the retention of an ancient founding organellar haplotype across marine channels that restrict immigration. Finally, the nesting of the mitochondrial genome of the range-restricted *A. d. ravitergum* within *A. d. ignigularis* is consistent with asymmetric introgression driven by demographic disparities (Ng and Glor, 2011). Because *A. d. ignigularis* maintains a vast effective population size relative to the localized *A. d. ravitergum*, neutral and advantageous mitochondrial haplotypes from donor *A. d. ignigularis* populations are prone to rapid fixation within *A. d. ravitergum* following secondary contact (Takahata, 1985).

### 4.3 Independent evolution of a convergent sexual signal

By reconstructing Y-chromosome history, we reject the hypothesis that the single sex-linked haplotype underlying orange and pale-yellow dewlaps in *A. d. favillarum* introgressed to drive color polymorphism across the entire group (de Mello et al., 2026). This aligns with theoretical expectations that strong local selection restricts the spread of adaptive alleles across contact zones (Barton and Hewitt, 1985). Sex-linked loci are particularly constrained: theory predicts that sex chromosomes contribute disproportionately to local adaptation under migration—a spatial analogue of the “large-X” effect—due to hemizygous exposure and sex-biased dispersal (Lasne et al., 2017; Presgraves, 2018). The confinement of this haplotype to *A. d. favillarum* indicates that divergent selection prevents the spread of locally adaptive signal haplotypes, demonstrating that orange and pale-yellow dewlaps in other *A. distichus* lineages evolved independently (Wooldridge et al., 2022).

Alternative mechanisms may explain these recurrent color patterns. First, superficially similar hues across Hispaniola could could be the byproduct of divergent biochemical pathways (Twomey et al., 2023), requiring spectral reflectance and chromatographic analyses to verify phenotypic convergence. Second, recurring phenotypes may involve autosomal loci. While the Y-chromosome explains most phenotypic variance in *A. d. favillarum*, autosomal regions also associate with orange/yellow variation (de Mello et al., 2026), suggesting that recurrent morphs could stem from standing ancestral variation at autosomal loci rather than sex-linked regions (Colosimo et al., 2005; Jones et al., 2012).

### 4.4 Taxonomic implications and future directions

Together, our genomic evidence demonstrates that the traditional subspecific taxonomy of *A. distichus sensu stricto* inadequately reflects the evolutionary history and lineage boundaries across the clade. While several named subspecies interbreed freely across continuous geographic contact zones, others maintain the genetic isolation and evolutionary independence expected of full species (De Queiroz, 2007). Because diagnostic morphological characters for these cryptic lineages remain elusive (Schwartz, 1968; Myers et al., 2020), formal taxonomic revisions must adopt an integrative framework combining genomic, ecological, and microhabitat data. Based on our phylogenomic reconstructions, explicit migration modeling, and previously established intrinsic barriers (Geneva and Glor, 2025), we advocate for the following primary taxonomic adjustments: (1) elevating the genetically isolated, nuclear-distinct subspecies *A. d. ravitergum* to full species status (*Anolis ravitergum*); (2) recognizing the monophyletic southwestern Hispaniolan lineages—including *A. d. favillarum* (*Anolis favillaceus*) and the distinct Tiburon peninsula population—as independent evolutionary species; (3) treating the geographically isolated Bahamian populations as a distinct insular species unit; and (4) synonymizing heavily admixed eastern Hispaniolan subspecies (i.e., *A. d. properus*) as regional phenotypic color morphs within a polymorphic, reticulate *Anolis distichus sensu stricto*.

Finally, while dewlap color variation does not reliably diagnose genome-wide lineage boundaries across *A. distichus*, visual signals remain ecologically paramount and strongly correlated with local light environments (Ng et al., 2013b). This pattern contrasts sharply with the sympatric *A. brevirostris* complex, where dewlap coloration closely mirrors species boundaries and facilitates reproductive character displacement (Webster and Burns, 1973; Lambert et al., 2013). Intriguingly, dewlap color polymorphisms in *A. distichus* are geographically concentrated in regions where its range overlaps with the *A. brevirostris* complex, suggesting that interspecific community dynamics—alongside abiotic light regimes—interact to shape sexual signal diversification. Our study confirms that lineages across multiple hierarchical tiers of divergence encompass the full spectrum of dewlap variation (Fig. 1). Determining whether this convergent phenotypic diversity is maintained by the retention of ancestral polymorphisms or independent molecular pathways across lineage boundaries remains a fundamental question. Developing the *A. distichus* species group as a premier squamate model for color pattern evolution—leveraging long-read assemblies, spatial transcriptomics, and functional genomics—will be essential to unraveling how sexual signals evolve along the speciation continuum.

## 6 Acknowledgments

We thank Jennifer Hackett of the University of Kansas Genome Sequencing Core for her help with establishing and carrying out the ddRADseq library preparation protocol. For providing the computation resources to analyze the data, we thank the helpful staff and administrators of Auburn University Easley high performance computing cluster. We thank Carlos Suriel, Gabriel de los Santos, Cristían Marte, and others at the Museo Nacional de Historia Natural and the Ministerio de Medio Ambiente y Recursos Naturales for facilitating our research in the Dominican Republic. For helping with the acquisition of samples for this research, we thank D. Luke Mahler, S. Blair Hedges, and James A. Schulte for sharing tissue samples; our valued collaborators from the Dominican Republic including Miguel Landestoy, Cristían Marte, Francis Reyes Polanco, Patricia Torres-Pineda, and Reveca Ramírez; and former members of the Glor Lab: Audrey Kelly, Daniel MacGuigan, Daniel Scantlebury, Julienne Ng, Kevin Chovanec, Ryane Logsdon, and Sabina Noll.

## 8 Data Availability Statement

Demultiplexed ddRADseq sequencing reads are available on the NCBI Sequence Read Archive under Bioproject PRJNA1065933. Processed data files used as inputs for analyses are available on the Dryad Digital Repository: https://datadryad.org/10.5061/dryad. m0cfxppm7. Scripts and parameter files used to carry out the analyses for this work can be found in a version-controlled Github repository (https://github.com/TannerMyers/barkanole-evolution).

## 9 Author Contributions

TCM, PLHM, and REG designed the study. TCM, PLHM, JT, AJG, and REG collected specimens. TCM, PLHM, and PMH performed lab work. TCM, PLHM, and JT analyzed the data. TCM wrote the first draft of the manuscript. All authors contributed to revising this manuscript.

