## Supplemental text and tables for "Adaptive signal evolution and gene flow drive variable speciation outcomes in bark anoles"

### 1 Supporting Information

##### List of Tables

1 **ddRADseq Sampling.** Information for the lizards sequenced for this study including voucher number, taxon assignment, sex, sampling locality coordinates. 8
2 **ND2 Sampling.** Information for the lizards with ND2 GenBank accessions including voucher number, taxon assignment, sex, and sampling locality coordinates. . . . 8
3 **Population genomic and phylogenetic datasets.** Information for the datasets used in this study including filtering parameters, number of individuals, and number of ddRAD markers. . . . 8
4 **IBD results.** . . . 8
5 **Sex chromosome and autosomal phylogeny support metrics.** . . . 8
6 **Sex chromosome and autosomal phylogeny discordance.** . . . 8
7 **Excess allele-sharing patterns with Dsuite.** . . . 8

8 **MSC-M convergence diagnostics**. 8
9 **Migration rates between geographically adjacent taxa in the *A. distichus* species group**. 8

List of Figures

1. **Distribution and population structure of *A. distichus*.** 9
2. **Distribution and population structure of the *A. brevirostris* species complex.** 10
3. **Mitochondrial DNA (ND2) sampling.** 11
4. **Number of ancestral populations (*K*) supported by ADMIXTURE for *A. brevirostris*.** 12
5. **Ancestry proportions for *A. brevirostris* across values of *K*.** 13
6. **Number of ancestral populations (*K*) supported by ADMIXTURE for *A. distichus*.** 14
7. **Ancestry proportions for *A. distichus* across values of *K*.** 15
8. **Nucleotide diversity, $\pi$ (represented as percentage along diagonal), and pairwise genetic differentiation ($F_{ST}$) within and among species of the *A. distichus* species group.** 16
9. ***A. brevirostris* IBD plots.** 17
10. ***A. d. dominicensis* IBD plots.** 18
11. ***A. d. ignigularis* IBD plots.** 19
12. **Maximum likelihood phylogeny of concatenated ddRAD loci.** 20
13. ***ND2* maximum likelihood phylogeny**. 21
14. **Mito-nuclear discordance in the *A. distichus* species group.** Maximum likelihood phylogenies estimated from ddRADseq loci and ND2 reveal discordant relationships, most notably with the placement of *A. d. ravitergum* and *A. d. sejunctus*. Gray lines connect individual specimens with discordant nuclear and mtDNA ancestry. Solid lines connect clades/individuals for which we have both ddRAD and mtDNA data while the dashed lines denote clades for which we obtained data from different specimens. 22
15. ***ND2* maximum likelihood phylogeny of *A. d. favillarum*.** 23
16. ***ND2* maximum likelihood phylogeny of western *A. d. dominicensis*.** 24
17. ***ND2* maximum likelihood phylogeny of *A. distichus* of the Tiburon penninsula.** 25
18. ***ND2* maximum likelihood phylogeny of *A. brevirostris* species complex.** 26
19. ***ND2* maximum likelihood phylogeny of eastern *A. d. dominicensis*.** 27
20. ***ND2* maximum likelihood phylogeny of *A. d. ravitergum* and *A. d. ignigularis*.** 28
21. ***ND2* maximum likelihood phylogeny of *A. d. properus* and *A. d. ignigularis*.** 29
22. ***ND2* maximum likelihood phylogeny of *A. d. properus* and *A. d. sejunctus*.** 30
23. **Pairwise genetic differentiation ($F_{ST}$) of the Y chromosome within and among populations of *A. distichus*.** 31

24. **Population structure patterns on the Y chromosome.** . . . . . . . . 32
25. **Maximum likelihood phylogeny of concatenated Y chromosome ddRAD loci.** . . . . . . . . . . . . . . . . . . . . . . . . . . . . . . . . . . . . . 33
26. **Maximum likelihood phylogeny of concatenated autosomal ddRAD loci.** . . . . . . . . . . . . . . . . . . . . . . . . . . . . . . . . . . . . . 34
27. **Maximum likelihood of concatenated ddRAD loci on the ancestral X chromosome (X1).** . . . . . . . . . . . . . . . . . . . . . . . . . . . 35
28. **Maximum likelihood of concatenated ddRAD loci on the neo-X chromosome (X2).** . . . . . . . . . . . . . . . . . . . . . . . . . . . . . 36
29. **X1 and X2 chromosome discordance.** . . . . . . . . . . . . . . . . . . . 37
30. **Summary of sex chromosome and autosome discordance.** . . . . . . 38
31. **SVDquartets species tree.** . . . . . . . . . . . . . . . . . . . . . . . . . . . 39
32. **Polymorphism-aware phylogenetic model (PoMo) species tree.** . . 40
33. ***f*-branch statistic heatmap.** . . . . . . . . . . . . . . . . . . . . . . . . . 41
34. **Traceplots for MSC-M analysis between *A. d. ignigularis* and *A. b. deserticola*.** . . . . . . . . . . . . . . . . . . . . . . . . . . . . . . . . . . . . 42
35. **Traceplots for MSC-M analysis between *A. d. favillarum* and *A. b. wetmorei*.** . . . . . . . . . . . . . . . . . . . . . . . . . . . . . . . . . . . . . 43
36. **Traceplots for MSC-M analysis between *A. d. ravitergum* and *A. b. deserticola*.** . . . . . . . . . . . . . . . . . . . . . . . . . . . . . . . . . . . 44
37. **Traceplots for MSC-M analysis between *A. b. deserticola* and *A. b. wetmorei*.** . . . . . . . . . . . . . . . . . . . . . . . . . . . . . . . . . . . . . 45
38. **Traceplots for MSC-M analysis between east *A. d. ravitergum* and west *A. d. ignigularis*.** . . . . . . . . . . . . . . . . . . . . . . . . . . . . 46
39. **Traceplots for MSC-M analysis between east *A. d. ravitergum* and west *A. d. dominicensis*.** . . . . . . . . . . . . . . . . . . . . . . . . . . . 47
40. **Traceplots for MSC-M analysis between east *A. d. dominicensis* and *A. d. ignigularis* populations.** . . . . . . . . . . . . . . . . . . . . . 48
41. **Traceplots for MSC-M analysis between the Tiburón penninsula subspecies of *A. distichus* and west *A. d. dominicensis*.** . . . . . . . 49
42. **Traceplots for MSC-M analysis between east *A. d. properus* and east *A. d. ignigularis*.** . . . . . . . . . . . . . . . . . . . . . . . . . . . . . 50
43. **Traceplots for MSC-M analysis between east and west *A. d. ignigularis* populations.** . . . . . . . . . . . . . . . . . . . . . . . . . . . . . . 51
44. **Traceplots for MSC-M analysis between east and west *A. d. dominicensis* populations.** . . . . . . . . . . . . . . . . . . . . . . . . . . . . 52
45. **Relationship between migration rate ($M$) and $F_{ST}$.** . . . . . . . . . . 53

#### 1.1 Supplemental Methods

##### 1.1.1 Sampling

Our initial sampling included 305 total representatives of all named taxa and geographic populations previously shown to be genetically distinct (e.g., *A. d. dominicensis*; Geneva et al. (2015); MacGuigan et al. (2017); Myers et al. (2025)) in the *Anolis distichus* species group (average $n = 12$ per lineage; Table 1). Specifically, from the *A. brevirostris* species complex, we sampled *A. brevirostris*, including *A. b. brevirostris* ($n = 10$), *A. b. deserticola* ($n = 17$), and *A. b. wetmorei* ($n = 13$); *A. caudalis* ($n = 3$), *A. marron* ($n = 3$), and *A. websteri* ($n = 3$). From the *A. distichus* species complex, we sampled *A. altavelensis* ($n = 2$), *A. d. aurifer* ($n = 4$), *A. d. biminiensis* ($n = 4$), *A. d. dapsilis* ($n = 5$), *A. d. distichoides* ($n = 5$), *A. d. distichus* ($n = 5$), Eastern *A. d. dominicensis* ($n = 52$), Northwestern *A. d. dominicensis* ($n = 6$), Southwestern *A. d. dominicensis* ($n = 12$), Western *A. d. dominicensis* ($n = 17$), *A. d. favillarum* ($n = 15$), *A. d. ignigularis* ($n = 44$), *A. d. ocior* ($n = 4$), *A. d. properus* ($n = 38$), *A. d. ravitergum* ($n = 28$), *A. d. sejunctus* ($n = 5$), *A. d. suppar* ($n = 5$), and *A. d. vinosus* ($n = 4$).

##### 1.1.2 ddRADseq library preparation

We used double digest restriction enzyme associated DNA sequencing (ddRADseq; (Peterson et al., 2012)) to obtain genome-scale markers for our populationg genomic and phylogenetic analyses. First, we extracted genomic DNA from tail tips or livers that had been preserved in 95% ethanol at -80°C. Second, we digested DNA using the SbfI and MspI restriction enzymes in one reaction over three hours at 37°C before barcoding and adding Illumina indexes. Third, we loaded 96 well plates by pooling four sets distinguished by Illumina indexes of 24 samples with unique barcodes on a 96 well plate. Fourth, we ligated adaptors and purified DNA fragments prior to size-selecting for a target range of 487 to 595 bp using a Blue Pippin size fractionator (Sage Science) and additional purification. Fifth, we amplified libraries using Illumina index primers and Phusion® polymerase. Finally, we sequenced the ddRADseq libraries using 150 bp paired-end reads on four lanes of an Illumina HiSeq X (Novogene Co.).

##### 1.1.3 Genomic dataset curation

For SNP-based analyses, we used bcftools v1.21 to filter the unfiltered VCF output by populations including all samples (Danecek et al., 2021). Prior to filtering for high-quality calls, we excluded individuals with large amounts of missing data by calculating per-sample missingness across the dataset and excluding all specimens missing data at ≥85% of sites (Cerca et al., 2021). This resulted in the elimination of 10 samples, leaving 297 in total. After excluding samples with excessive missingness, we applied depth, allele count, and additional missingness filters to retain high quality biallelic SNPs (“-m2 -M2 -v snps") with minor allele counts of ≥3 (“–min-ac 3"), ≥5x depth of coverage (“DP > 5"), and at least 90% complete (O’Leary et al., 2018). For analyses assuming unlinked markers, we used PLINK v1.9 to keep only 1 SNP per ddRAD locus (i.e., at least 1,000 bp apart) (Purcell et al., 2007). To minimize the impact of allelic dropout and maximize SNP yield, we applied these filters both across all samples and for each of the *A. distichus* and *A. brevirostris* species complexes separately.

For phylogenetic and coalescent-based migration analyses that require both variant and constant sites, we used the Stacks v2.65 module populations to export alignments of whole ddRAD loci harboring high quality SNPs in PHYLIP format. To compare autosomal and sex chromosome relationships, we considered only male specimens of *A. distichus*, verifying assignments made in the field by visualizing depth of coverage on Y-chromosome scaffolds. For the autosomal phylogeny, we filtered for SNPs that were at least 90% complete and 1 kb apart across the scaffolds corresponding to the autosomes in the AnoDis1.0 assembly (Falvey et al., 2026; de Mello et al., 2026). For the sex chromosomes, we filtered SNPs from the the Y-chromosome (scaffolds 12 $\leq$ 8.7 Mb, 13, and 16), the ancestral X chromosome (scaffold 8), and the neo-X (scaffold 12 $\geq$ 8.7 Mb) (Falvey et al., 2026; de Mello et al., 2026). Then, we dropped specimens from consideration that had $>$ 80% missing data across the Y-linked scaffolds before excluding sites with depth less than 5 and depth greater than three times the third quartile (72), keeping biallelic SNPs that were at least 50% complete. Due to the lack of markers that met thresholds combined with reduced recombination of the sex chromosomes, we did not filter based on physical linkage for the sex chromosome phylogenies.

To prepare alignments for our MSC-M analyses, we selected 8-12 representative specimens per lineage from each pair of geographically overlapping lineages. We then obtained ddRAD-seq loci with the "populations" module using the 90% complete linkage-filtered dataset as a starting point, additionally requiring that loci be present in both lineages ("–min-populations 2") and that the locus must be found across all specimens per lineage ("–min-samples-per-pop 1.0"). Specifics for each alignment may be found in Table S3.

###### 1.1.4 Quantifying autosome-sex chromosome phylogenetic discordance with Robinson-Foulds distances

Because standard maximum likelihood outputs from IQ-TREE2 are unrooted, the autosome and sex chromosome trees each comprise 179 internal bipartitions across $n = 182$ tips with a single arbitrary 3-way trifurcation at the output root. We calculated all Robinson-Foulds (RF) metrics, split counts, and support statistics on unrooted trees, ensuring that all metrics remain strictly root-independent. For clade-level and orientation analyses, we rooted phylogenies at the basal edge subtending the 34-tip {*A. d. ravitergum* + West *A. d. ignigularis*} clade (which displays maximal SH-aLRT 100 / UFBoot 100 support across autosomes, $X_1$, and $X_2$), and validated root sensitivity under diameter-based midpoint rooting.

We quantified pairwise topological distance using the unrooted Robinson-Foulds symmetric difference metric across all tree pairs. We canonicalized internal bipartitions to the minority tip set, excluding trivial splits (singletons and the full leaf set). We calculated normalized Robinson-Foulds distance (nRF) as $\text{nRF} = \text{RF}/(2 \times [n-3])$, where $n = 182$ tips, yielding a theoretical maximum RF distance of $\text{RF}_{\text{max}} = 358$. For each pairwise comparison, we enumerated the numbers of shared bipartitions, splits unique to Tree 1, and splits unique to Tree 2.

To discriminate between deep macro-evolutionary incongruence and localized tip-level reticulation (e.g., within-clade shuffling due to incomplete lineage sorting or contemporary introgression), we performed a collapsed-backbone Robinson-Foulds analysis. For each phylogeny, we contracted each maximal monophyletic subspecific block into a single composite terminal. We partitioned paraphyletic taxa into multiple numbered blocks according to their discrete phylogenetic core groupings. We then compared reduced backbone topologies over the intersection of shared taxon blocks and computed backbone RF as $\mathrm{RF}_{\mathrm{bb}}/(2 \times [n_{\mathrm{shared\_blocks}} - 3])$.

#### 1.2 Supplemental Results

##### 1.2.1 Topological discordance between autosomes and sex chromosomes

All four trees resolved 179 internal bipartitions across the 182 male specimens. The autosomal phylogeny exhibited the highest overall statistical support (mean SH-aLRT = 92.3, median = 100.0; mean UFBoot = 97.0, median = 100.0), with 79.3% of nodes receiving SH-aLRT ≥ 95 and 82.1% receiving UFBoot ≥ 95 (63.7% receiving SH-aLRT = 100; Table S5). The ancestral X ($X_1$) tree demonstrated intermediate support (mean SH-aLRT = 82.3, median = 95.3; mean UFBoot = 87.4, median = 96.0; 52.5% UFBoot ≥ 95), which the neo-X ($X_2$) tree closely matched (mean SH-aLRT = 81.2, median = 92.9; mean UFBoot = 87.9, median = 95.0; 53.1% UFBoot ≥ 95). The Y-chromosome tree exhibited lower node support (mean SH-aLRT = 73.1, median = 86.5; mean UFBoot = 92.0, median = 99.0; 40.8% SH-aLRT ≥ 95 and 64.8% UFBoot ≥ 95), consistent with the non-recombining, male-limited nature and reduced effective population size of the Y chromosome.

We recovered substantial phylogenetic discordance across all genomic compartments at the individual tip level, yet topological distances adhered to a strict biological hierarchy dictated by transmission mode and effective population size (Table S6):

1. **Autosomes vs. Ancestral X ($X_1$) [Highest Concordance]:** The ancestral X chromosome ($X_1$) was topologically closest to the autosomal genome, sharing 73 bipartitions (RF = 212/358, nRF = 0.592; 106 unique splits per tree). This close concordance reflects the relatively large effective population size of the ancestral X ($N_e = 3/4$ that of autosomes) and its biparental recombination in females, which preserves the deep organismal branching history.

2. **Autosomes vs. Y and Neo-X ($X_2$):** The autosomal phylogeny was next closest to the Y chromosome (RF = 226/358, nRF = 0.631; 66 shared splits, 113 unique splits per tree), followed by the autosomes versus the neo-X ($X_2$) chromosome (RF = 256/358, nRF = 0.715; 51 shared splits, 128 unique splits per tree).

3. **Sex Chromosome Cross-Comparisons (Y vs. $X_1$ and $X_1$ vs. $X_2$):** Topological divergence between sex-chromosome compartments exceeded autosomal-sex comparisons. Pairwise comparison of the Y chromosome versus ancestral X ($X_1$) yielded RF = 262/358 (nRF = 0.732; 48 shared splits, 131 unique splits). Similarly, direct comparison between the two X-chromosome compartments—the ancestral X ($X_1$) versus the neo-X ($X_2$)—revealed high tip-level incongruence: RF = 268/358 (nRF = 0.749; 45 shared splits, 134 unique splits).

4. **Y Chromosome vs. Neo-X ($X_2$) [Maximum Divergence]:** The maximum topological divergence across the entire dataset occurred between the Y chromosome and

the neo-X ($X_2$) chromosome, which shared only 35 internal bipartitions (RF = 288/358, nRF = 0.804; 144 unique splits in each tree). This pronounced discordance reflects cumulative divergence between a strictly patrilineal, non-recombining locus (Y) and a fused neo-sex chromosome ($X_2$) undergoing rapid sorting and sex-biased introgression.

To resolve whether the elevated tip-level discordance among sex chromosomes (particularly $X_1$ vs. $X_2$, nRF = 0.749) stems from ancient, deep macro-evolutionary discordance or shallow, localized tip reticulation, we collapsed maximal monophyletic taxon blocks into single terminals (Table S6): In the Ancestral X versus Neo-X comparison, the dataset contracted into 23 monophyletic blocks for $X_1$ and 25 blocks for $X_2$, yielding 20 shared backbone terminals. Upon collapsing terminal tips, the Robinson-Foulds distance dropped by 68.6%—from nRF = 0.749 at the tip level down to a remarkably low backbone distance of RF = 8/34 (nRF = 0.235). Across the organismal backbone, 13 bipartitions were identical between $X_1$ and $X_2$, with only 4 splits unique to each partition. This fundamental result proves that the ancestral X and neo-X chromosomes share nearly identical macro-phylogenetic backbones, and that fine-scale tip interleaving and reticulation within terminal subspecific hybrid zones (specifically within the Eastern complex hybrid zone between *A. d. properus* and *A. d. ignigularis*) overwhelmingly drive their elevated tip-level discordance.

In the autosomes–Y-chromosome comparison, the dataset contracted into 25 autosomal blocks and 21 Y-chromosome blocks, sharing 19 backbone terminals. Here, backbone discordance dropped more moderately from nRF = 0.631 at the tip level to RF = 14/32 (nRF = 0.438), sharing 9 splits with 7 splits unique to each partition. Unlike the $X_1$ vs. $X_2$ comparison, the Y chromosome retains substantial deep backbone discordance attributable to genuine lineage capture events—most notably the 11-tip East *A. d. dominicensis* lineage that forms a 100/100 supported sister group to the Bahamas clade on the Y chromosome, but nests within the remaining East *A. d. dominicensis* populations on the autosomes.

Curated deep clade analysis showed that island and peripheral lineages maintained high genealogical fidelity across all four genomic compartments. The Bahamas clade (19 tips) formed a completely monophyletic clade with near-maximal support across all four genome compartments (autosomes: 100/100; Y: 99.5/100; $X_1$: 100/100; $X_2$: 100/100). Similarly, the basal Hispaniolan clade *A. d. ravitergum* + West *A. d. ignigularis* (34 tips) maintained 100/100 support across autosomes, $X_1$, and $X_2$, but broke on the Y chromosome, where West *A. d. ignigularis* intermingled with the eastern complex in a 60-tip basal assemblage (97/96 support).

Individual-level cross-genome analysis revealed that 126 of 182 males (69.2%) grouped concordantly within their respective taxon cores between autosomes and Y, while 122 males (67.0%) showed concordance between $X_1$ and $X_2$. In $X_1$ vs. $X_2$, 60 individuals formed strays in at least one partition (16 in $X_1$ only [8.8%], 25 in $X_2$ only [13.7%], and 19 in both [10.4%]). The 19 shared strays concentrated within the Eastern clade (11 *A. d. properus*, 1 East *A. d. ignigularis*, and 6 West *A. d. ignigularis*), and 1 Bahamas (*A. d. distichoides*: JASIL3373), demonstrating that sex-chromosomal discordance is largely circumscribed to contact zones on Hispaniola (Fig. S29-30).

Table S1. **ddRADseq Sampling.** Information for the lizards sequenced for this study including voucher number, taxon assignment, sex, sampling locality coordinates.

Table S2. **ND2 Sampling.** Information for the lizards with ND2 GenBank accessions including voucher number, taxon assignment, sex, and sampling locality coordinates.

Table S3. **Population genomic and phylogenetic datasets.** Information for the datasets used in this study including filtering parameters, number of individuals, and number of ddRAD markers.

Table S4. **IBD results.**

Table S5. **Sex chromosome and autosomal phylogeny support metrics.**

Table S6. **Sex chromosome and autosomal phylogeny discordance.**

Table S7. **Excess allele-sharing patterns with Dsuite.**

Table S8. **MSC-M convergence diagnostics**.

Table S9. **Migration rates between geographically adjacent taxa in the *A. distichus* species group**.

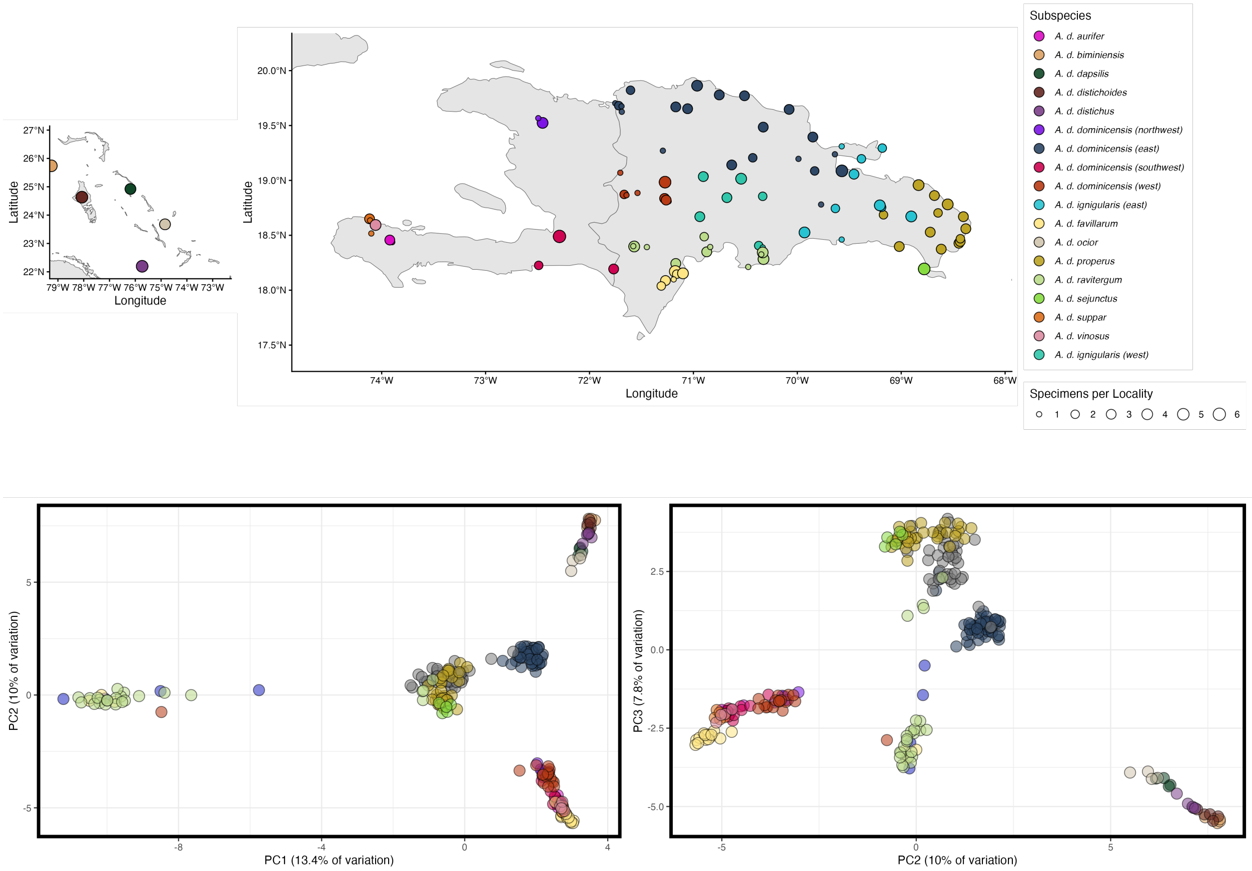

Figure S1. **Distribution and population structure of *A. distichus*.**

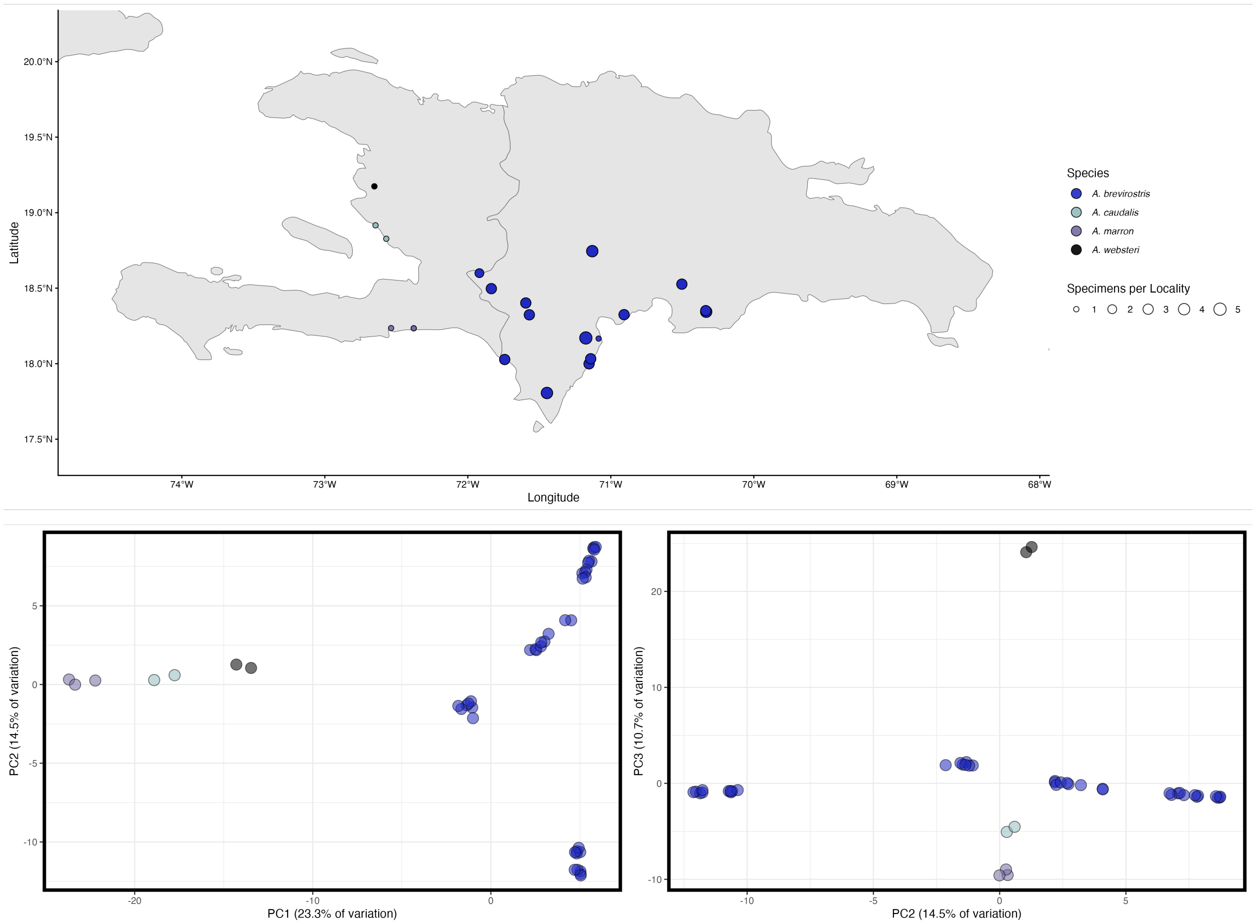

Figure S2. **Distribution and population structure of the *A. brevirostris* species complex.**

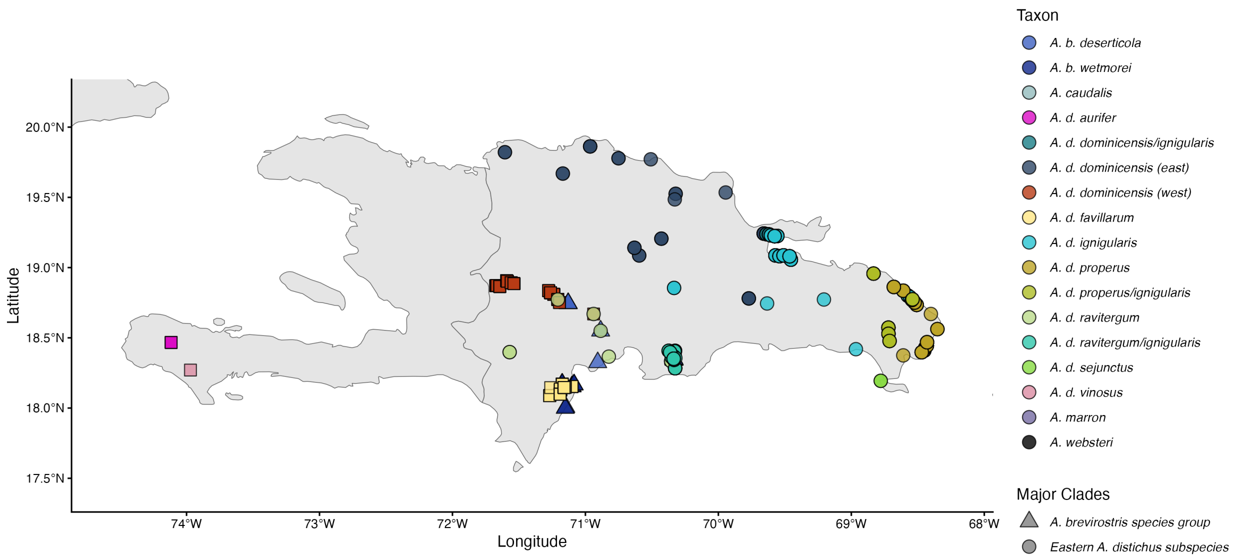

Figure S3. **Mitochondrial DNA (ND2) sampling.**

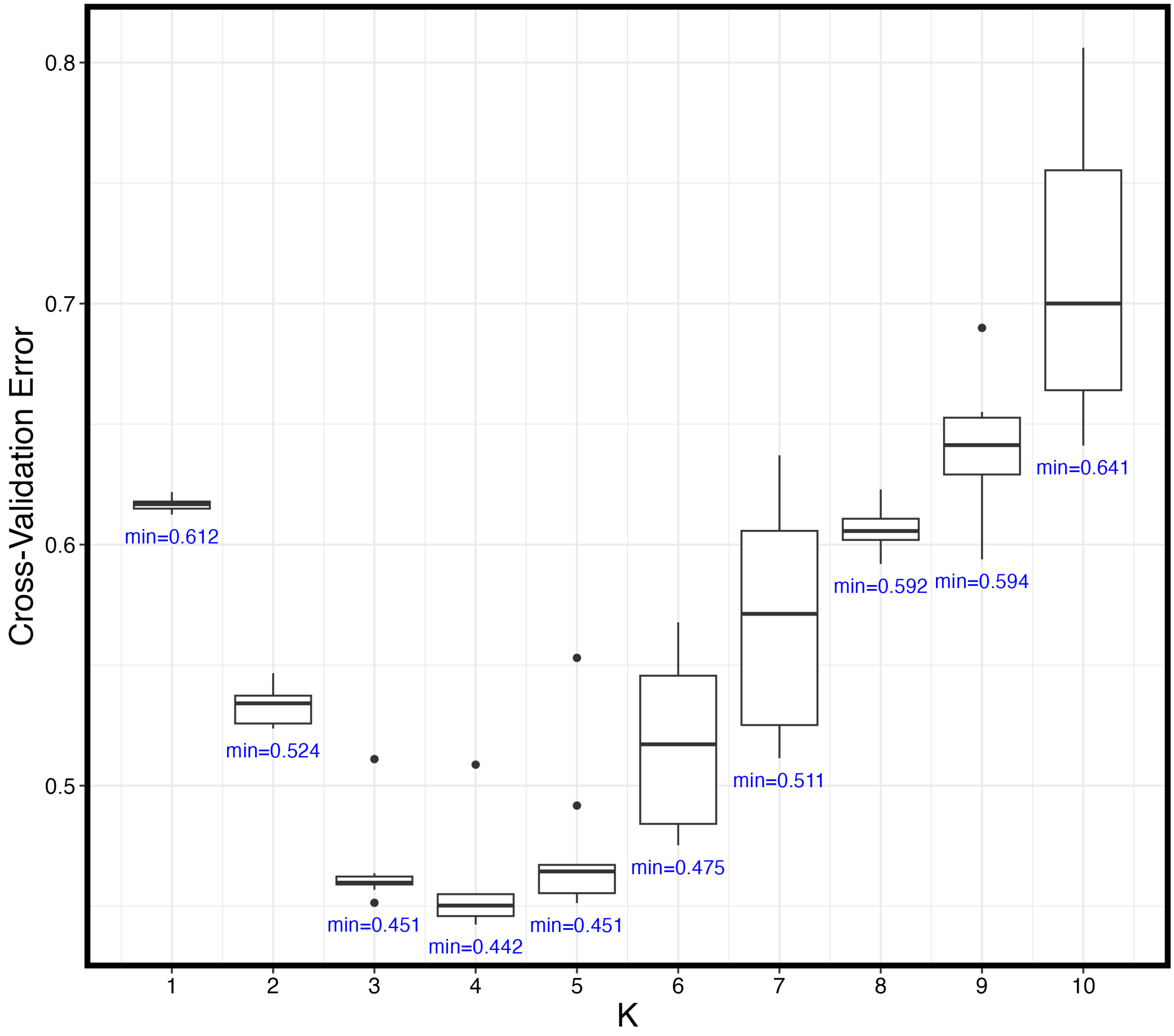

Figure S4. **Number of ancestral populations (*K*) supported by ADMIXTURE for *A. brevirostris*.**

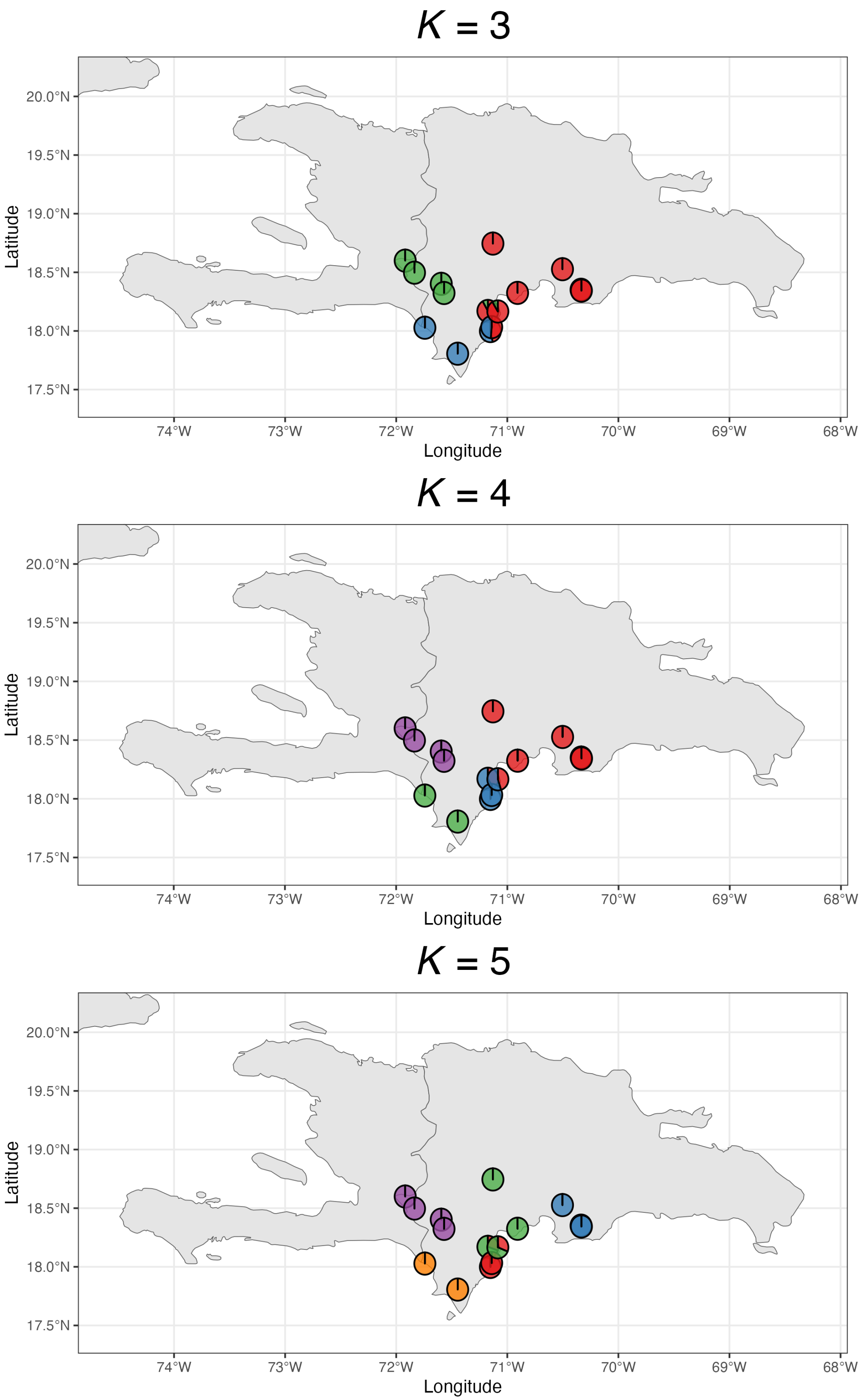

Figure S5. **Ancestry proportions for *A. brevirostris* across values of $K$.**

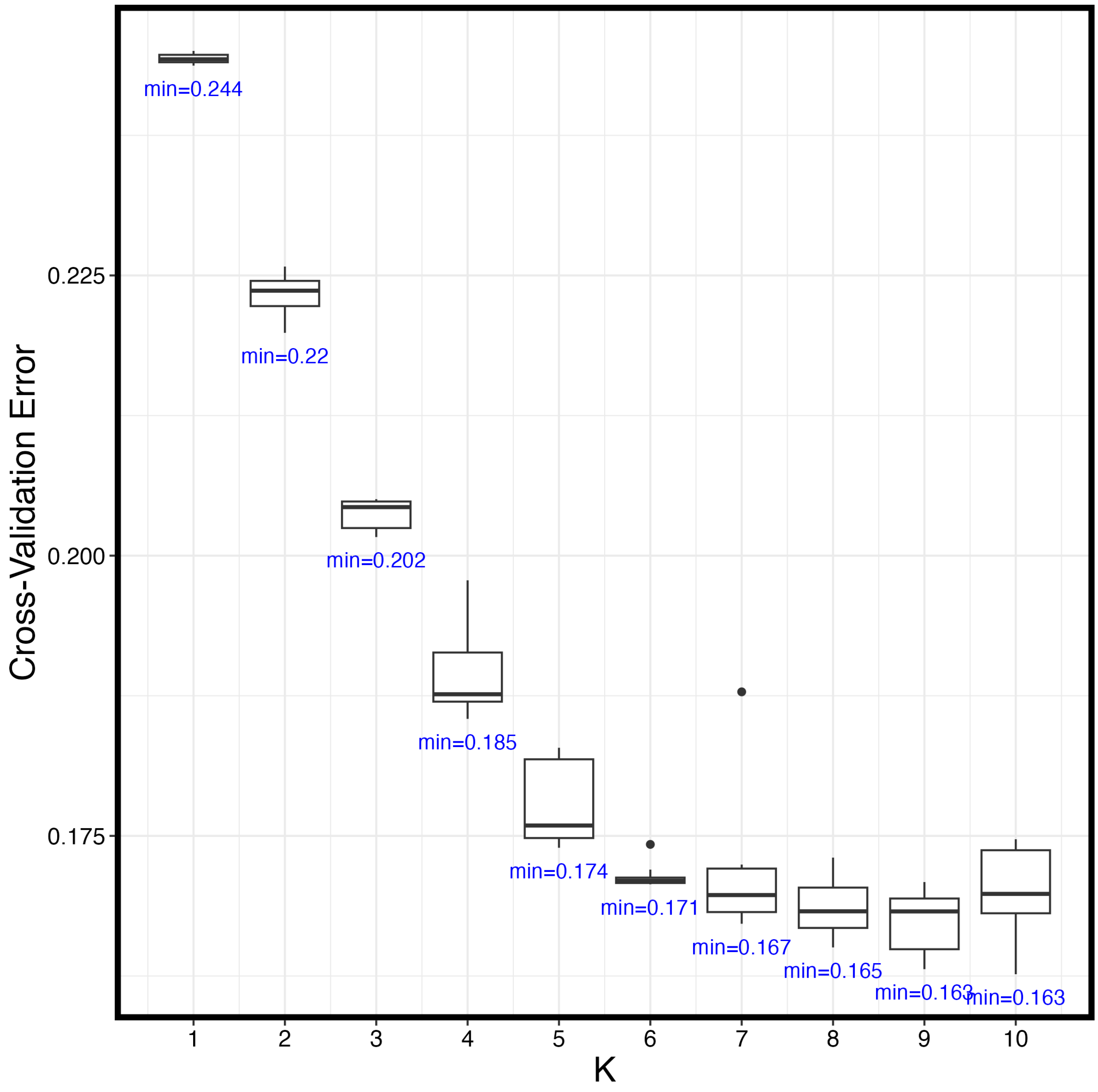

Figure S6. **Number of ancestral populations ($K$) supported by ADMIXTURE for *A. distichus*.**

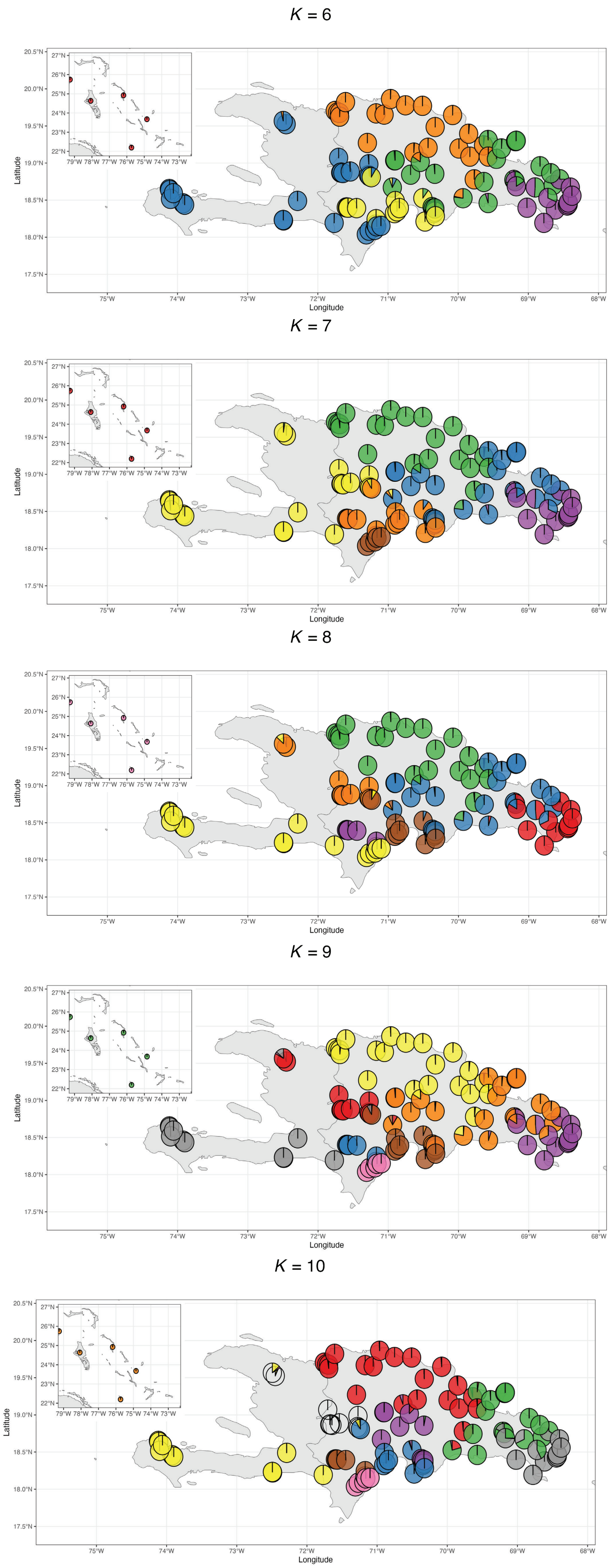

Figure S7. **Ancestry proportions for *A. distichus* across values of *K*.**

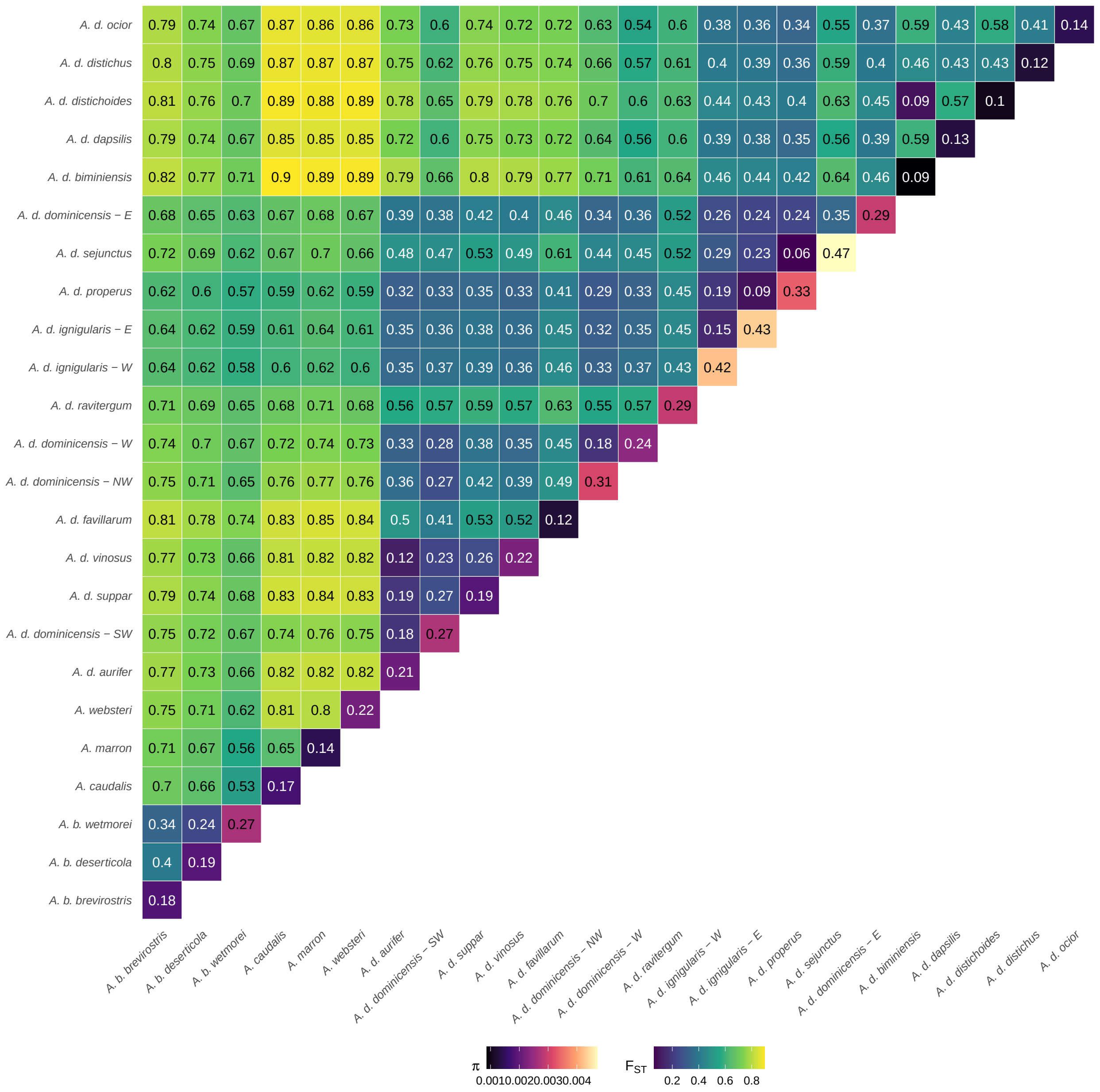

Figure S8. **Nucleotide diversity, $\pi$ (represented as percentage along diagonal), and pairwise genetic differentiation ($F_{ST}$) within and among species of the *A. distichus* species group.**

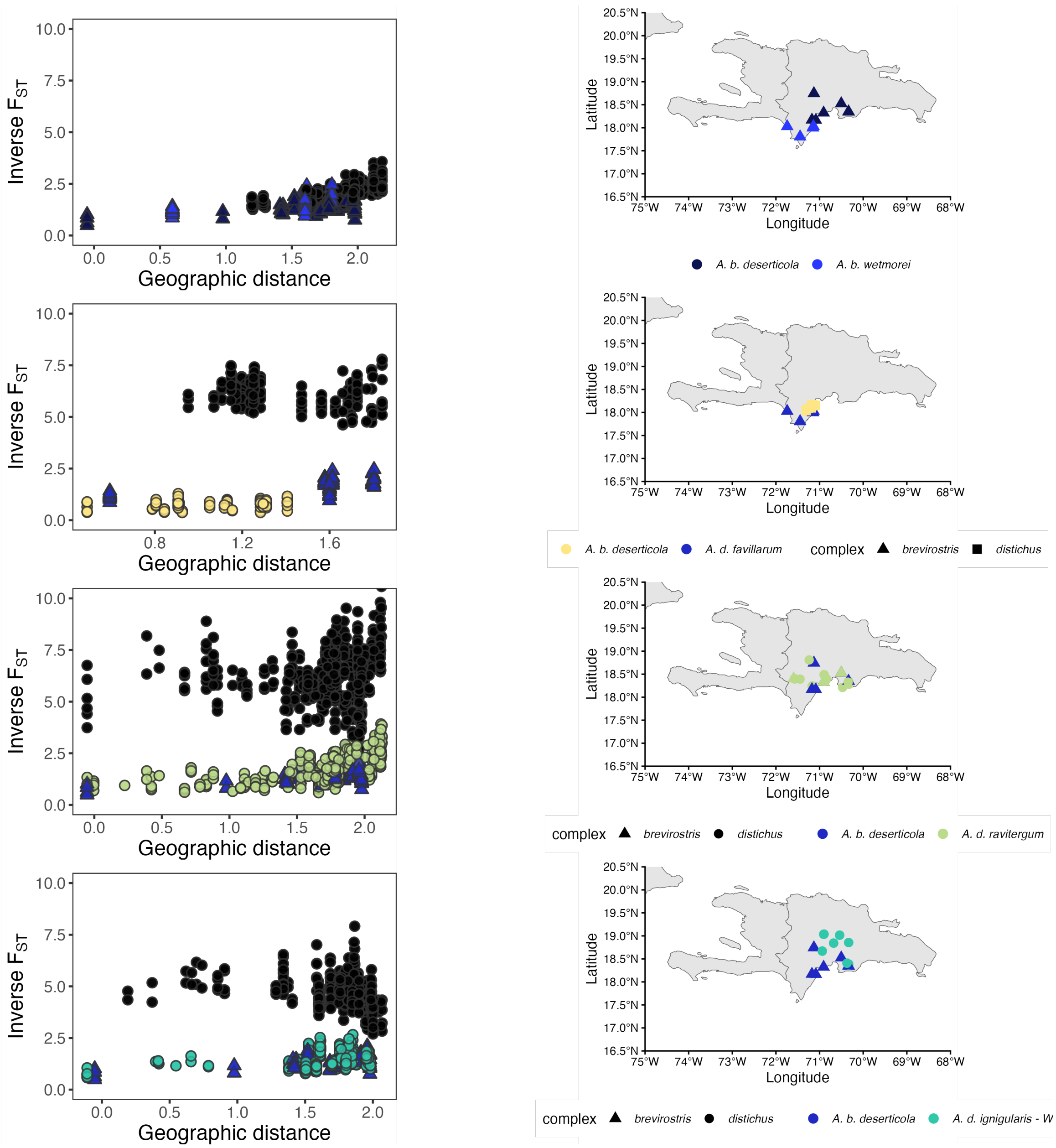

Figure S9. ***A. brevirostris* IBD plots.**

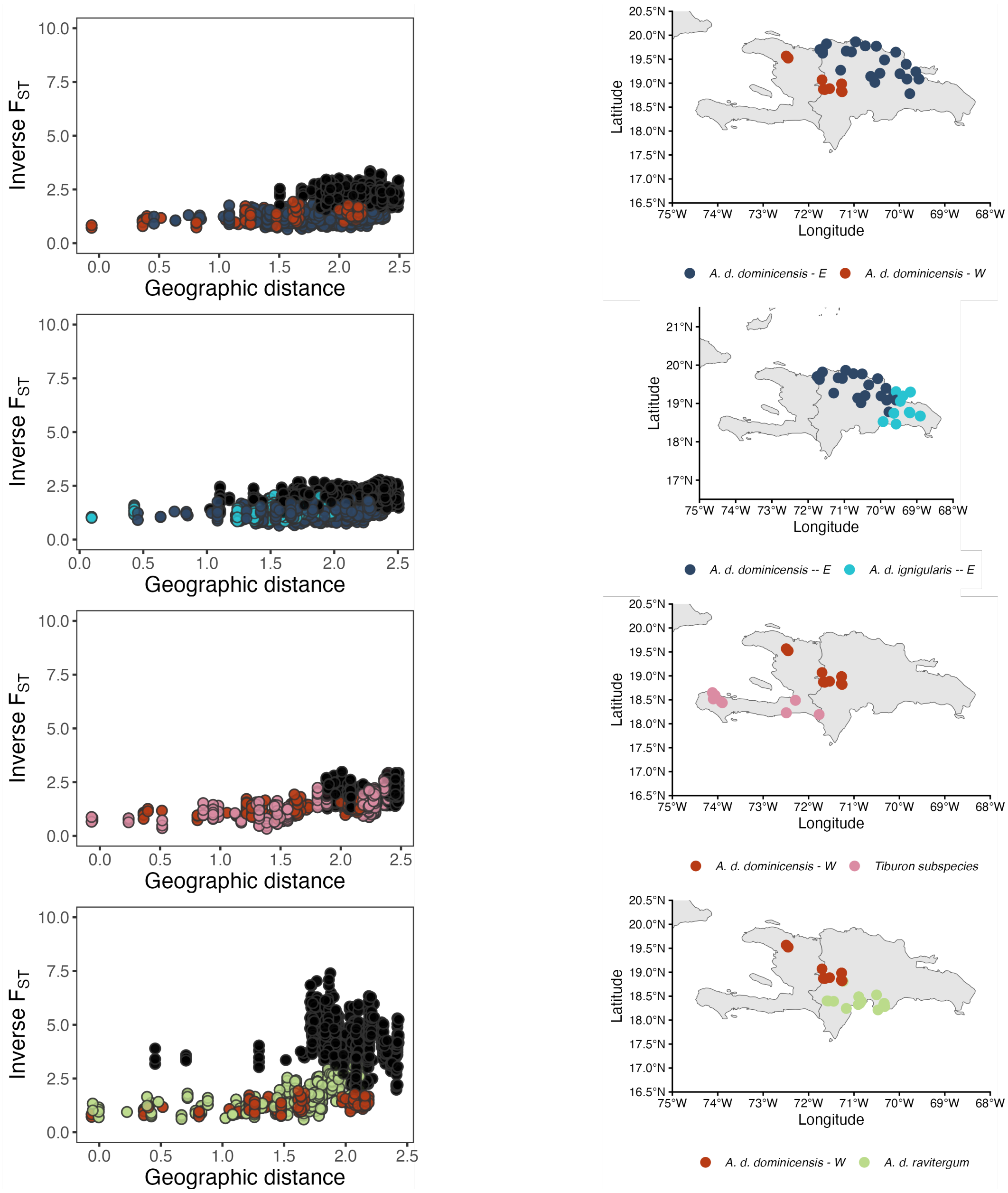

Figure S10. ***A. d. dominicensis*** **IBD plots.**

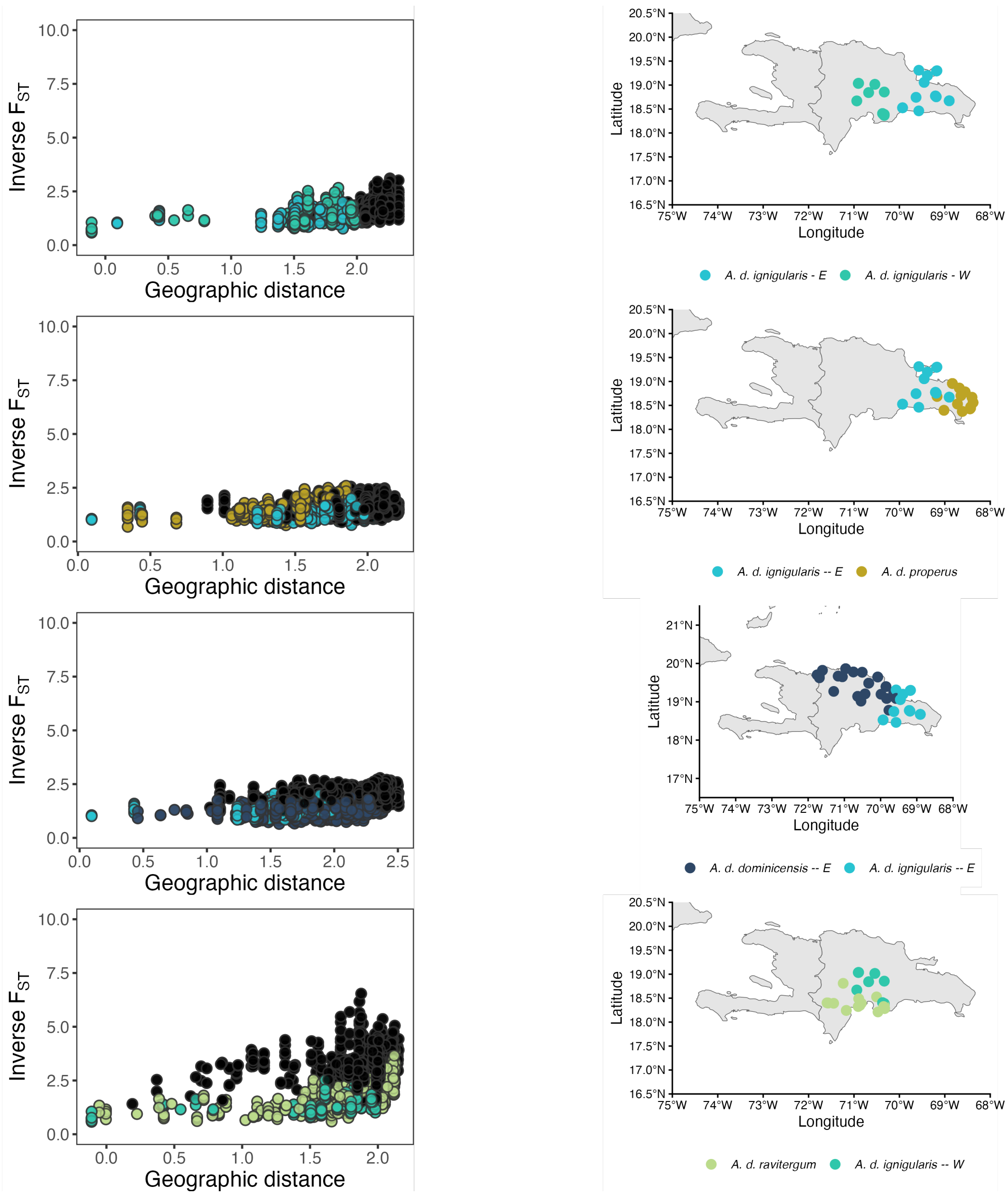

Figure S11. ***A. d. ignigularis* IBD plots.**

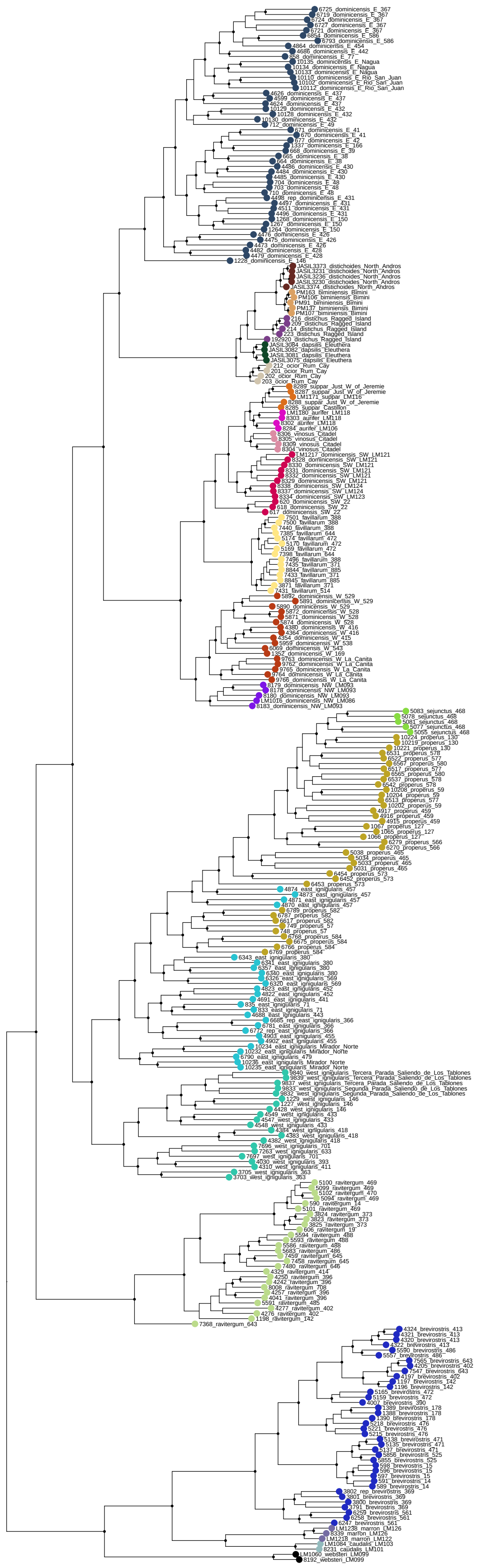

Figure S12. **Maximum likelihood phylogeny of concatenated ddRAD loci.**

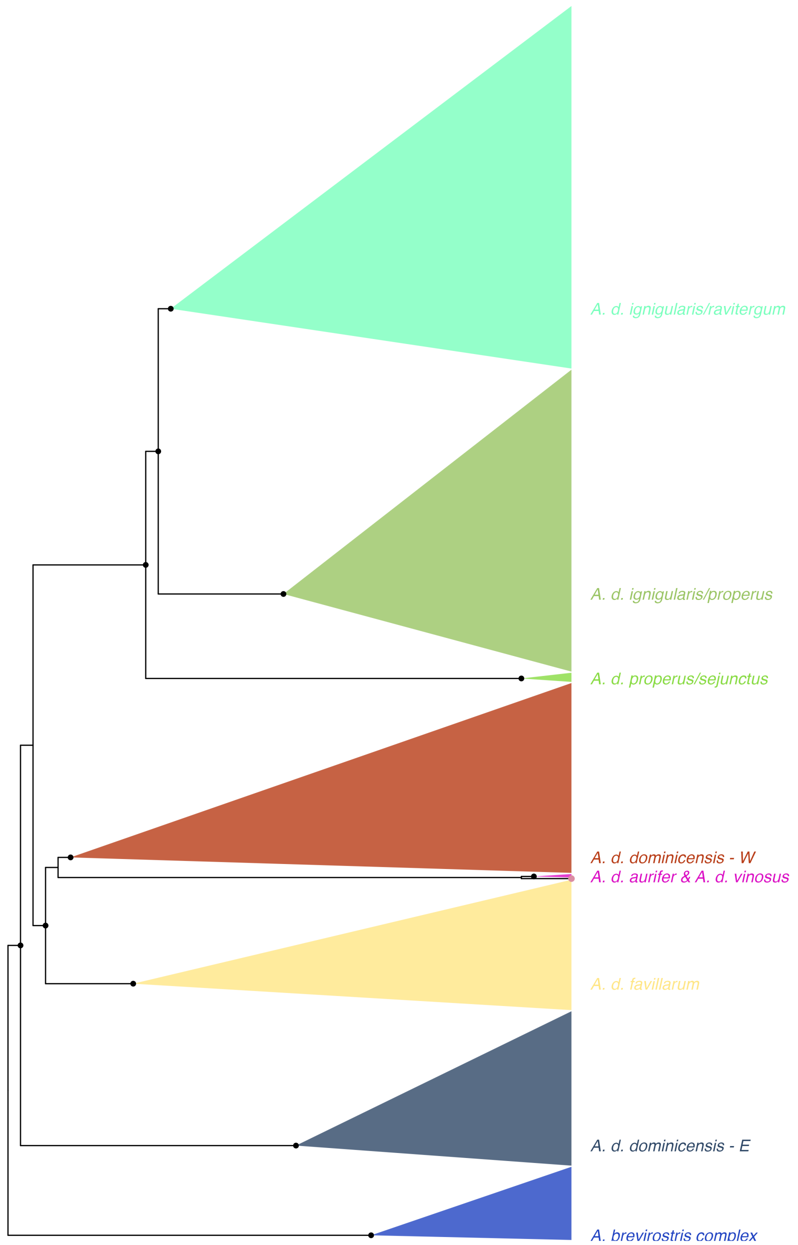

Figure S13. ***ND2* maximum likelihood phylogeny.**

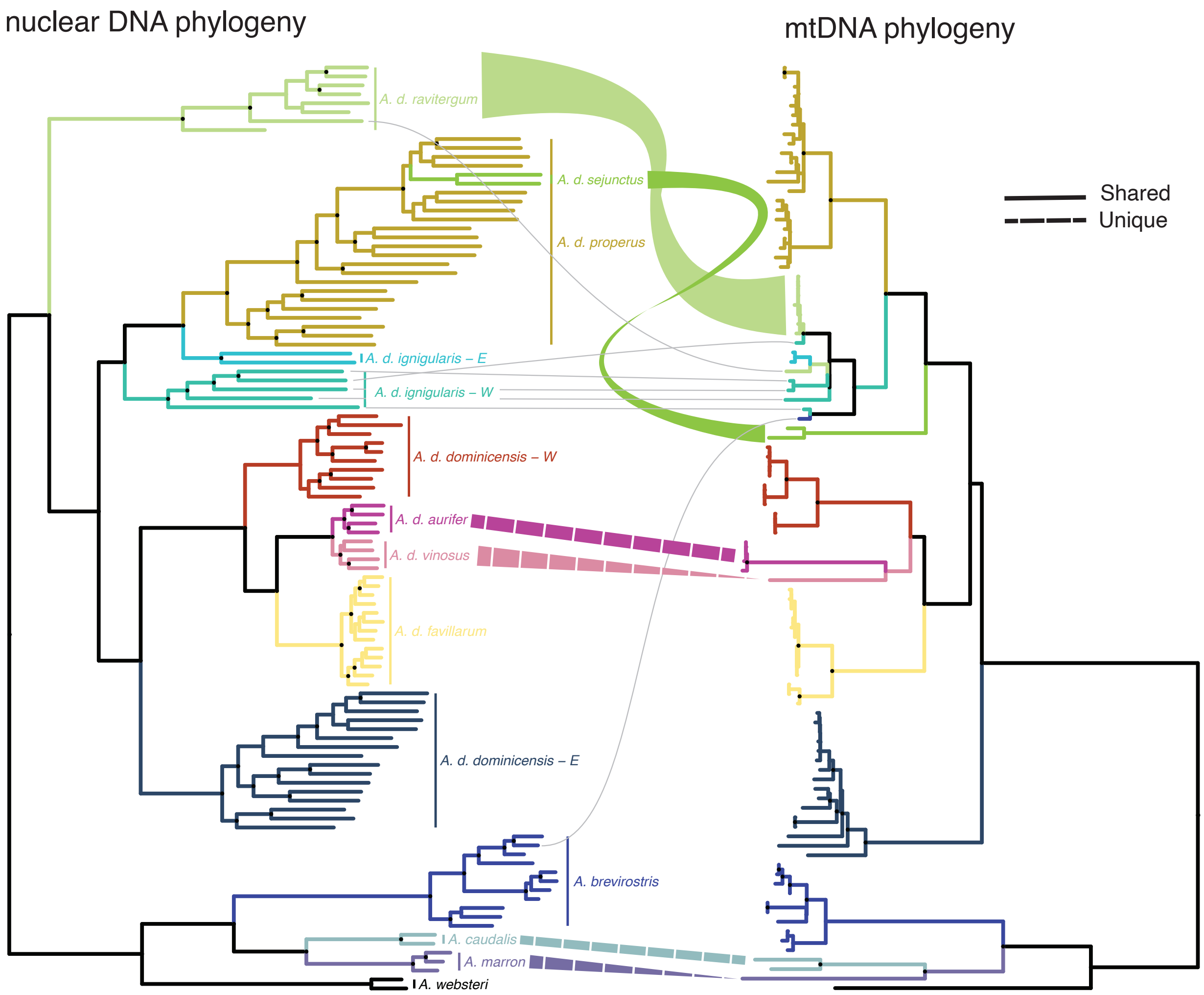

Figure S14. **Mito-nuclear discordance in the *A. distichus* species group.** Maximum likelihood phylogenies estimated from ddRADseq loci and ND2 reveal discordant relationships, most notably with the placement of *A. d. ravitergum* and *A. d. sejunctus*. Gray lines connect individual specimens with discordant nuclear and mtDNA ancestry. Solid lines connect clades/individuals for which we have both ddRAD and mtDNA data while the dashed lines denote clades for which we obtained data from different specimens.

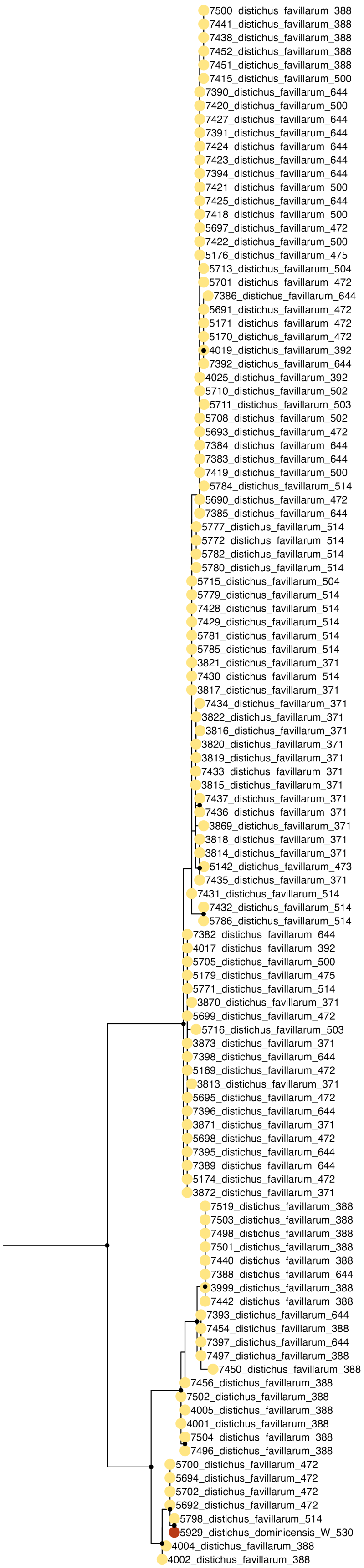

Figure S15. ***ND2* maximum likelihood phylogeny of *A. d. favillarum*.**

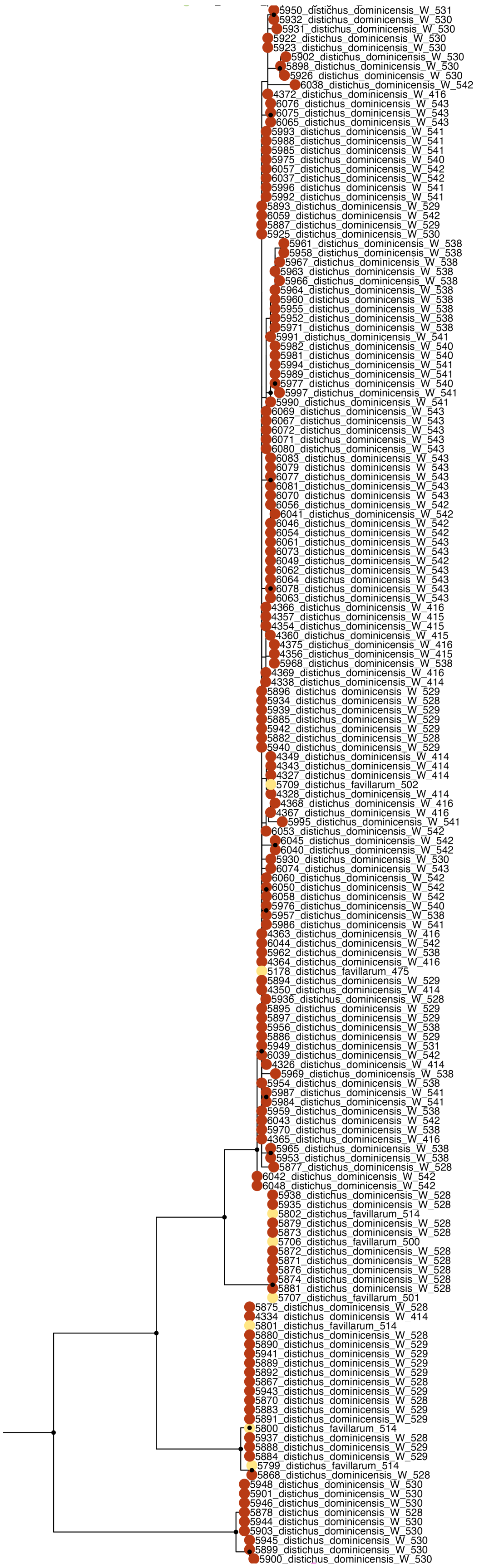

Figure S16. ***ND2*** **maximum likelihood phylogeny of western** ***A. d. dominicensis*****.**

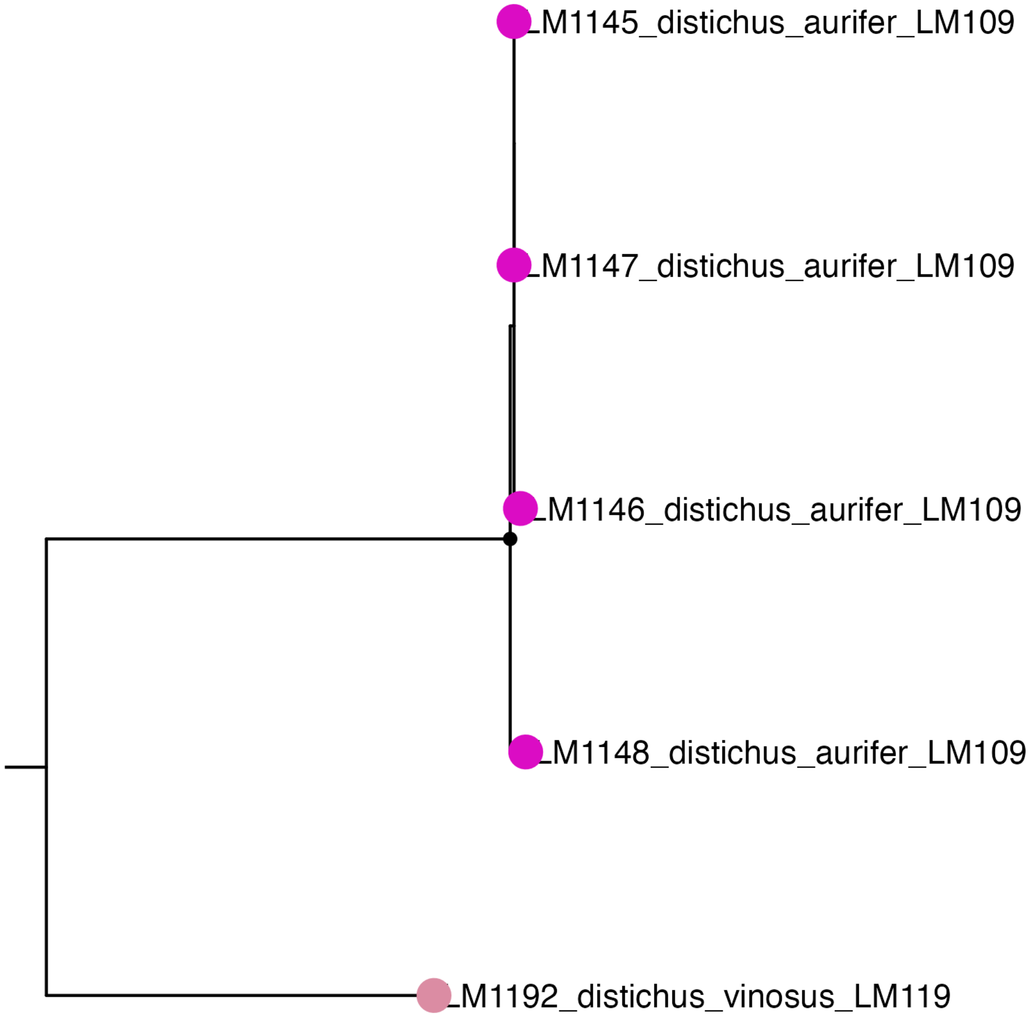

Figure S17. ***ND2* maximum likelihood phylogeny of *A. distichus* of the Tiburon peninsula.**

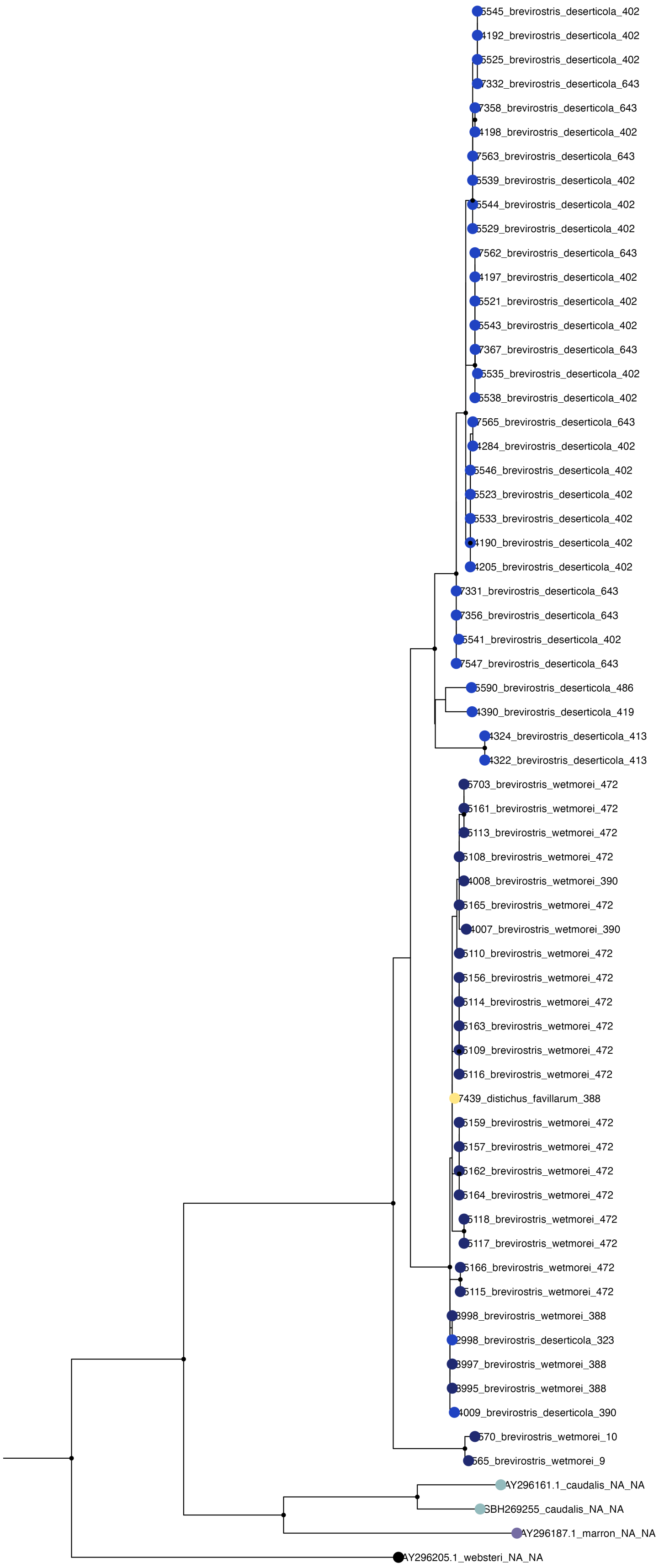

Figure S18. ***ND2* maximum likelihood phylogeny of *A. brevirostris* species complex.**

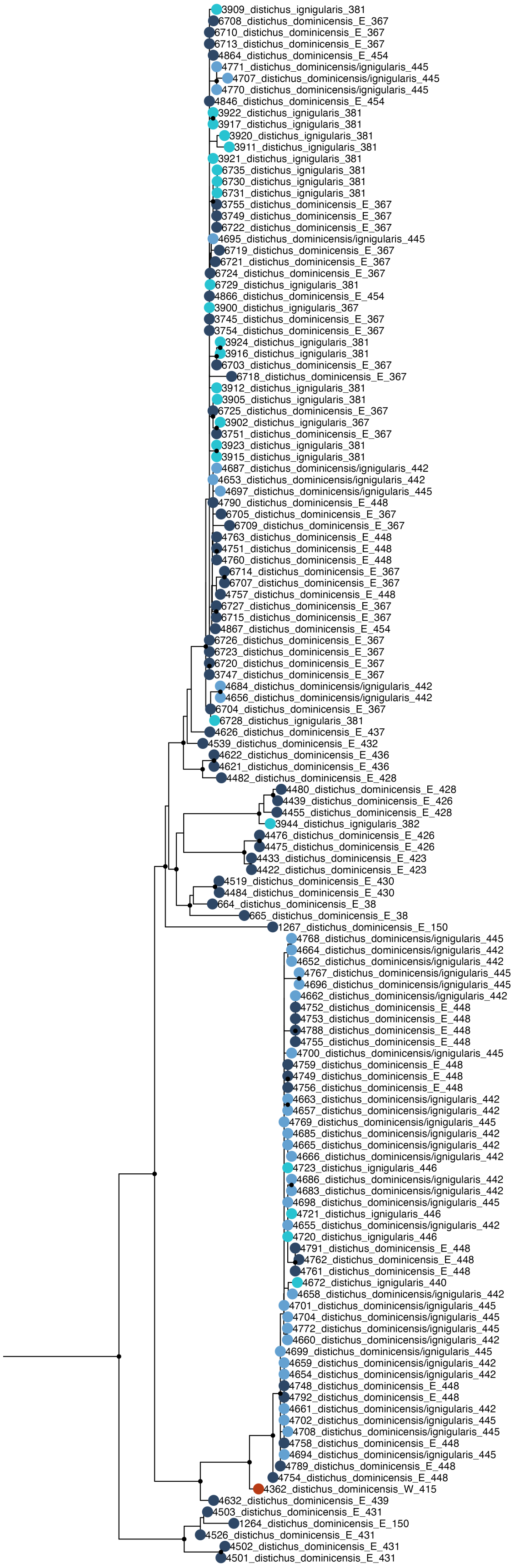

Figure S19. ***ND2*** **maximum likelihood phylogeny of eastern** ***A. d. dominicensis*****.**

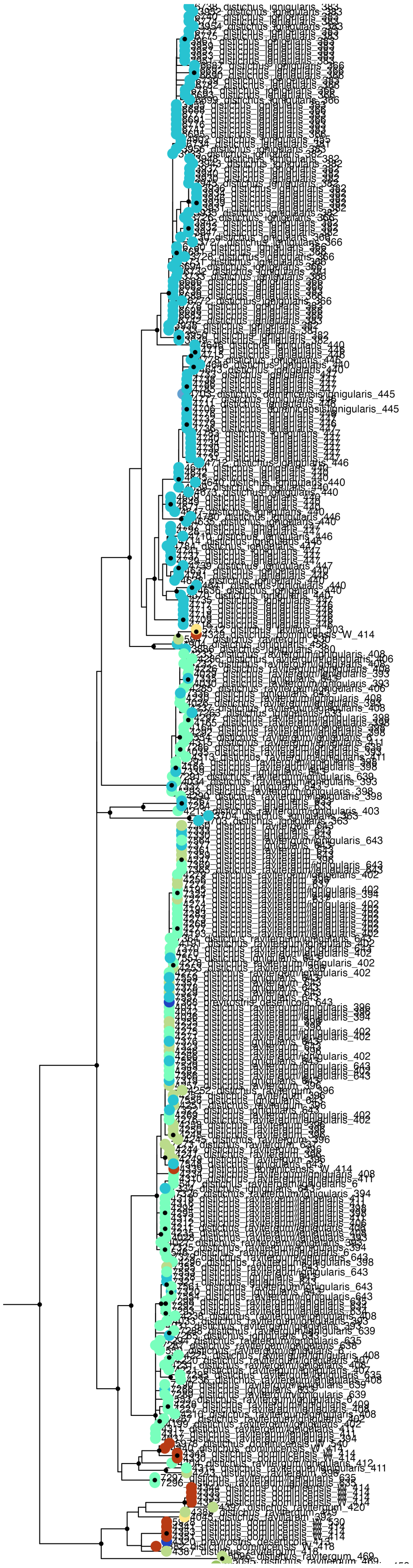

Figure S20. ***ND2* maximum likelihood phylogeny of *A. d. ravitergum* and *A. d. ignigularis*.**

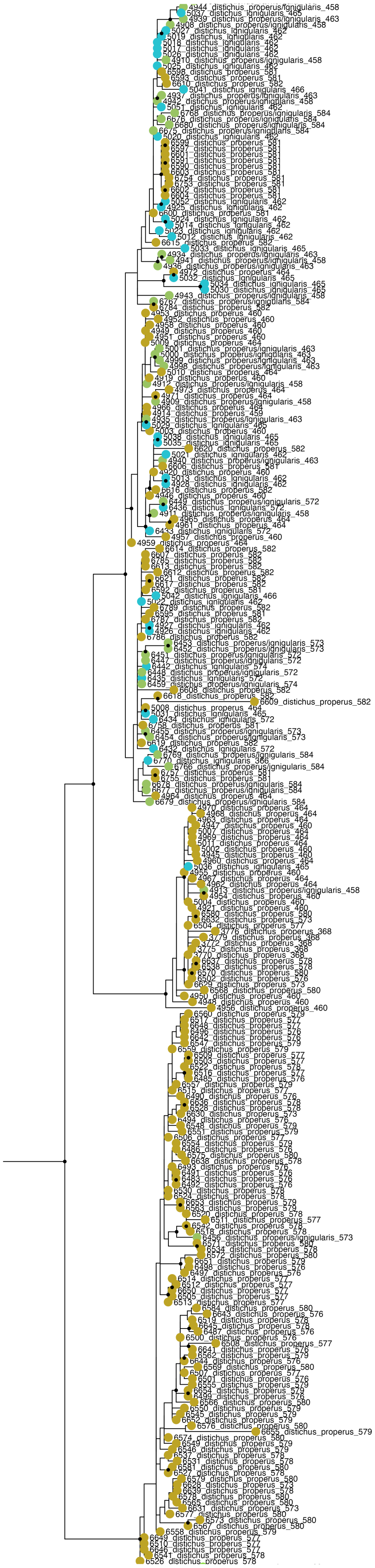

Figure S21. ***ND2* maximum likelihood phylogeny of *A. d. properus* and *A. d. ignigularis*.**

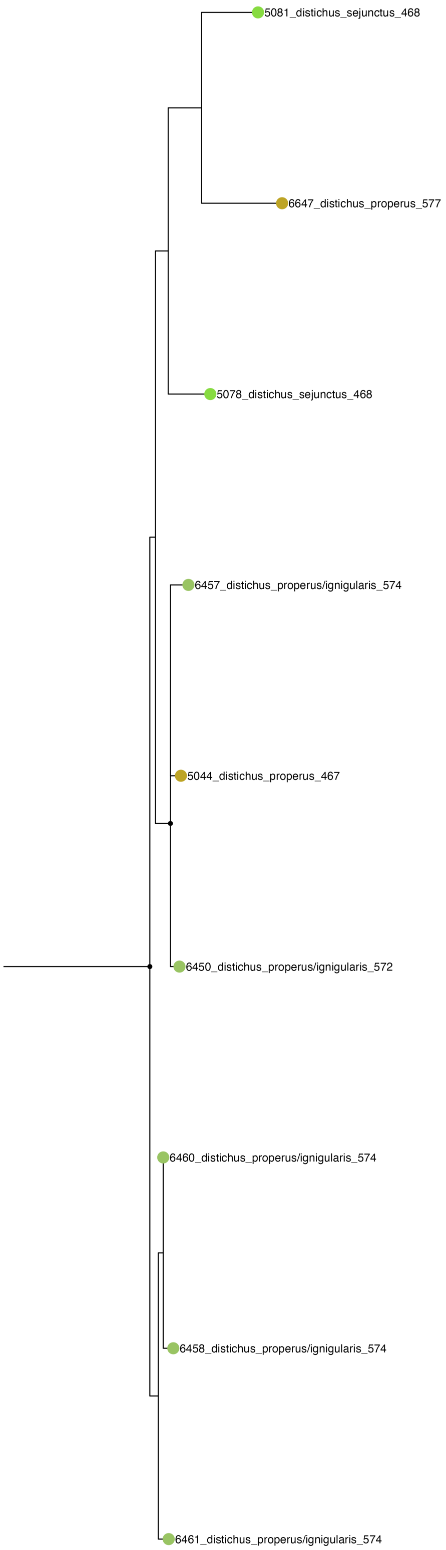

Figure S22. ***ND2* maximum likelihood phylogeny of *A. d. properus* and *A. d. sejunctus*.**

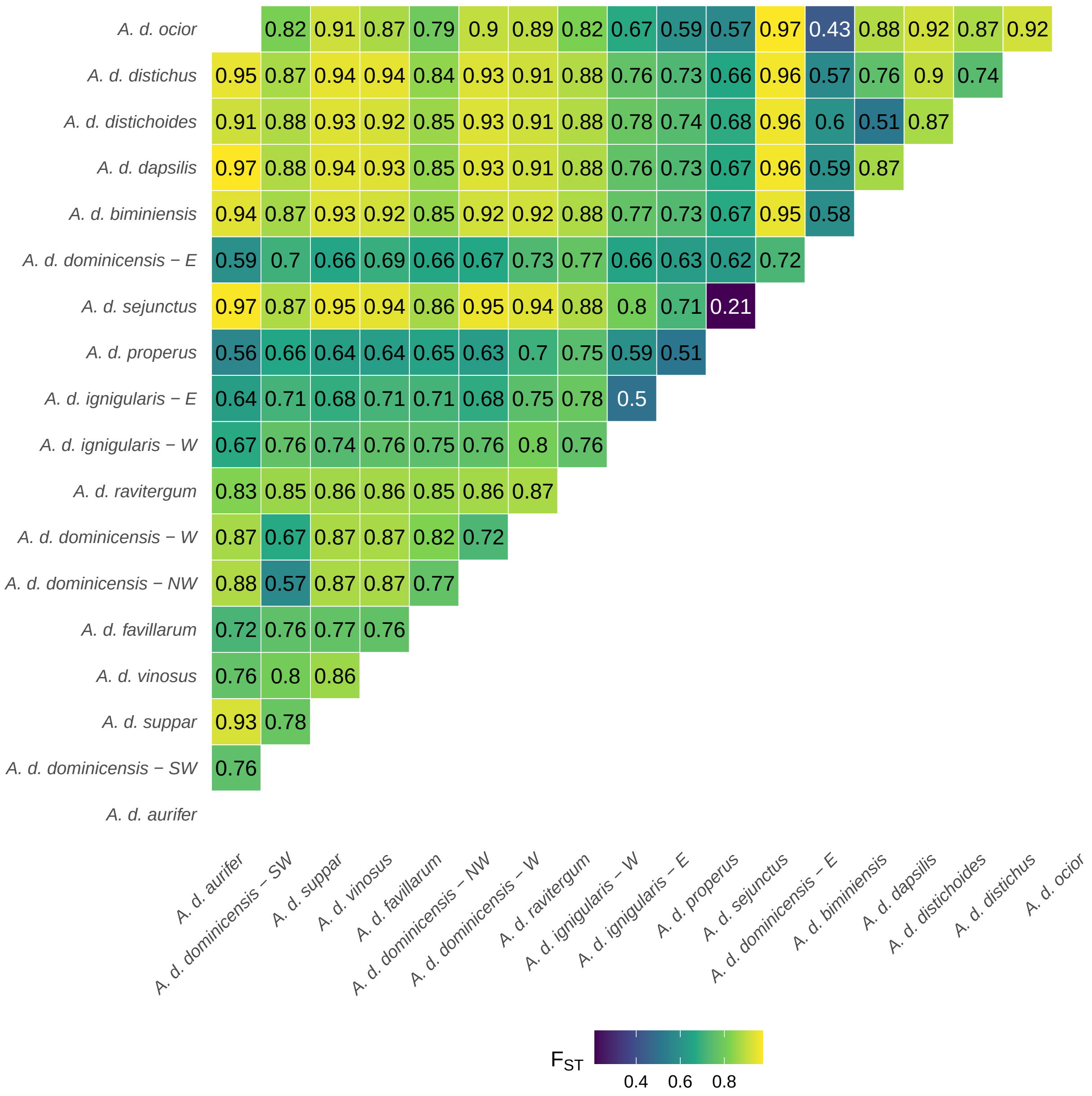

Figure S23. **Pairwise genetic differentiation ($F_{ST}$) of the Y chromosome within and among populations of *A. distichus*.**

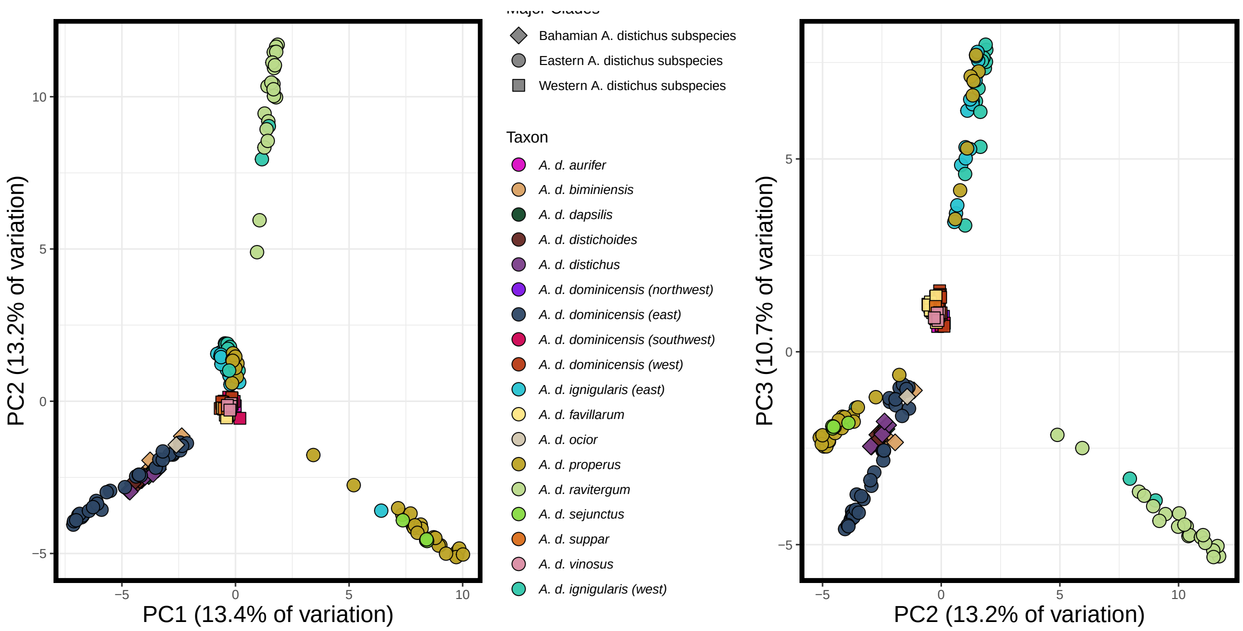

Figure S24. **Population structure patterns on the Y chromosome.**

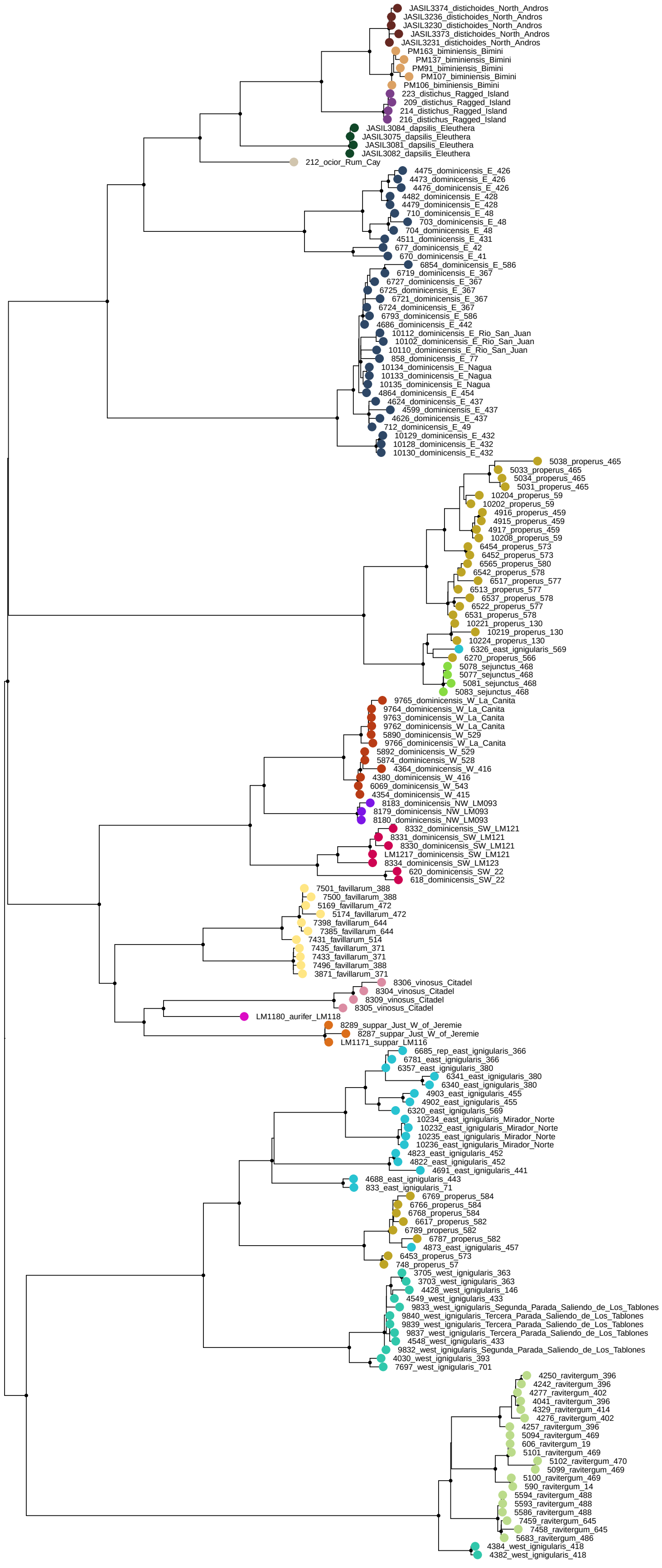

Figure S25. **Maximum likelihood phylogeny of concatenated Y chromosome ddRAD loci.**

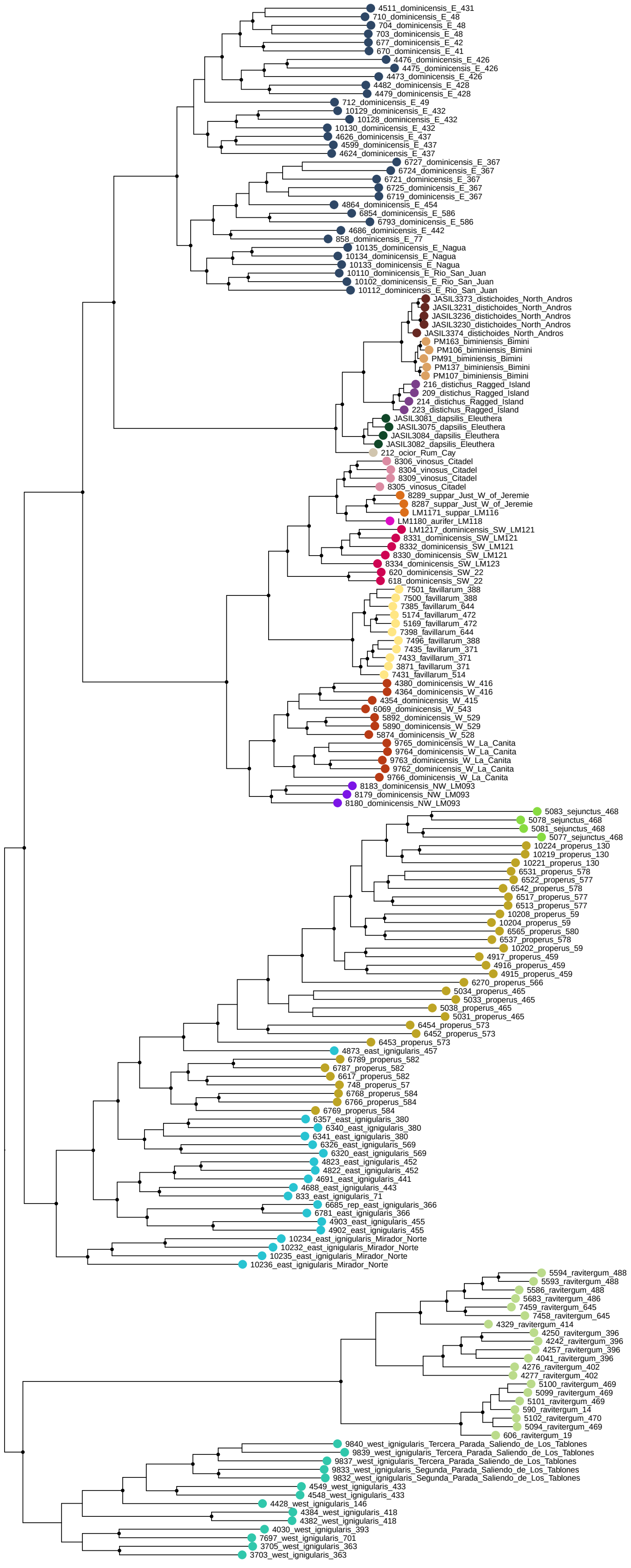

Figure S26. **Maximum likelihood phylogeny of concatenated autosomal ddRAD loci.**

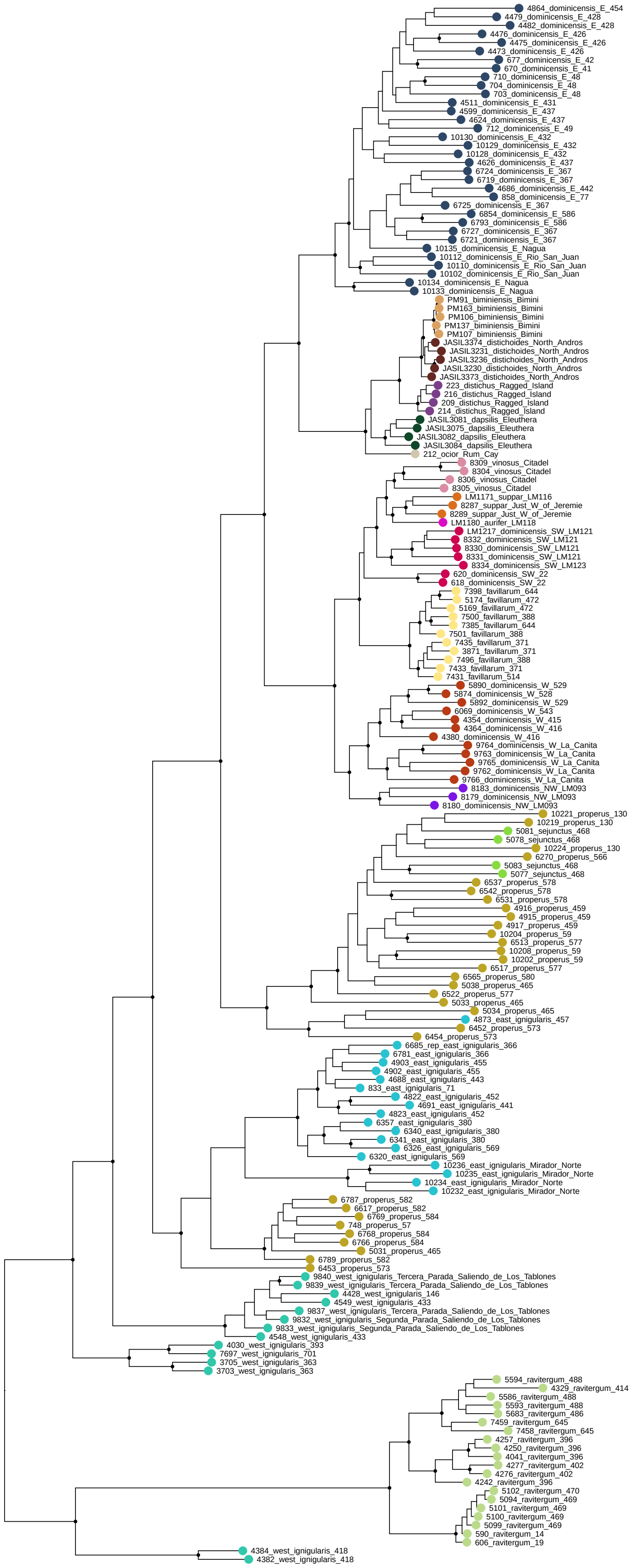

Figure S27. **Maximum likelihood of concatenated ddRAD loci on the ancestral X chromosome (X1).**

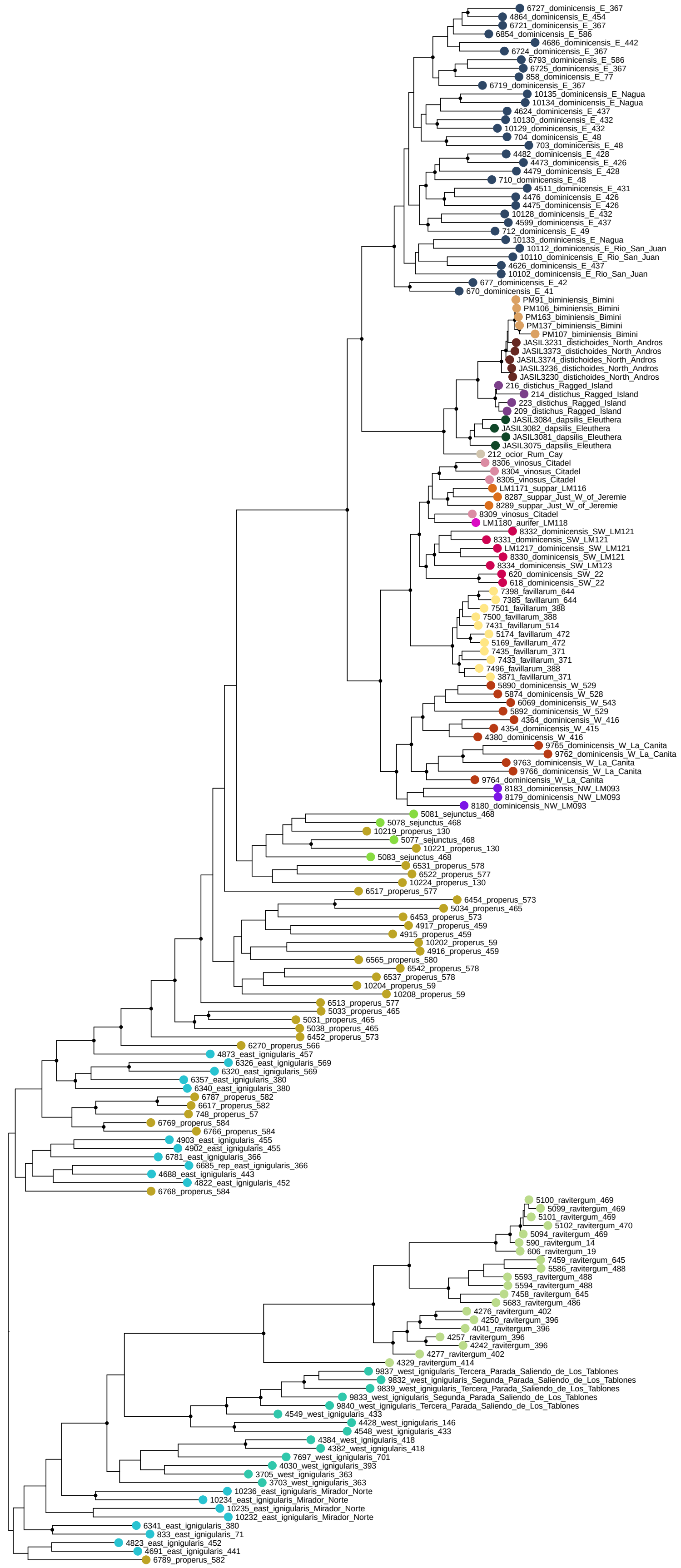

Figure S28. **Maximum likelihood of concatenated ddRAD loci on the neo-X chromosome (X2).**

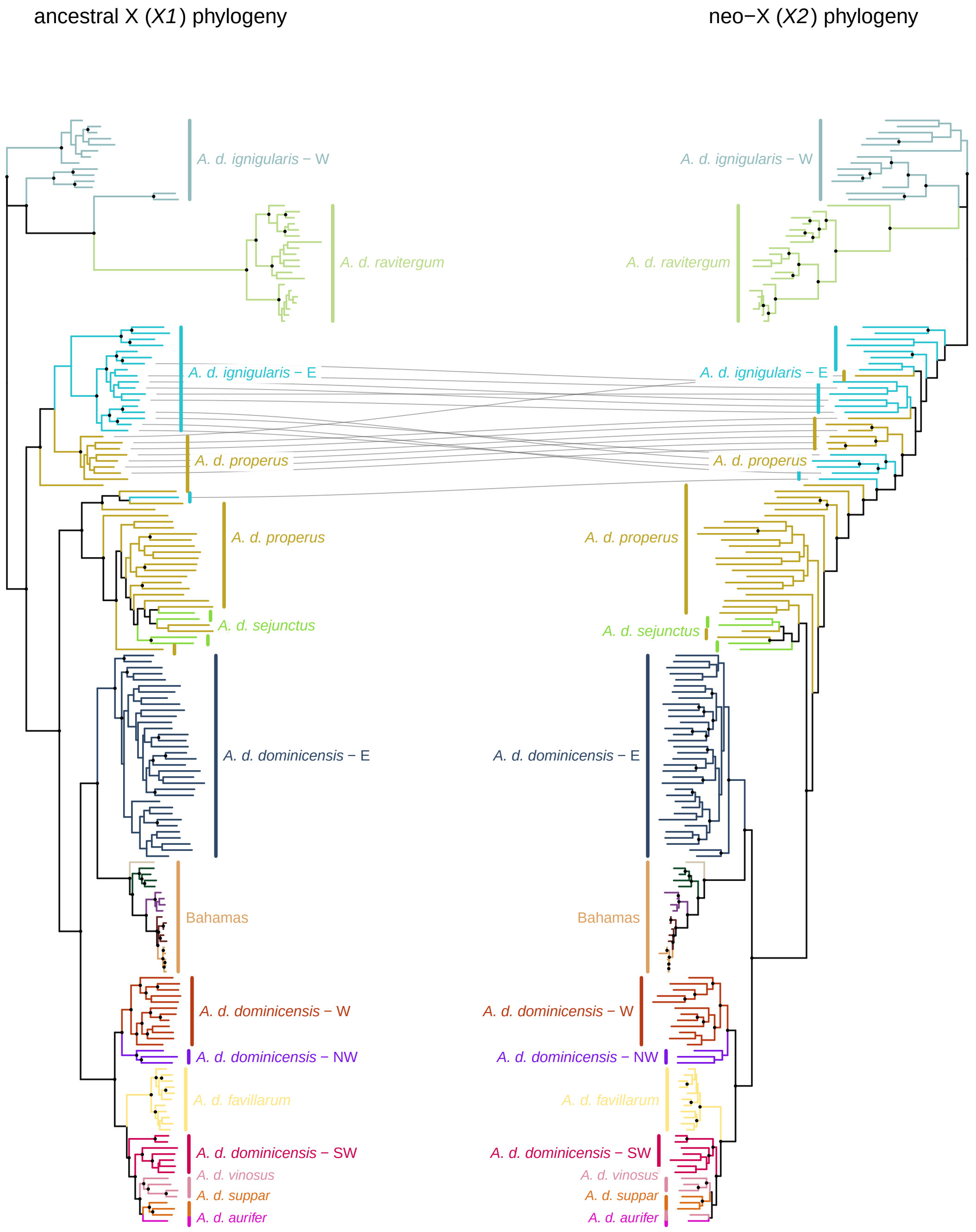

Figure S29. **X1 and X2 chromosome discordance.**

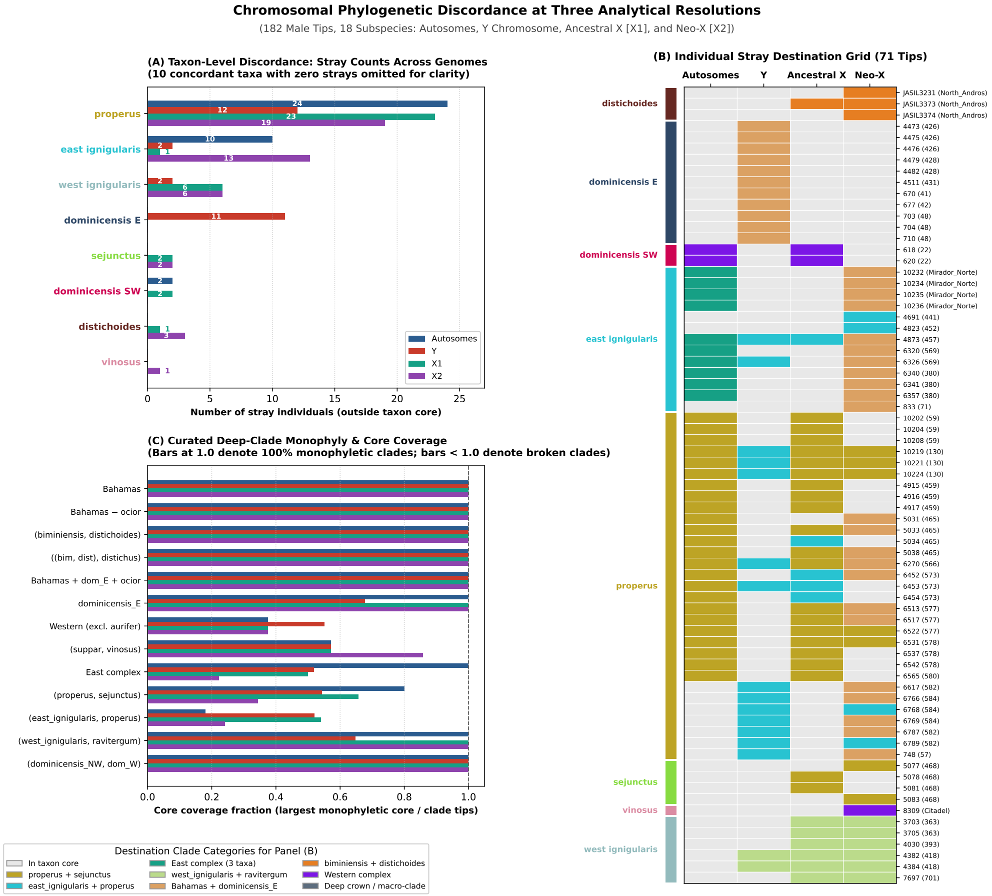

Figure S30. **Summary of sex chromosome and autosome discordance.**

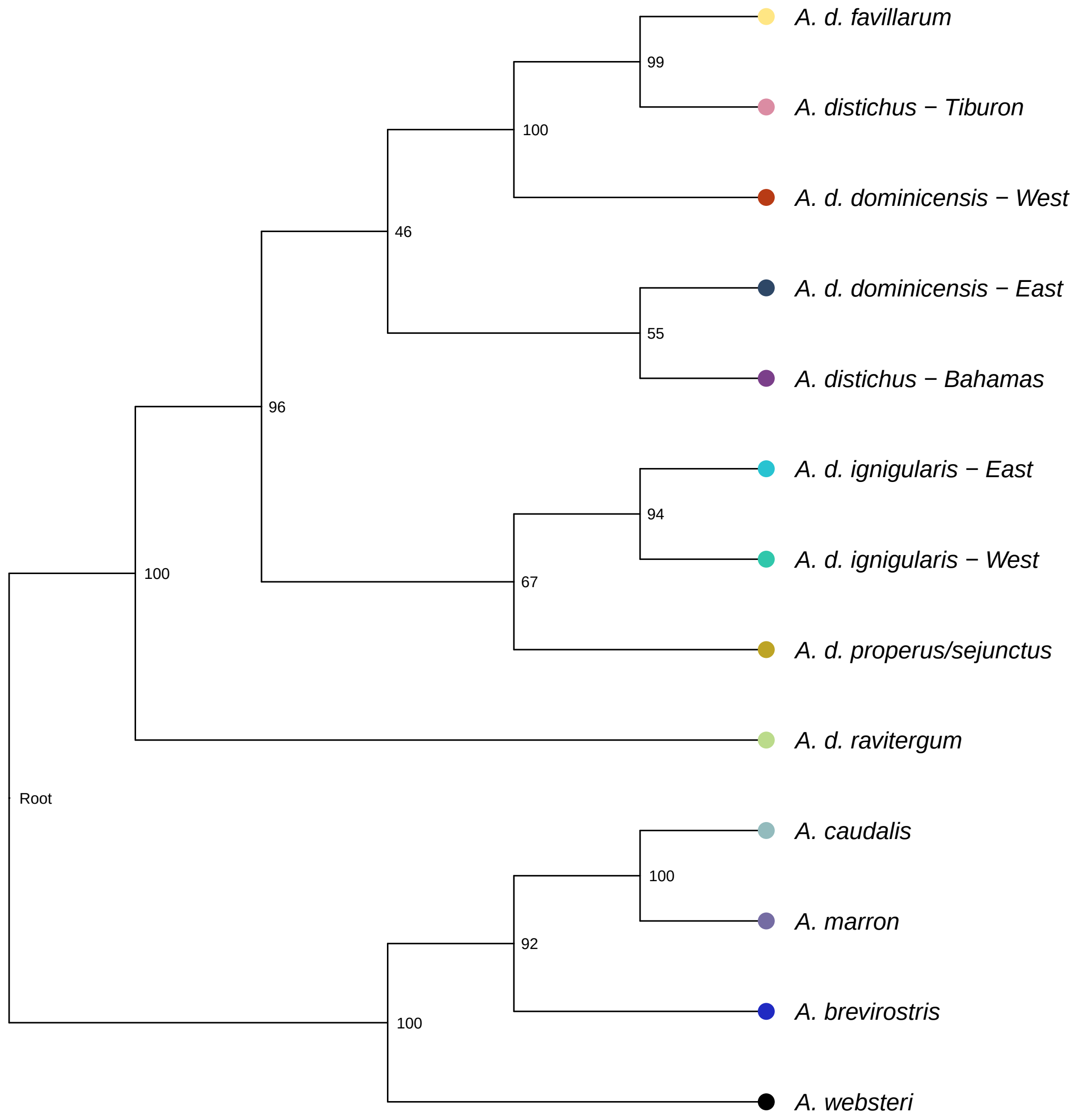

Figure S31. **SVDquartets species tree.**

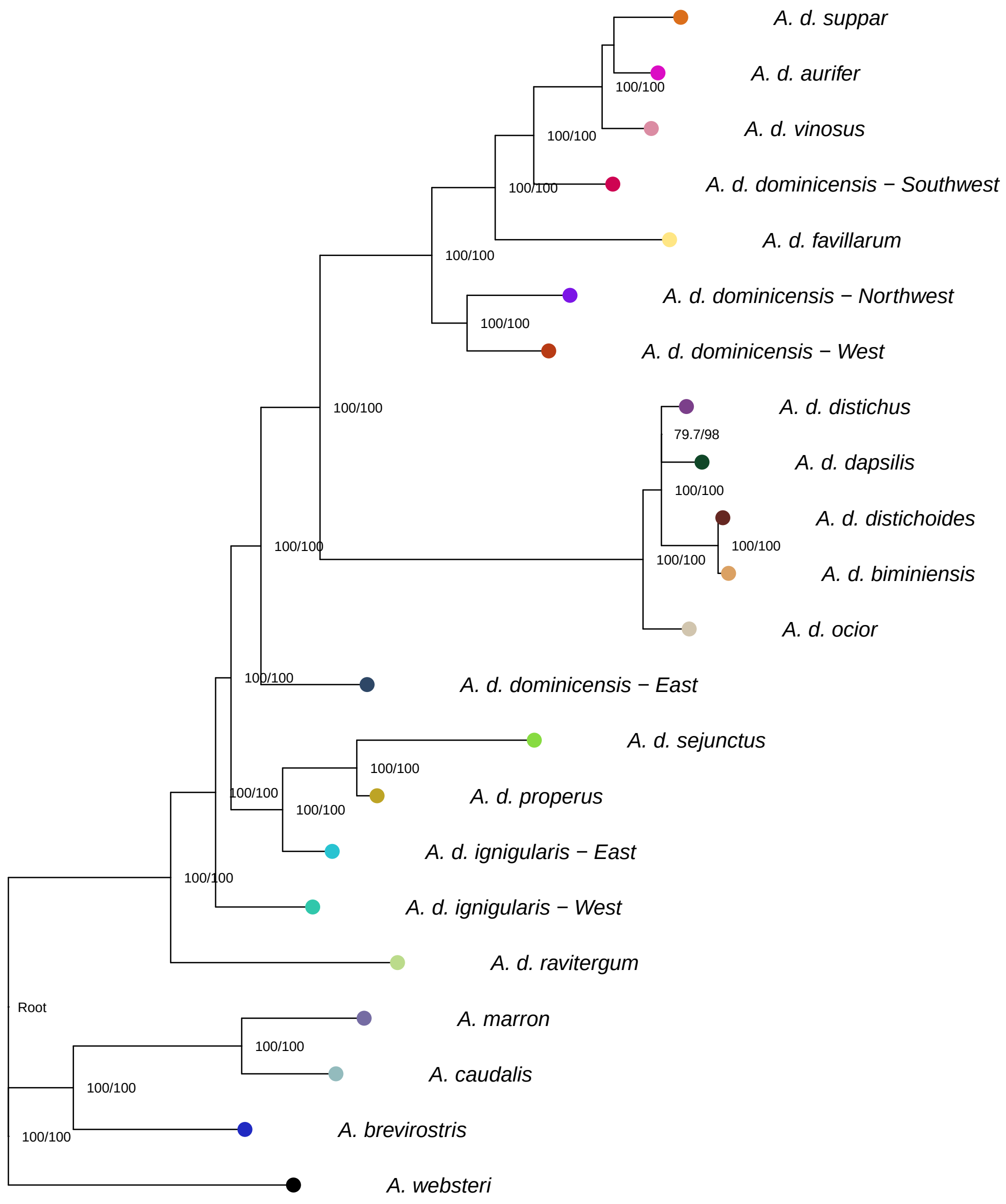

Figure S32. **Polymorphism-aware phylogenetic model (PoMo) species tree.**

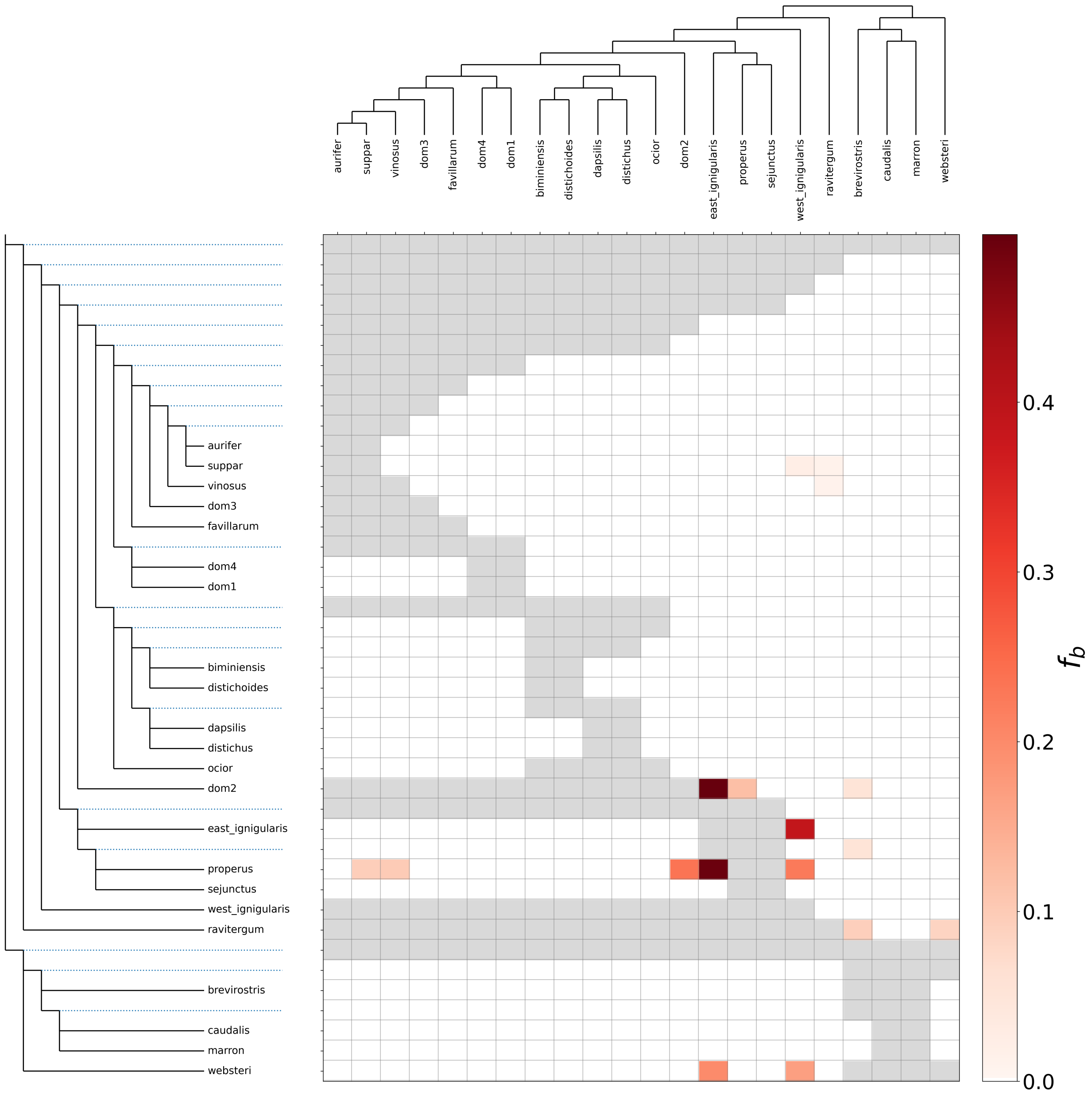

Figure S33. ***f*-branch statistic heatmap.**

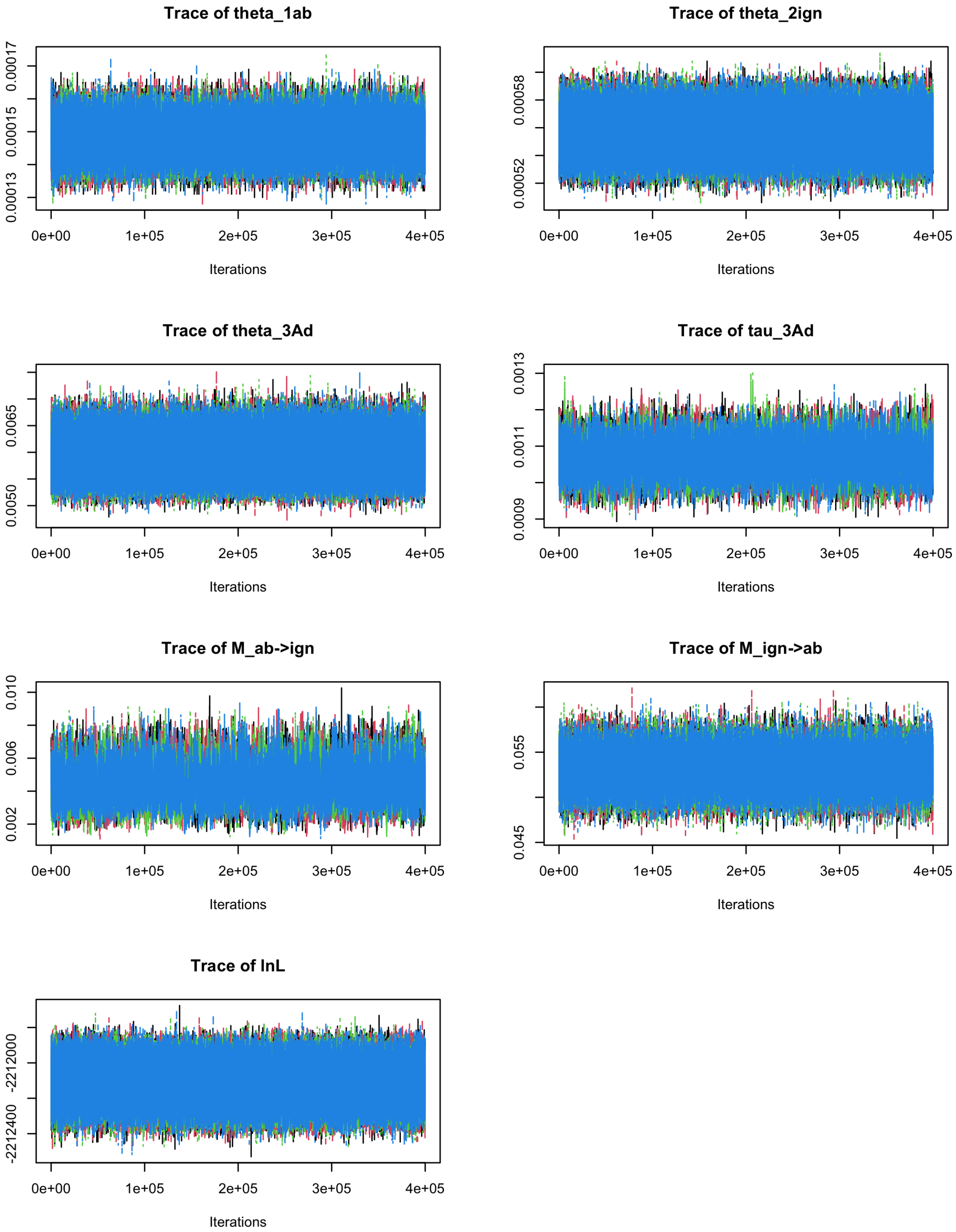

Figure S34. **Traceplots for MSC-M analysis between *A. d. ignigularis* and *A. b. deserticola*.**

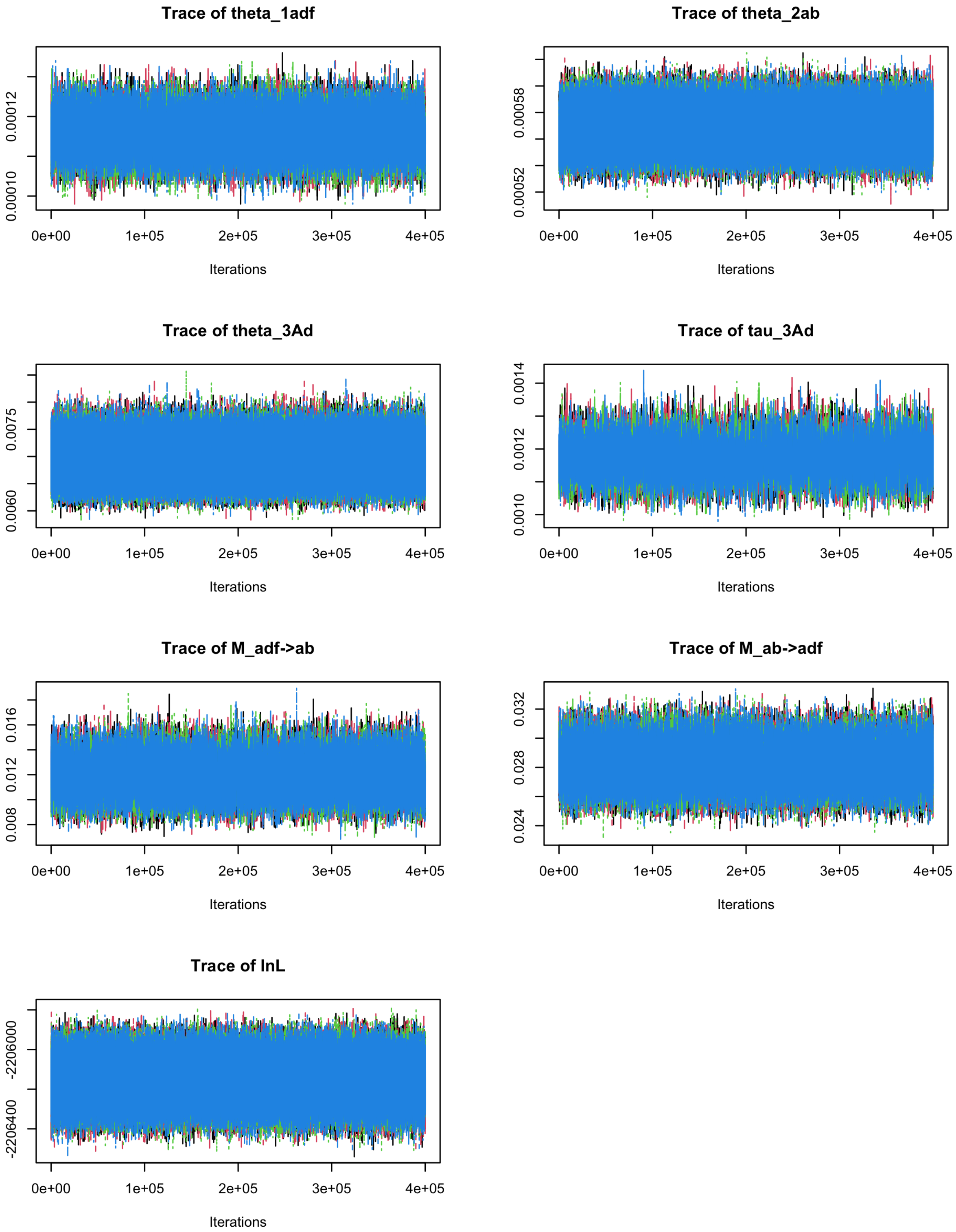

Figure S35. **Traceplots for MSC-M analysis between *A. d. favillarum* and *A. b. wetmorei*.**

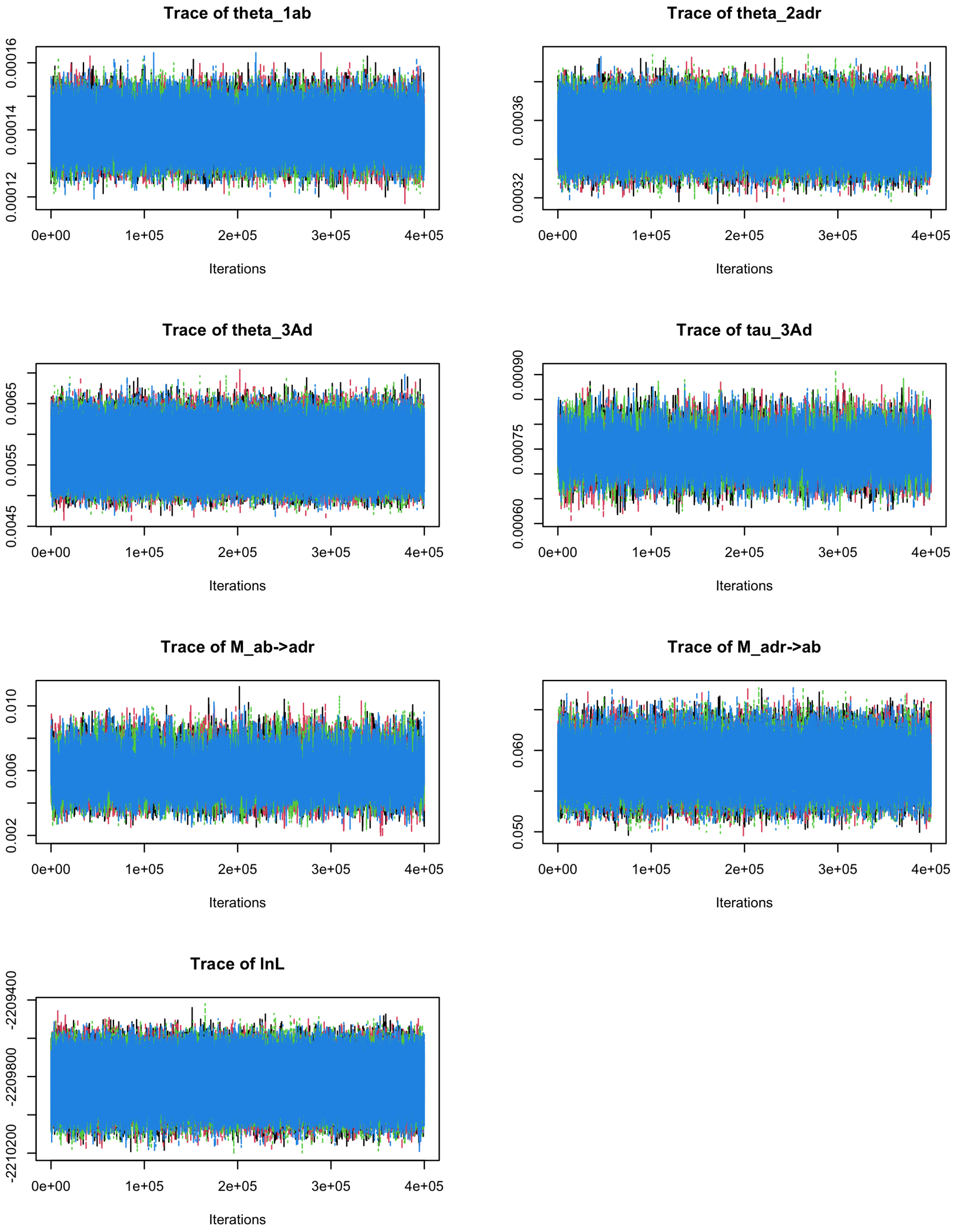

Figure S36. **Traceplots for MSC-M analysis between *A. d. ravitergum* and *A. b. deserticola*.**

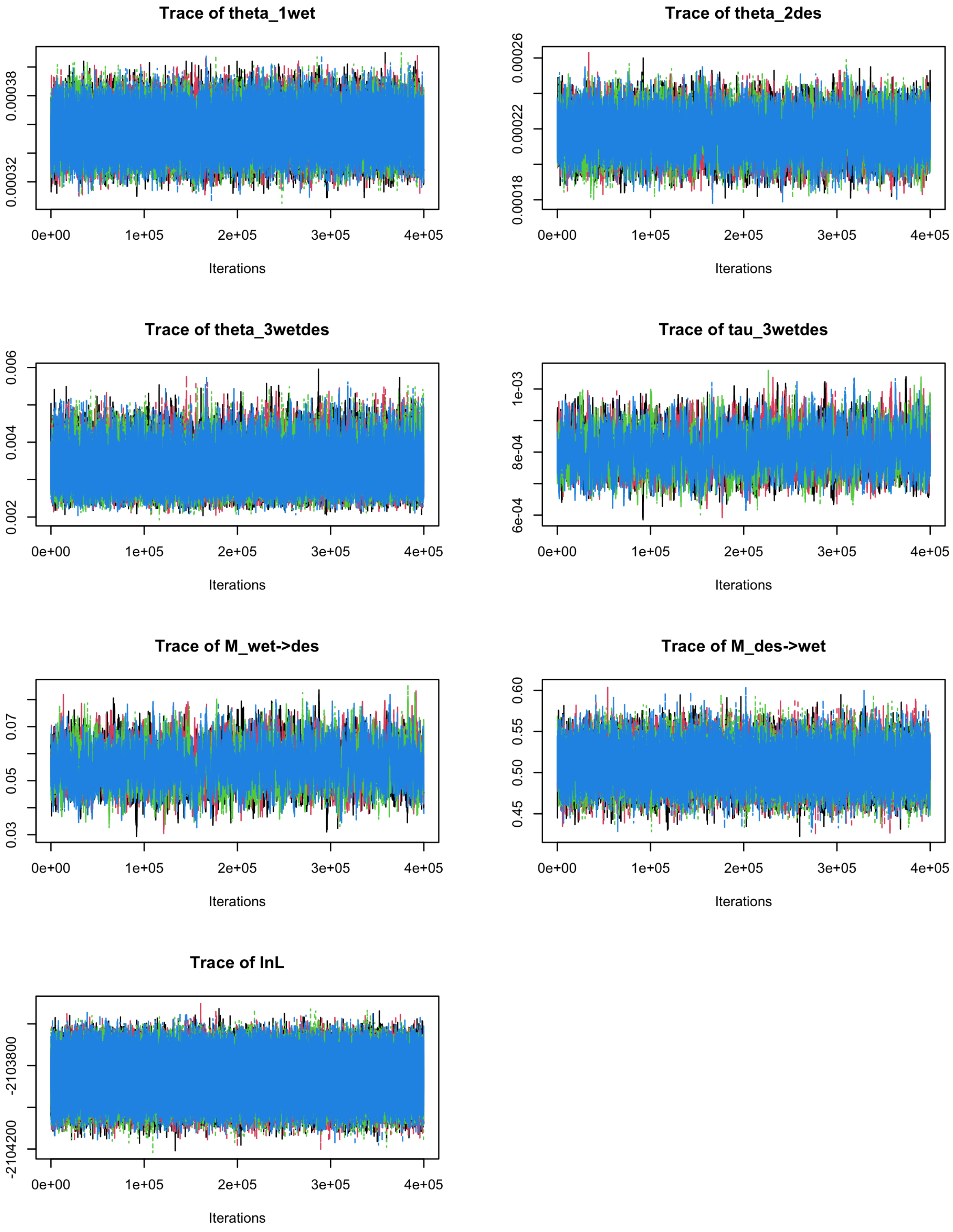

Figure S37. **Traceplots for MSC-M analysis between *A. b. deserticola* and *A. b. wetmorei*.**

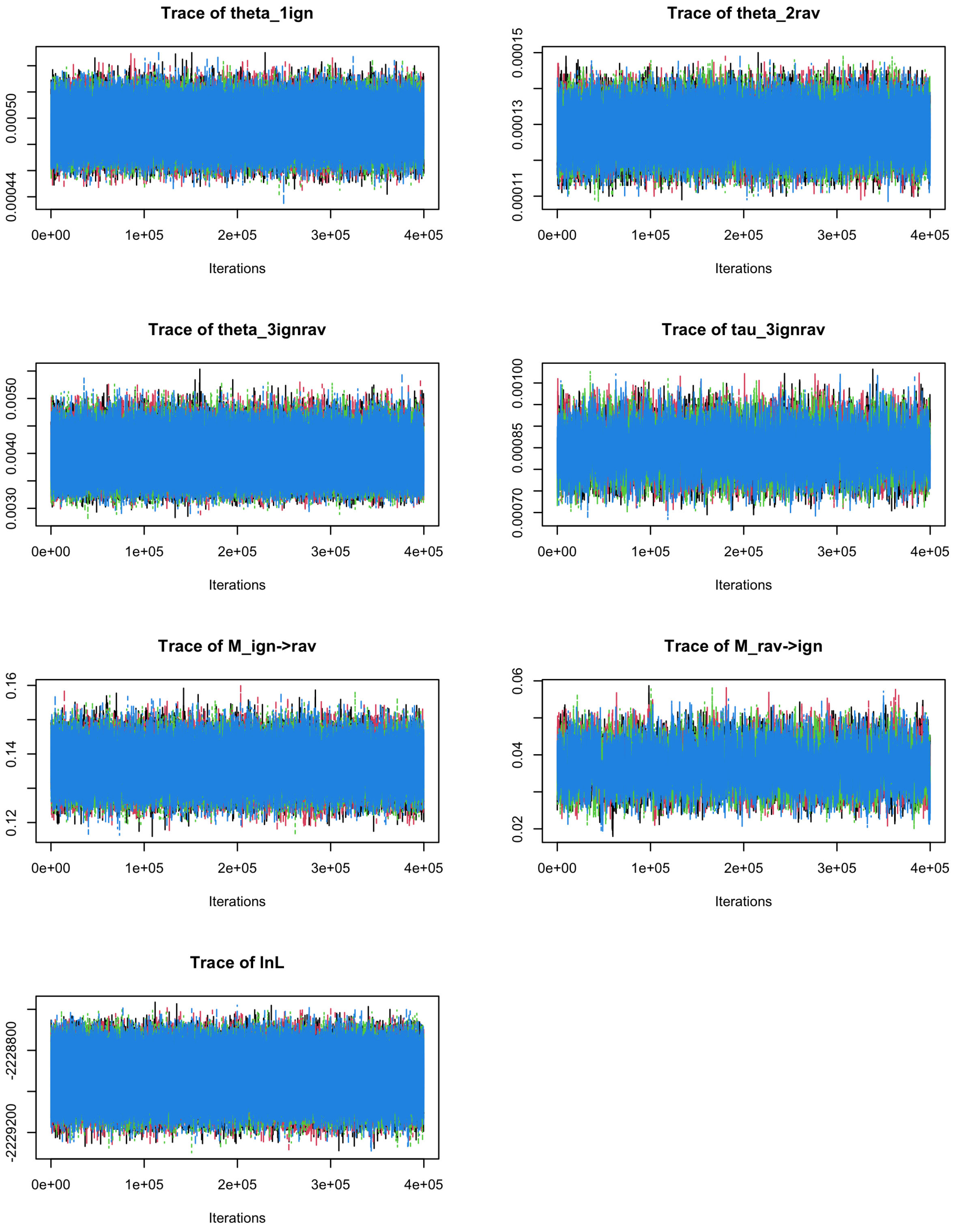

Figure S38. **Traceplots for MSC-M analysis between east *A. d. ravitergum* and west *A. d. ignigularis*.**

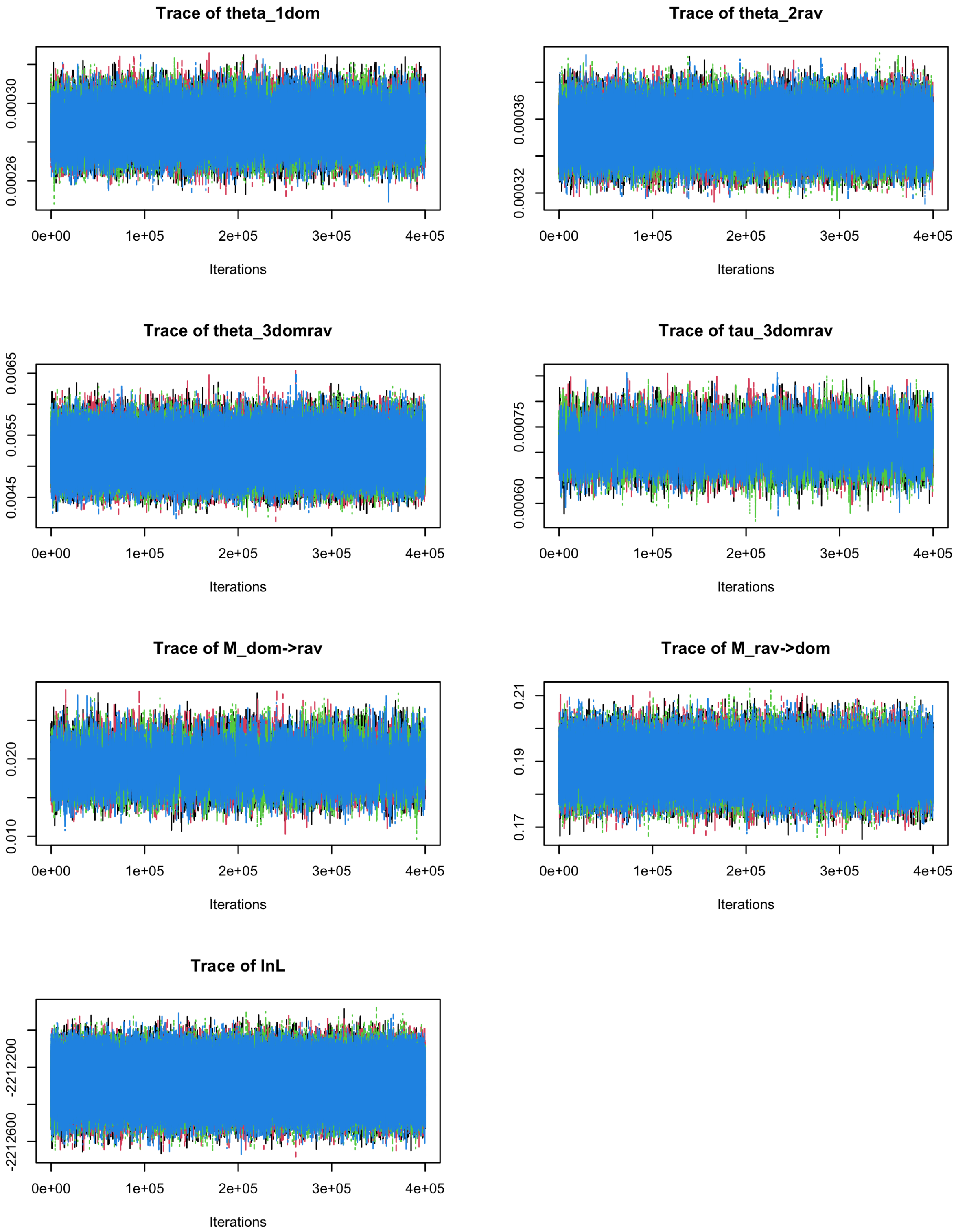

Figure S39. **Traceplots for MSC-M analysis between east *A. d. ravitergum* and west *A. d. dominicensis*.**

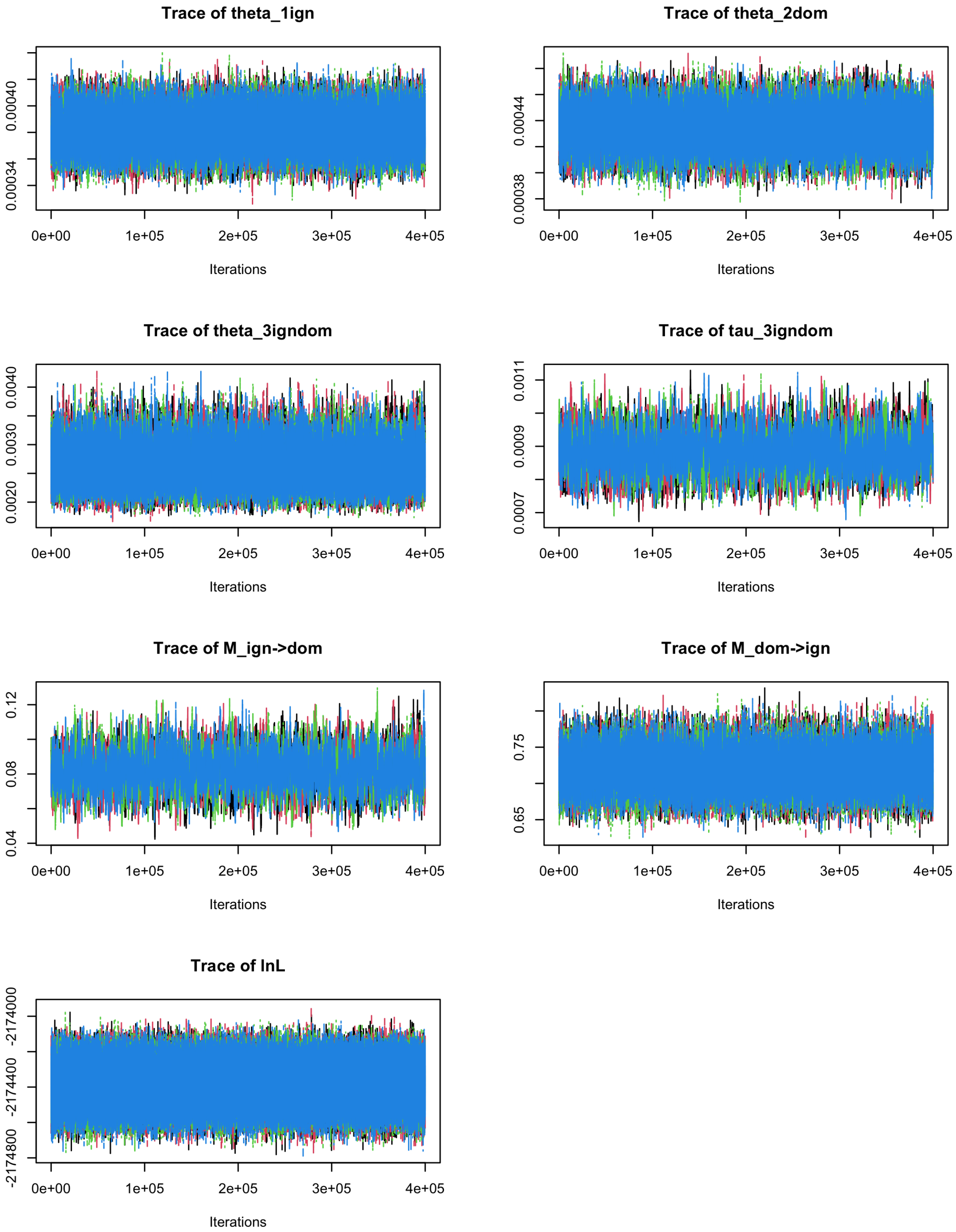

Figure S40. **Traceplots for MSC-M analysis between east *A. d. dominicensis* and *A. d. ignigularis* populations.**

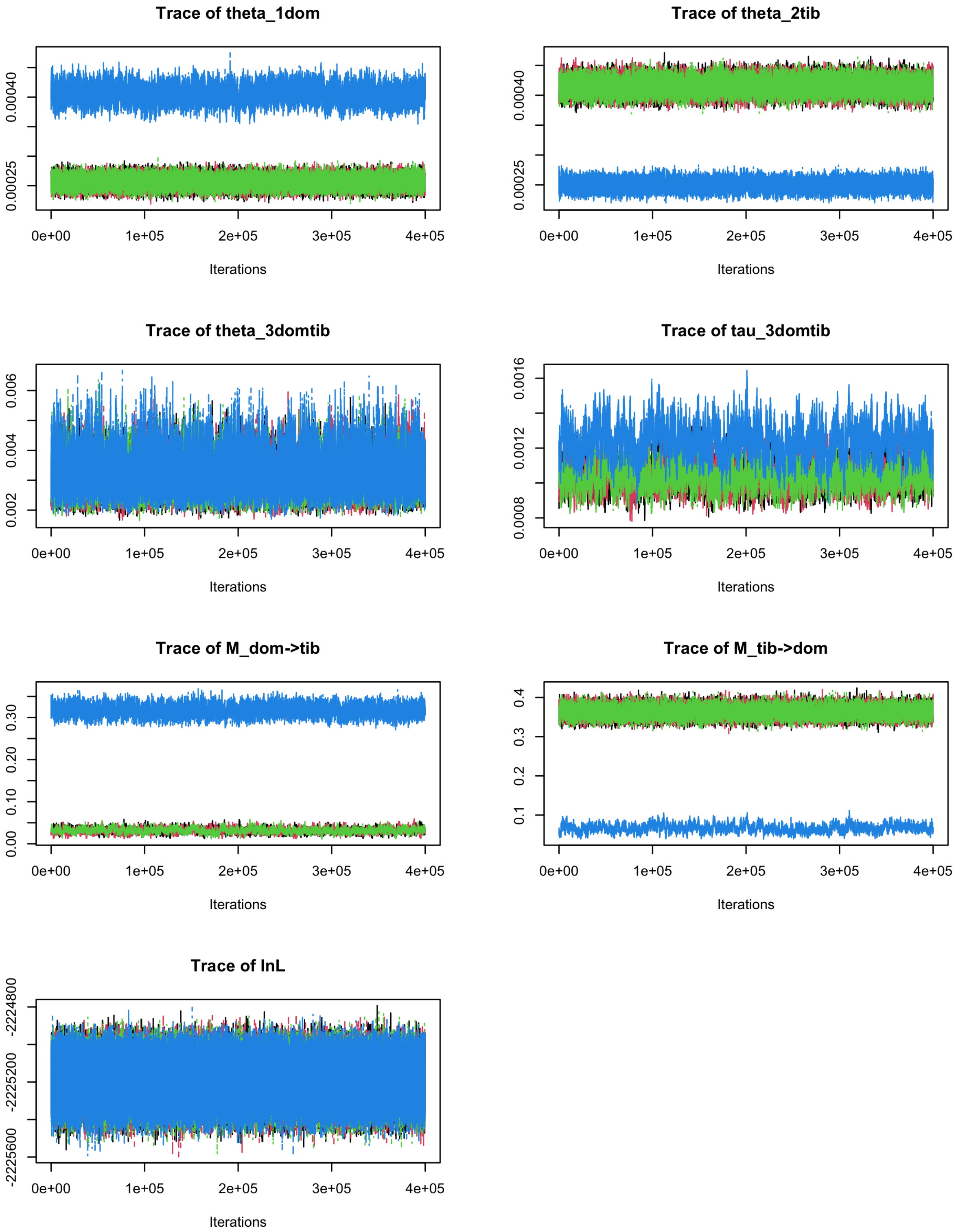

Figure S41. **Traceplots for MSC-M analysis between the Tiburón penninsula subspecies of *A. distichus* and west *A. d. dominicensis*.**

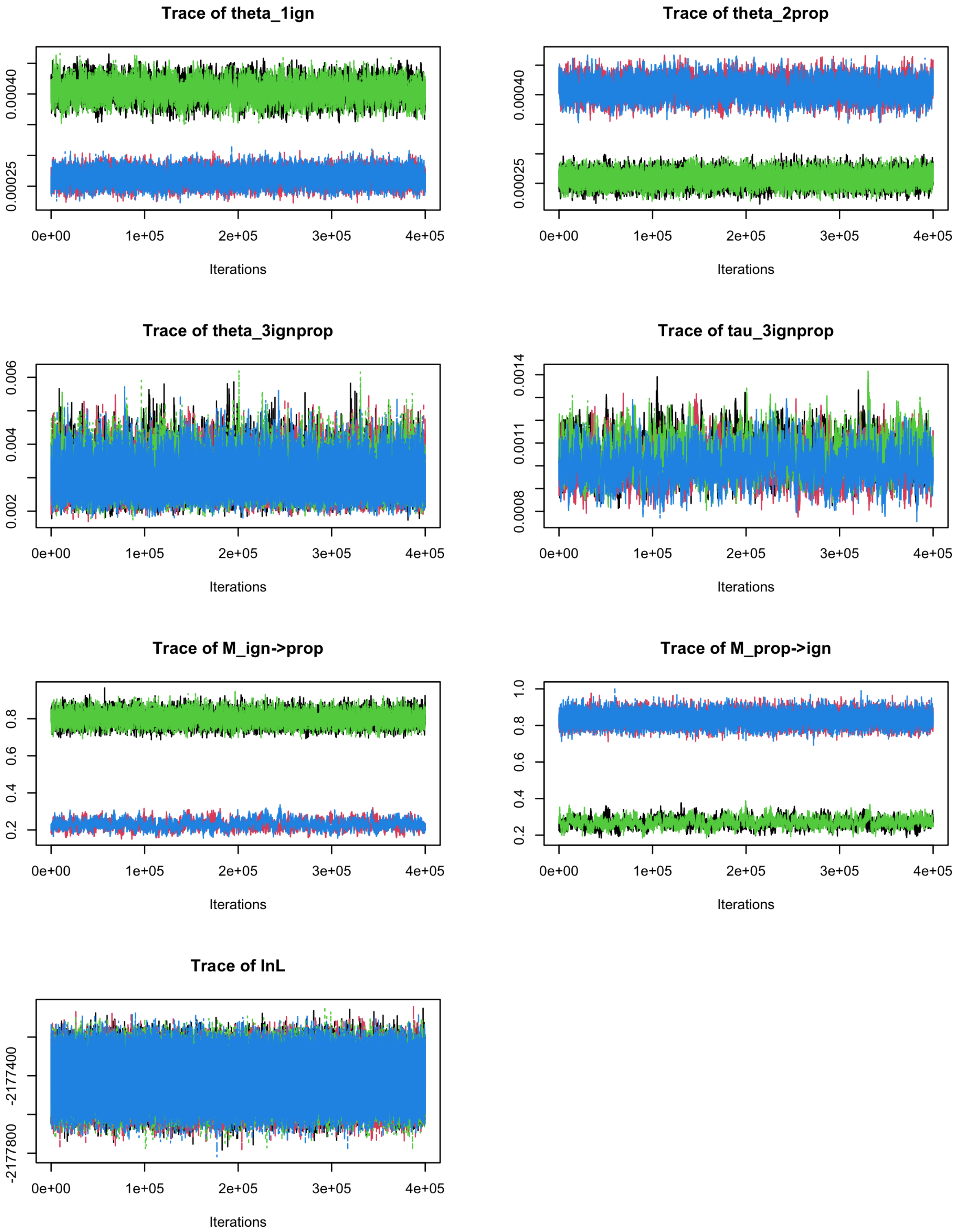

Figure S42. **Traceplots for MSC-M analysis between east *A. d. properus* and east *A. d. ignigularis*.**

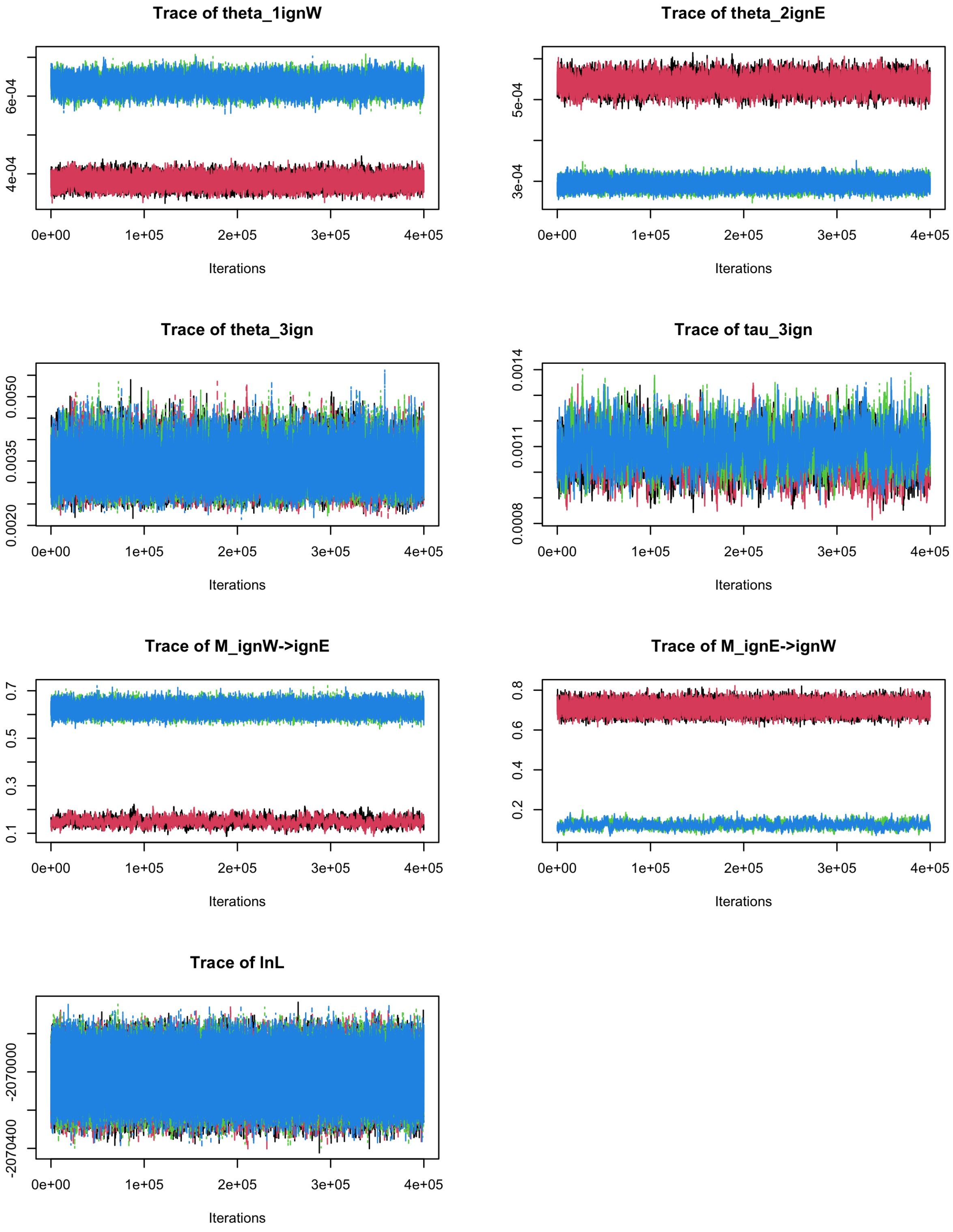

Figure S43. **Traceplots for MSC-M analysis between east and west *A. d. ignigularis* populations.**

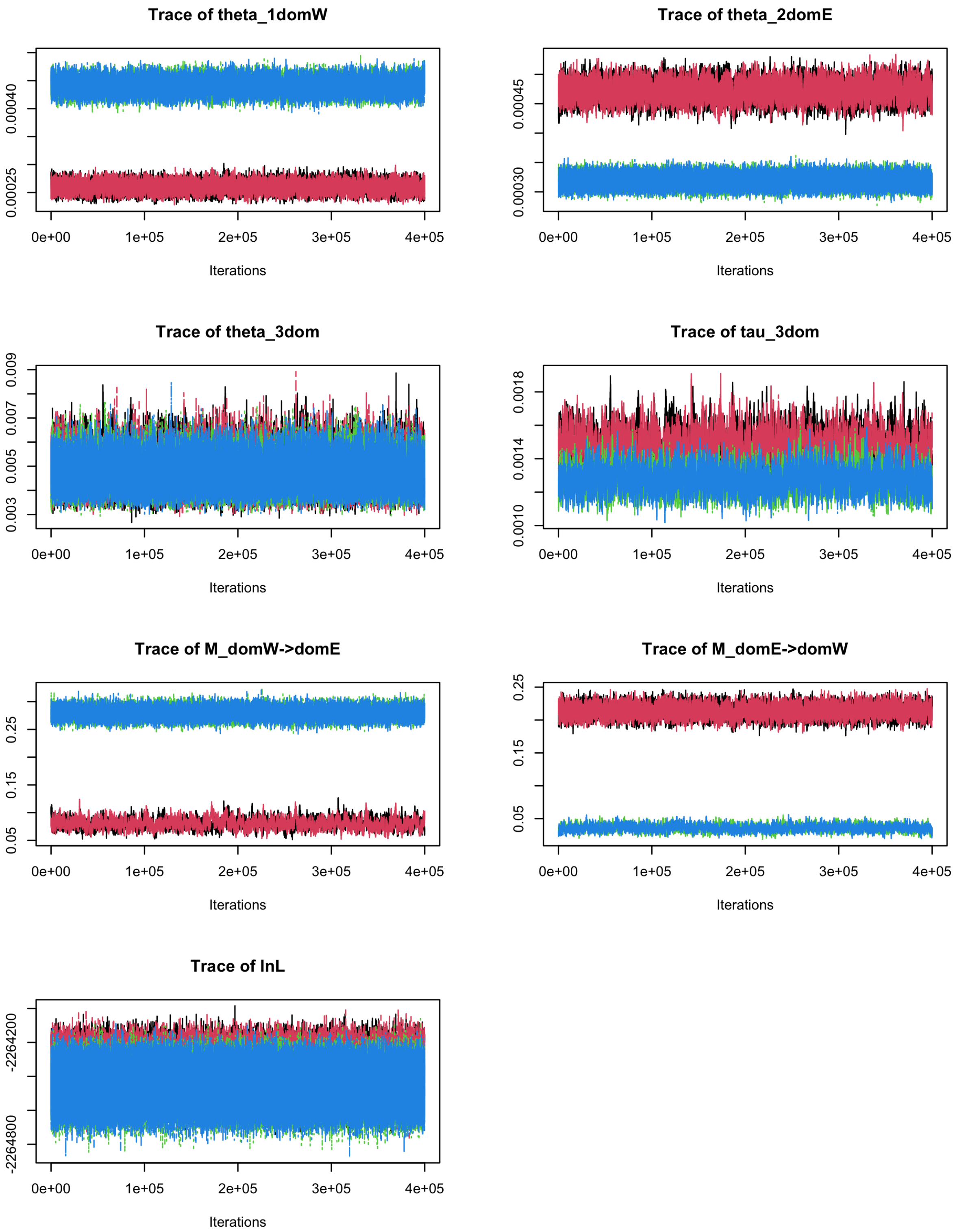

Figure S44. **Traceplots for MSC-M analysis between east and west *A. d. dominicensis* populations.**

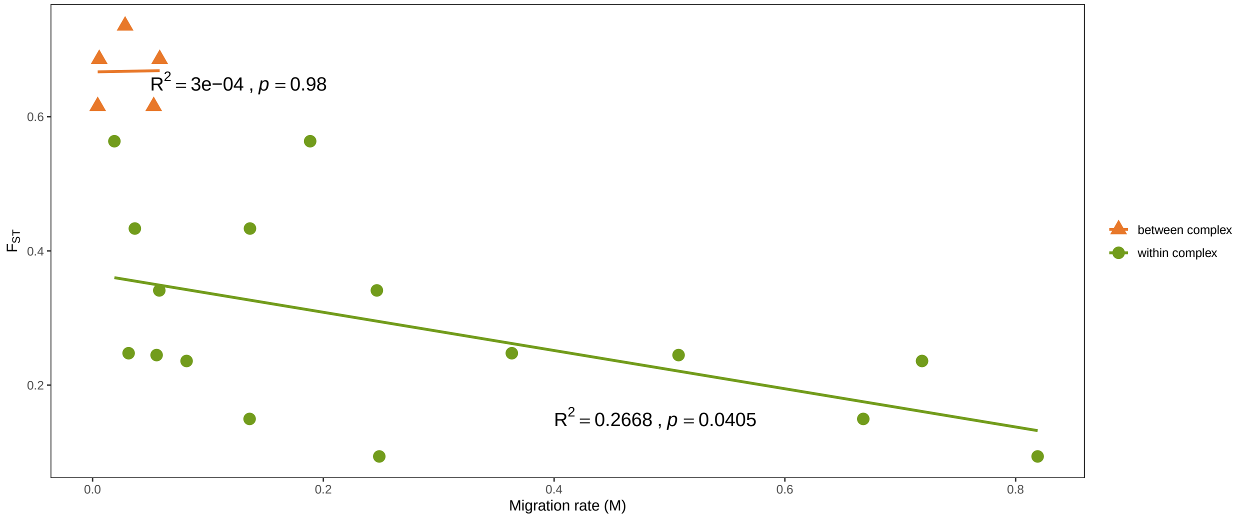

Figure S45. **Relationship between migration rate (*M*) and $F_{ST}$.**
